# Combinatorial single-cell profiling of all major chromatin types with MAbID

**DOI:** 10.1101/2023.01.18.524584

**Authors:** Silke J.A. Lochs, Robin H. van der Weide, Kim L. de Luca, Tessy Korthout, Ramada E. van Beek, Hiroshi Kimura, Jop Kind

## Abstract

Gene expression programs result from the collective activity of many regulatory factors. To obtain insight into the mechanisms that govern gene regulation, it is imperative to study their combined mode of action and interconnectivity. However, it has been challenging to simultaneously measure a combination of these factors within one sample. Here, we introduce MAbID, a method that combines genomic profiling of many histone modifications and chromatin-binding proteins in a single reaction. MAbID employs antibody-DNA conjugates to enable genomic barcoding of chromatin at sites of epitope occupancy. This barcoding strategy allows for the combined incubation of multiple antibodies in a single sample to reveal the genomic distributions of many epigenetic states simultaneously. We used MAbID to profile both active and inactive chromatin types in human cell lines and multiplexed measurements in the same sample without loss of data quality. Moreover, we obtained joint measurements of six epitopes covering all major chromatin types in single cells during mouse *in vitro* neural differentiation and captured associated changes in multifactorial chromatin states. Thus, MAbID holds the potential to gain unique insights into the interplay between gene regulatory mechanisms, especially in settings with limited sample material and in single cells.

---

Gene regulation involves the coordinated activity of many factors at different genomic scale levels. On a larger scale, chromosomes occupy distinct territories within the nuclear space^1,2^ that are further partitioned into compartments of similar chromatin states^3,4^. At a more local level, DNA methylation^5^, histone post-translational modifications (PTMs)^6^ and chromatin remodeling complexes^7^ synergistically modulate interactions between promoters and transcription factors, thereby regulating gene activity. The collective action of all these layers of regulation ultimately determines cellular identity and function. To gain a deeper understanding of the mechanisms governing gene expression, technologies capable of simultaneously measuring multiple components of gene regulation are required.

In recent years, there has been a vast advancement of multi-omic strategies that enable the profiling of several modalities in single cells. Most prominently among these are methods linking transcriptional heterogeneity to variations in DNA methylation^8–10^, chromatin accessibility^9,11,12^, protein-DNA binding^13,14^, nuclear architecture^13–15^ and histone PTMs^16–20^. However, these techniques can generally only profile one of these modalities in combination with transcription, while methods that can simultaneously measure multiple gene-regulatory components are limited.

Several promising approaches have recently been developed in which up to three histone PTMs can be profiled in the same cell^21–26^. Such methods hold great potential for dissecting the interdependencies and sequence of events underlying the mechanistic basis of gene regulation. These recent multifactorial methodologies almost exclusively rely on antibody detection followed by Tn5-mediated tagmentation, which is commonly used in state-of-the-art genomic profiling techniques^27^. The advantage of this approach is that Tn5 is very efficient and yields specific data at a high resolution. However, its propensity towards integrating into open chromatin regions^28,29^ may introduce accessibility biases and limits measuring modalities that reside in constitutive heterochromatin.

Here, we present Multiplexing Antibodies by barcode Identification (MAbID), a method that is independent of Tn5-mediated tagmentation and instead employs standard restriction-digestion and ligation steps. Genomic profiling with MAbID generates a signal of comparable distribution and resolution to that obtained by state-of-the-art methods like Chromatin Immunoprecipitation sequencing (ChIP-seq). Moreover, the approach enables measurements of epitopes across all major chromatin types, including active chromatin as well as facultative and constitutive heterochromatin. We employed secondary or primary antibody-DNA conjugates to generate low-input and single-cell readouts for up to six epitopes simultaneously. The quality and specificity of the data are independent of the number of antibodies multiplexed in one sample, which shows the potential of MAbID to significantly increase multiplexing beyond the six epitopes presented here. We demonstrated that this method is able to capture single-cell changes in chromatin states associated with cellular transitions in mouse embryonic stem cells differentiating towards the neural lineage. We anticipate that MAbID provides a new technology to obtain further insight into the regulation of gene expression in dynamic and complex biological settings

## Results

### MAbID enables genomic profiling of a broad spectrum of chromatin states

To multiplex measurements of several chromatin states within the same sample, we devised a strategy to uniquely barcode antibodies and map their epitope-positions on the chromatin via specific restriction-ligation steps. For this purpose, each antibody is first covalently linked to a double-stranded DNA-adapter (antibody-adapter) using a basic two-step Cu^2+^-free click-chemistry approach (SPAAC)^30–32^ (Extended Data Fig. 1a-b). These antibody-DNA conjugates are then employed in the MAbID protocol (Fig. 1a), which starts with 1) harvesting of ~250,000 cells, nuclei isolation and mild fixation, 2) incubation with primary antibodies followed by incubation with uniquely barcoded secondary antibody-DNA conjugates, 3) Fluorescent-Activated Cell Sorting (FACS) into tubes or 384-well plates, 4) digestion of the genome with the MseI restriction enzyme, which recognizes TTAA sequence motifs, 5) dephosphorylation of the digested genome to prevent self-ligation of genomic fragments, 6) digestion of the antibody-adapter with the NdeI restriction enzyme, which leaves a MseI-compatible overhang with a 5’ phosphate and 7) ligation of the antibody-adapter into the digested genome using the matching overhangs. The position of the antibody-adapter within the genome hereby becomes a proxy for the localization of the epitope of interest. The protocol continues with 8) lysis and protein degradation followed by 9) digestion with the NotI restriction enzyme to enable subsequent ligation of a sample-adapter. This includes a T7 RNA polymerase promoter sequence, an Illumina P5-sequence and a unique-molecular identifier (UMI) interspersed with a sample-barcode (Extended Data Fig. 1b). The sample-adapter enables pooling multiple samples for linear amplification by *in vitro* transcription and subsequent Illumina library preparation as previously described^13^.

**Fig. 1.**
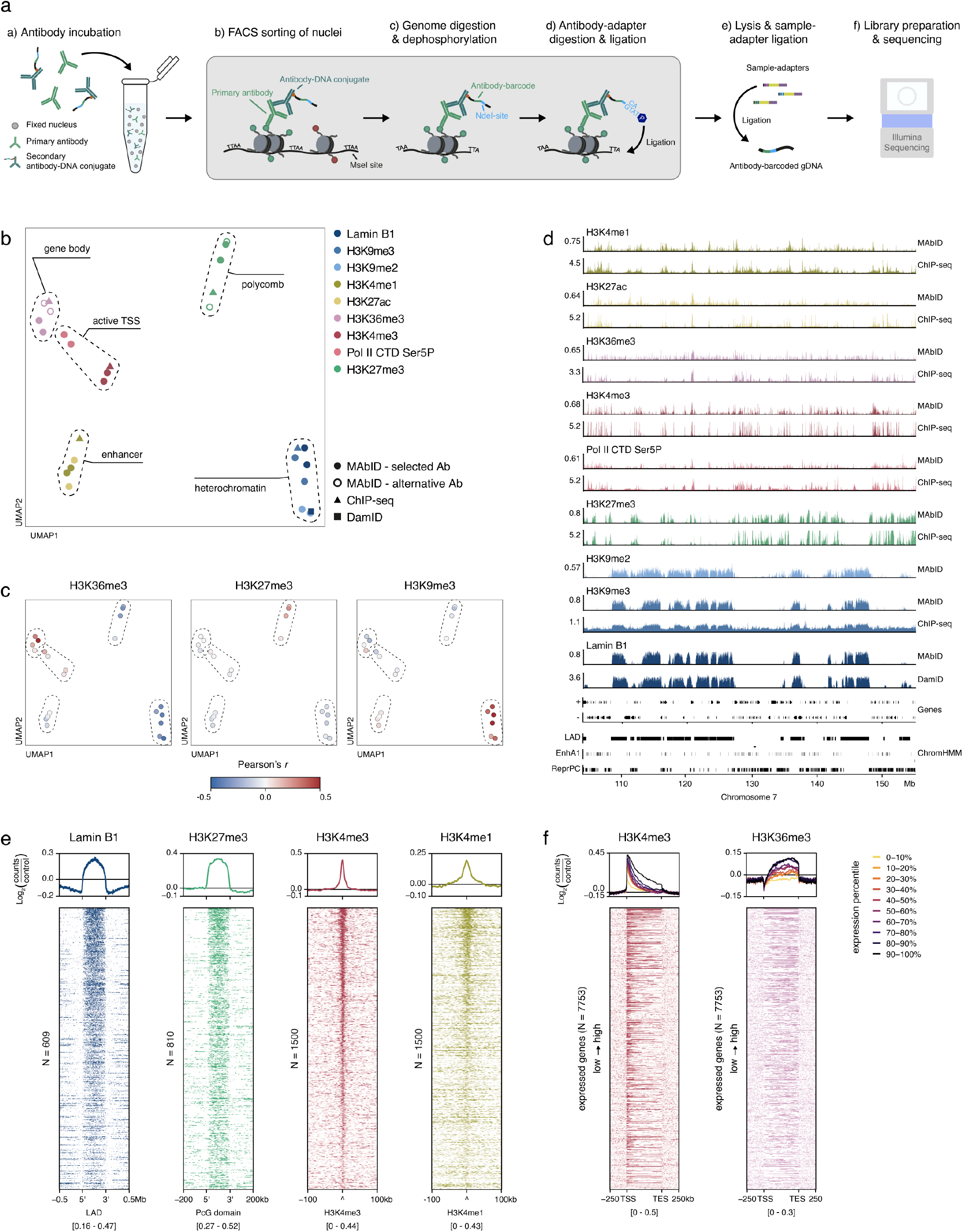
Genomic profiling of a broad range of epigenetic markers with MAbID. **a)** Schematic representation of the MAbID procedure. The protocol starts with a) incubation of nuclei with primary antibodies and secondary antibody-DNA conjugates, followed by b) FACS sorting and c) digestion of the genome and dephosphorylation of digested ends. Next, d) the antibody-adapter, containing an antibody-specific barcode, is digested and ligated into the digested genome at the location of the epitope of interest. After ligation, e) nuclei are lysed and sample-adapters are ligated to the antibody-barcoded genomic DNA in order to multiplex samples, followed by f) library preparation and next-generation sequencing. **b)** UMAP embedding of MAbID replicates together with ChIP-seq and DamID samples. Colouring is based on the epitope of interest and chromatin types are encircled. One reference dataset is included per chromatin type (if publicly available). Selected Ab indicates the primary antibody used in the next panels, alternative Ab represents a different primary antibody against the same epitope. **c)** UMAP as in (b), coloured by correlation with ChIP-seq samples of H3K36me3, H3K27me3 and H3K9me3. The colour scale represents the Pearson’s *r* correlation coefficient of MAbID samples with the indicated ChIP-seq sample. **d)** Genome browser tracks of MAbID with ChIP-seq or DamID samples. Genes (+, forward; -, reverse) and ENCODE/ChromHMM domain calls (LAD, Lamina-associated domain; EnhA1, Active enhancer 1; ReprPC, Repressed PolyComb) are indicated. Values on the y-axis reflect positive log_2_(counts/control) values for MAbID and DamID and fold change (IP/input) for ChIP-seq. **e)** MAbID signal enrichment of Lamin B1 around LAD regions (ENCODE, -/+ 0.5Mb), H3K27me3 around Polycomb-group domains (ChromHMM, -/+ 200kb) as well as H3K4me3 and H3K4me1 around their respective ChIP-seq peak calls (-/+ 100kb). Top line plot shows the average enrichment of signal, bottom heatmap shows signal per genomic region (sorted on MAbID signal). The number (N) of genomic regions included per heatmap is indicated. The heatmap data range is indicated underneath. **f)** MAbID signal enrichment of H3K4me3 and H3K36me3 around active genes (-/+ 250kb). Genes were stratified on expression level and categorized in percentiles, line plots indicate the average signal enrichment per percentile group. The heatmap below shows the signal per set of genes based on expression percentiles, ordered from high to low, including 7553 genes (N) in total. The heatmap data range is indicated underneath.

We first benchmarked the approach with individual measurements of a diverse set of epitopes, including several active and repressive histone PTMs, as well as RNA Polymerase II and the Lamin B1 protein. For this purpose, we performed MAbID with different primary antibodies (Supplementary Table 1) in populations of 1000 sorted K562 nuclei. We initially employed secondary antibody-DNA conjugates to more accurately compare the quality of multiple genomic profiles in parallel. These experiments were performed in biological replicates using one primary antibody against an epitope of interest per sample, which was targeted by the secondary antibody-DNA conjugate in a subsequent incubation. In parallel, a control sample was generated in which the primary antibody was omitted during the first incubation step. This sample serves as a non-specific binding control and is used as an input (mock IP) dataset to normalize the MAbID data (Extended Data Fig. 1c). On average 78.9% of the reads contained the correct sequence structure consisting of the antibody- and sample barcodes and 97.7% of these are located at the expected TTAA sequence motif (Extended Data Fig. 1d).

Visualization of the normalized data by uniform manifold approximation and projection (UMAP) shows good concordance between biological replicates and consistent grouping of the 1000-cell MAbID samples with the corresponding ChIP-seq datasets obtained from millions of cells (Fig. 1b and Extended Data Fig. 1e). Genome-wide MAbID signal correlates best with ChIP-seq data of matching histone PTMs, with mean Pearson’s correlation coefficients ranging from 0.18 to 0.47 for active chromatin types and 0.24 to 0.47 for heterochromatin types (Fig. 1c and Extended Data Fig. 1f). On a local scale, the MAbID profiles across the linear genome show the expected patterns of enrichment and similarity to ChIP-seq or DamID datasets (Fig. 1d and Extended Data Fig. 1g). To further explore the on-target specificity of MAbID, we calculated the enrichment of MAbID signal over relevant genomic regions, such as ChIP-seq peak calls or ChromHMM domains^33^. All epitopes mapped with MAbID displayed an increase in signal over the matching genomic regions, irrespective of their chromatin type (Fig. 1e). For histone PTMs H3K4me3 and H3K36me3, associated with active gene expression, the level of MAbID signal also scales according to the transcriptional activity of these genes (Fig. 1f).

Finally, we determined the resolution of MAbID by quantifying the signal distribution of H3K4me3 over transcription start sites (TSS) and H3K27me3 over Polycomb-group domains (based on ChromHMM domain calls) and comparing this to the corresponding ChIP-seq datasets (Extended Data Fig. 1h). We found that for H3K4me3 the signal decays to 50% (compared to 100% at the TSS) at 3-4 kb distance from the top of the peak. For H3K27me3, this distance corresponds to 7-8 kb around the domain border. Compared to ChIP-seq, MAbID signal thus generally extends an additional 1-2 kb in either direction. In summary, these results show that with MAbID, we have developed a new method to accurately profile diverse epigenetic modifications and chromatin-binding proteins in as little as 1000 cells.

### Multiplexing several antibodies in one sample without loss of individual data quality

MAbID is designed to enable multiplexing of several antibodies in the same sample and thereby profile many epigenetic landscapes together. To test this, we performed experiments in K562 cells in which we combined four antibodies of different host-origin, along with uniquely barcoded secondary antibody DNA-conjugates specific for each host (Fig. 2a). Since most of the commercially available antibodies are raised in rabbits, we only included the H3K27me3 (rabbit) and RNA Polymerase II (CTD Ser5-phosphorylated, rat) antibodies from the initial dataset. In addition, we used two antibodies of mouse and sheep origin that target H3K36me3 and histone H3 respectively (Supplementary Table 1).

**Fig. 2.**
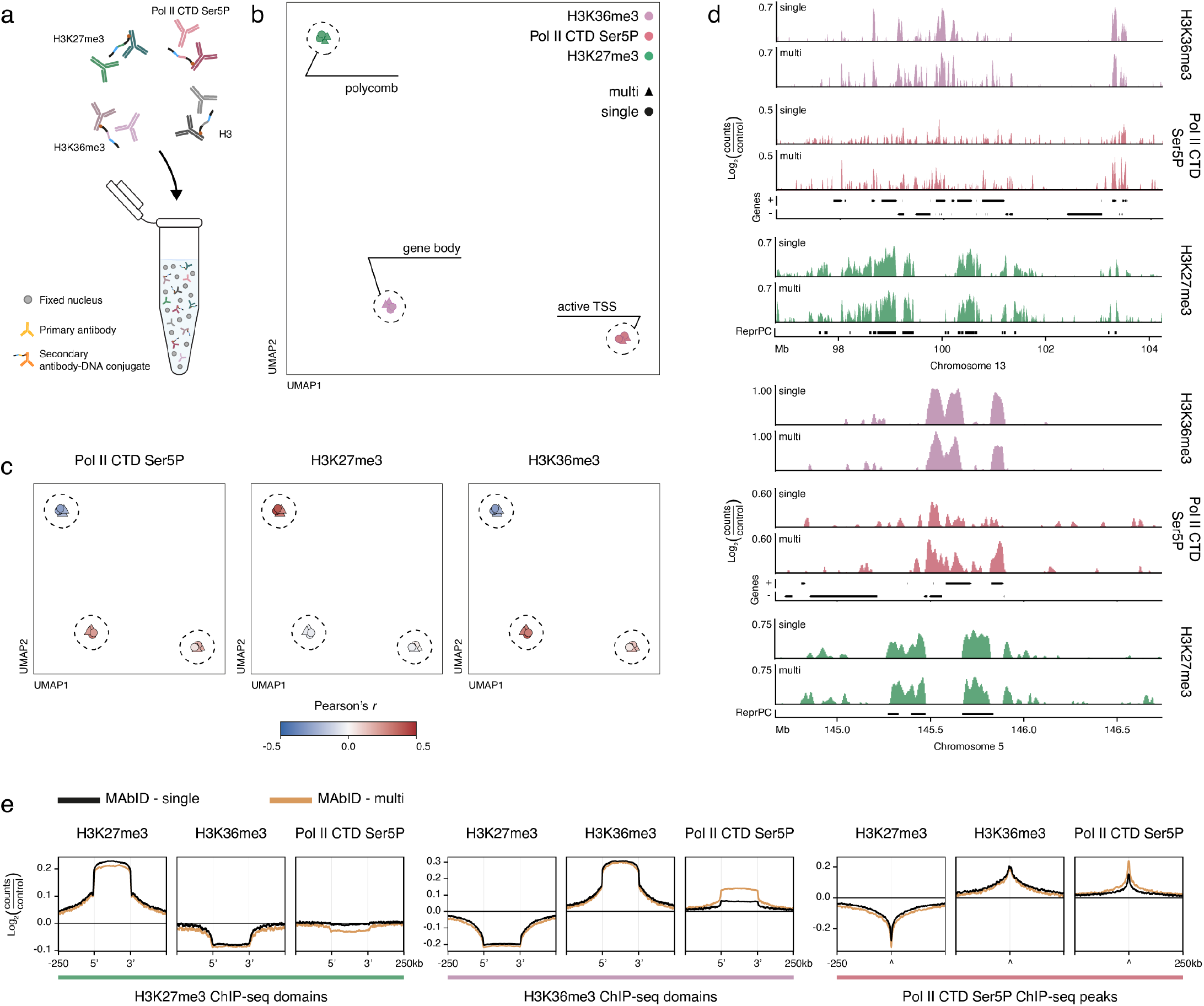
MAbID enables multiplexing of several antibodies on one sample. **a)** Schematic showing the multiplexing of primary antibodies from different species-of-origin with several species-specific secondary antibody-DNA conjugates. **b)** UMAP of MAbID replicates from combined (multi) or individual (single) measurements. Colouring is based on the epitope of interest, chromatin types are encircled. **c)** UMAP as in (b), coloured by correlation with ChIP-seq samples of Pol II CTD Ser5P, H3K27me3 and H3K36me3. The colour scale represents the Pearson’s *r* correlation coefficient of MAbID samples with the indicated ChIP-seq sample. **d)** Genome browser tracks at a broad (top) or narrow (bottom) genomic scale comparing MAbID samples from combined (multi) or individual (single) measurements. Genes (+, forward; -, reverse) and ChromHMM domain calls (ReprPC, Repressed PolyComb) are indicated. Values on the y-axis reflect positive log_2_(counts/control) values. **e)** Average MAbID signal enrichment of H3K27me3, H3K36me3 and Pol II CTD Ser5P around the same domains/peaks called on ChIP-seq data (-/+ 250kb), comparing MAbID samples from combined (multi) or individual (single) measurements.

To unbiasedly assess data quality between multiplexed and individual measurements, we generated MAbID data from samples that were either incubated with each antibody individually (single) or with a combination of all antibodies simultaneously (multi). UMAP embedding shows that samples group based on epitope with high concordance between biological replicates (Fig. 2b). Importantly, this is independent of whether the incubation was done in an individual or combined set-up. The yield and statistics of the processed reads are very comparable between MAbID experiments performed with individual or combined antibody incubations (Extended Data Fig. 2a-b). Moreover, the genome-wide correlation coefficients with ChIP-seq data of corresponding targets are generally independent of the number of multiplexed antibodies (Fig. 2c and Extended Data Fig. 2c). This suggests that competition between antibody DNA-conjugates over antibody-binding sites or restriction-ligation motifs does not influence genome-wide measurements for this set of epitopes.

For H3 specifically, we observed an unexpectedly high correlation with H3K27me3 ChIP-seq data for the samples in which all four antibodies were combined, but not for the individually incubated samples (Extended Data Fig. 2c). We anticipate that cross-reactivity between the anti-sheep secondary antibody-DNA conjugate and primary rabbit H3K27me3 IgG causes this correspondence. We therefore excluded H3 from subsequent analysis and only added it to function as a crowding reagent during following experiments. The unaltered correlation coefficients of the other epitopes verify the specificity of the other secondary antibody-DNA conjugates for their respective target species (Extended Data Fig. 2c).

Next, we visualized the distribution of MAbID signal across the genome to evaluate the impact of combining measurements on a local level. The signal of individual and combined samples shows matching patterns of enrichment on both broad and more narrow genomic scales (Fig. 2d). When comparing both of these MAbID sample types to publicly available ChIP-seq data, the signal enrichments are highly similar and located at the expected genomic regions (Extended Data Fig. 2d). Furthermore, the enrichment of signal over corresponding genome-wide ChIP-seq peaks is similar for individual and combined samples (Fig. 2e).

For the signal enrichments over active genes, we noted that MAbID signal from the Polymerase II CTD Ser5P antibody is not only strongly enriched at the TSS, but also along the gene body (Extended Data Fig. 2e). Since the CTD-Ser5 residue is reported to be phosphorylated on initiating and early elongating RNA Polymerases^34,35^, it may indicate that this antibody has broader affinity for other CTD phospho-modifications. Regardless, the signal is specifically enriched at the most highly transcribed genes and is therefore representative of active gene expression (Extended Data Fig. 2e). Upon comparison of the individual and combined samples, the Polymerase II CTD Ser5P antibody even displays an increase in average signal enrichment over ChIP-seq peaks and active genes (Fig. 2e and Extended Data Fig. 2e). Together, these results verify that antibodies retain on-target specificity and equal data quality between individual and combined measurements. Thus, MAbID enables robust identification of the genomic localization of several different epitopes in a multiplexed assay.

### Genomic integration of the antibody-adapter can be tailored to the epitope of interest

After confirming the ability to multiplex measurements, we sought to increase the modularity of MAbID by adopting another pair of restriction enzymes in addition to MseI & NdeI. This addition allows the MAbID approach to be tailored to the epitope of interest by increasing the theoretical resolution and potentially enhancing the signal. We selected the combination of MboI & BglII to target GATC motifs, because of i) the high efficiency of MboI to digest cross-linked chromatin^36,37^, ii) the different genomic distribution of the GATC motif compared to the TTAA motif (Extended Data Fig. 3a) and iii) the compatibility of the nucleotide overhangs that remain after MboI and BglII digestion. This format enables mixing of secondary antibody-DNA conjugates that are compatible with either pair of restriction enzymes in a single reaction.

We explored this extended strategy in K562 cells by using the H3K36me3, H3K27me3 and Polymerase II CTD Ser5P antibodies along with species-specific secondary antibody-DNA conjugates. Based on motif enrichments per chromatin state (Extended Data Fig. 3a), we generated secondary antibody-DNA conjugates with BglII-compatible adapters for H3K27me3 and Pol II CTD Ser5P and NdeI-compatible adapters for H3K36me3. To make a comprehensive comparison, we tested these in individual (TTAA; GATC) or combined (TTAA or GATC) measurements. A combined sample in which both types of secondary antibody-DNA conjugates were added for all epitopes (TTAA and GATC) was also included, since this combination increases the total number of potential adapter integration sites. All sample types group based on epitope and display the expected enrichment of signal, regardless of the choice of recognition motif or the number of multiplexed antibodies (Extended Data Fig. 3b-c). Both the resolution and distribution of MAbID signal were surprisingly independent of the choice of antibody-adapter and the complexity of the data was similar between all approaches (Extended Data Fig. 3c-d). These outcomes underscore the robustness and flexibility of MAbID. The modular design of the antibody-DNA conjugates offers opportunities to design experiments in accordance with the genomic distribution of the targets. In the following experiments, we therefore matched the choice of antibody-adapter and restriction enzyme-pair with the epitope of interest.

### Primary antibody-DNA conjugates increase the multiplexing potential of MAbID

Next, to increase the number of multiplexed measurements, we explored the potential of directly conjugating the antibody-adapter to primary rather than secondary antibodies. This is more challenging, because i) only one antibody-DNA conjugate can bind per epitope, versus many secondary antibodies binding to a primary antibody and ii) the conjugation procedure could potentially affect the epitope-binding site of monoclonal primary antibodies. Nevertheless, using primary antibody-DNA conjugates eliminates the dependency on antibody host-origins and therefore vastly increases the number of epitopes that can be examined in the same sample. We selected primary antibodies against a variety of chromatin types and conjugated each to a uniquely barcoded antibody-adapter (Extended Data Fig. 3e). The conjugation procedure was slightly modified to account for differences in buffer compositions and the type of antibody-adapter was selected based on the relative TTAA or GATC motif enrichment of the epitope (Extended Data Fig. 3a).

We performed MAbID using individual primary antibody-DNA conjugates in biological replicates of 1000 K562 nuclei (Fig. 3a). The genomic profiles largely overlap with those from ChIP-seq and display general similarity to corresponding MAbID samples obtained with secondary antibody-DNA conjugates (Fig. 3b-c). As expected, the MAbID signal amplitudes and signal-to-noise ratios are lower with primary antibody-DNA conjugates compared to MAbID performed with secondary antibodies. This is most apparent for active histone PTMs, particularly for H3K4me3 and H3K36me3. We presume this relates to the narrower genomic windows in which these types of epitopes generally reside. Nevertheless, upon UMAP embedding these samples group in accordance with their chromatin type and their respective secondary antibody-DNA conjugate sample (Fig. 3d). Moreover, the yield and statistics of the processed reads are as expected and there is a high correlation between biological MAbID replicates (Extended Data Fig. 3f-g). We examined the epitope-specificity of the primary antibody-DNA conjugates further by comparing the signal enrichment to that of MAbID samples using secondary antibody-DNA conjugates (Fig. 3e). The distribution of signal around respective ChromHMM or ChIP-seq domains is highly similar between both sample types, validating that the antibodies maintain their specificity after conjugation. Combined, these results confirm that using primary antibody-DNA conjugates with MAbID generates specific genomic profiles for different chromatin types.

**Fig. 3.**
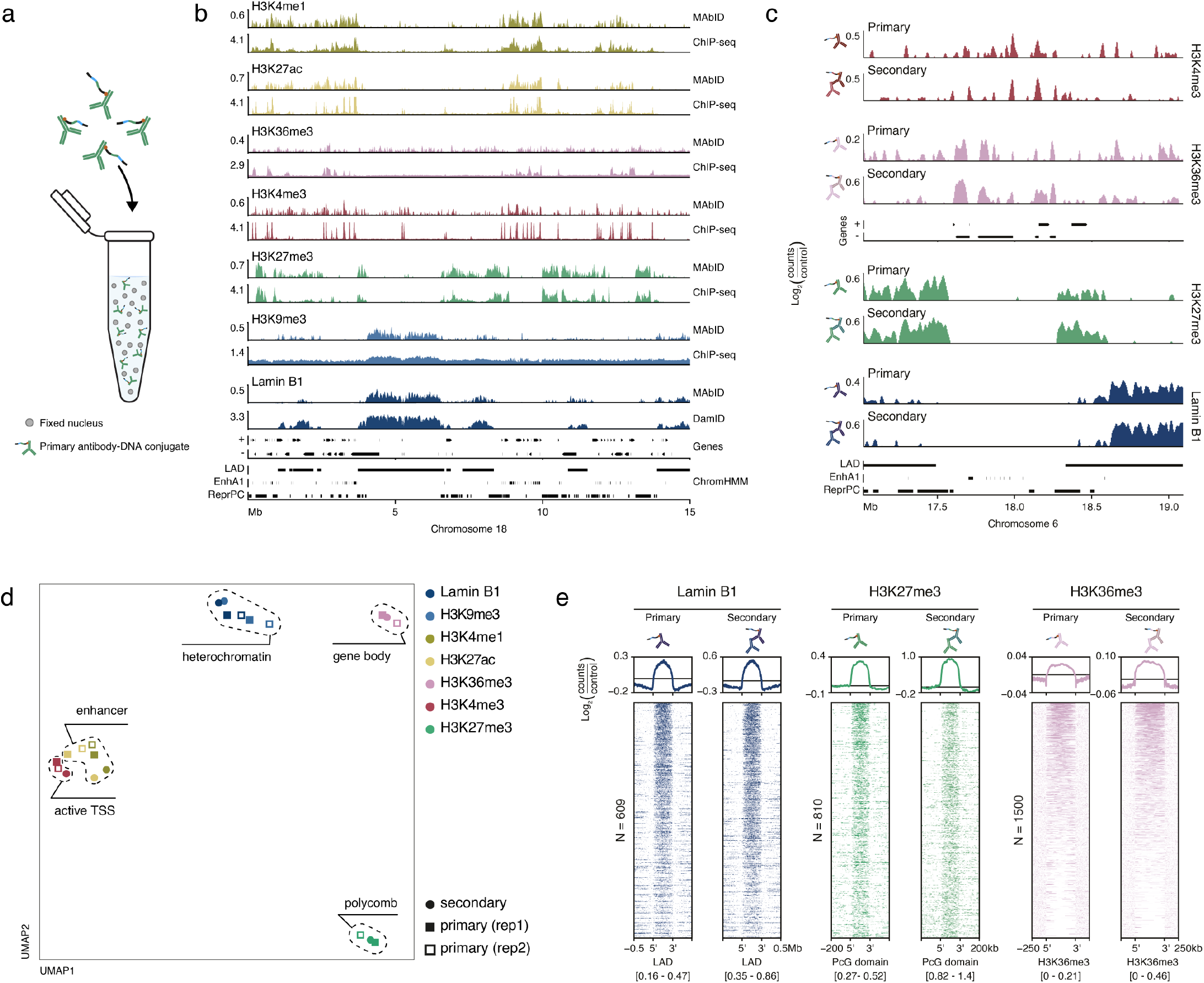
Expanding MAbID with primary antibody-DNA conjugates. **a)** Schematic showing nuclei incubation with primary antibody-DNA conjugates. **b)** Genome browser tracks comparing MAbID samples using primary antibody-DNA conjugates with ChIP-seq or DamID samples. Genes (+, forward; -, reverse) and ENCODE/ChromHMM domain calls (LAD, Lamina-associated domain; EnhA1, Active enhancer 1; ReprPC, Repressed PolyComb) are indicated. Values on the y-axis reflect positive log_2_(counts/control) values for MAbID and DamID and fold change (IP/input) for ChIP-seq. **c)** Genome browser tracks comparing MAbID samples using primary antibody-DNA conjugates or secondary antibody-DNA conjugates (in combination with a primary antibody). Genes and ENCODE/ChromHMM domain calls are indicated. Scaling is based on the minimum and maximum value per sample and values on the y-axis reflect positive log_2_(counts/control) values. **d)** UMAP of MAbID replicates using primary antibody-DNA conjugates and a MAbID sample of merged replicates using secondary antibody-DNA conjugates (in combination with a primary antibody). Colouring is based on the epitope of interest, chromatin types are encircled. **e)** MAbID signal enrichment of Lamin B1, H3K27me3 and H3K36me3 around the same domains/peaks from ChromHMM or ChIP-seq data, comparing MAbID samples using primary or secondary antibody-DNA conjugates. Top line plot shows the average enrichment of signal, bottom heatmap shows signal per genomic region (sorted on MAbID signal). The number (N) of genomic regions included per heatmap is indicated. LAD regions, ENCODE, -/+ 0.5Mb; Polycomb-group domains, ChromHMM, -/+ 200kb; H3K36me3, ChIP-seq domain calls (-/+ 250kb). The heatmap data range is indicated underneath.

We then investigated the performance of primary antibody-DNA conjugates in a multiplexed setting. Based on the data quality of the individual measurements, we chose the best primary antibody-DNA conjugates encompassing a comprehensive set of chromatin types. K562 cells were simultaneously incubated with six primary antibody-DNA conjugates and sorted as samples of 100 nuclei in 384-well plates. The combined MAbID samples group together with the previously generated individual samples in UMAP space, verifying the similarity between the sample types (Extended Data Fig. 3h). Genome-wide correlations with ChIP-seq are comparable between individual and combined measurements, similar to our previous observations for multiplexed samples (Extended Data Fig. 3i). Thus, direct conjugation of antibody-adapters to primary antibodies potentiates profiling of an increasingly complex set of histone PTMs and chromatin-binding proteins.

### scMAbID measures combinatorial epigenetic landscapes at single-cell resolution

We have previously optimized single-cell genomic profiling methods using plate-based protocols, which are very compatible with MAbID. We therefore sought to integrate these protocols with MAbID to generate combined epigenomic measurements at a single-cell resolution (scMAbID). This encompassed sorting in 384-well plates and using liquid-handling robots to increase throughput and reduce sample handling (Fig. 4a). To investigate whether scMAbID can discern chromatin states in different cell types, we differentiated mouse embryonic stem cells (mESCs) for five days towards early Neural Progenitor Cells (early NPCs)^38^. Both cell types were incubated with a combined set of six primary antibody-DNA conjugates targeting a range of chromatin types. In addition, human K562 cells were incubated with the same set of conjugates and also sorted into each well to benchmark scMAbID against the bulk datasets (Fig. 4a). Reads are subsequently assigned to either the human or mouse genome during computational analysis. When testing this approach with control samples, the median number of misannotated reads was below 0.4%, indicating that this is a robust way to assign the cell of origin (Extended Data Fig. 4a).

**Fig. 4.**
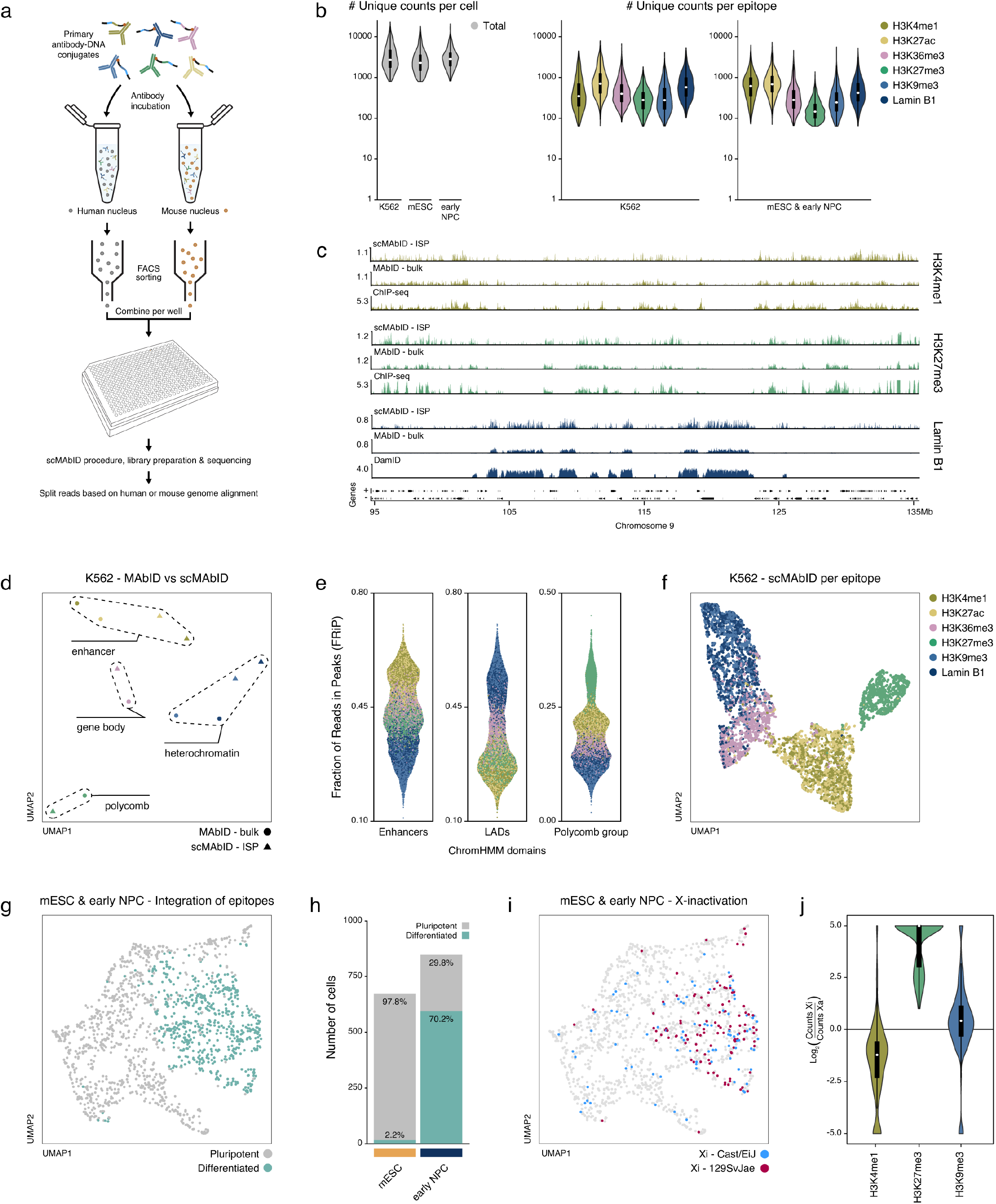
Integration of six multiplexed MAbID measurements in single cells. **a)** Schematic representation of the scMAbID experiment. K562 (human), mESC and early NPC (both mouse) cells are incubated separately with a combination of six primary antibody-DNA conjugates. Single nuclei of each cell type are sorted into a well of a 384-well plate, so that each well contains one nucleus of human origin and one of mouse origin. After performing the scMAbID procedure, library preparation and sequencing, the cell type of origin is determined by alignment to a hybrid genome. **b)** Violin plots showing the total number of unique scMAbID counts per cell and the number of unique counts per epitope within one cell for all cell types. White dot represents the median, boxes indicate the interquartile range. **c)** Genome browser tracks comparing K562 scMAbID ISP (n=1248) and bulk K562 MAbID samples of H3K4me1, H3K27me3 and Lamin B1 with ChIP-seq or DamID samples. Genes (+, forward; -, reverse) are indicated. Values on the y-axis reflect positive log_2_(counts/control) values for MAbID and DamID and fold change (IP/input) for ChIP-seq. **d)** UMAP of K562 scMAbID ISP (n=1248) and bulk K562 MAbID samples. Colouring is based on the epitope of interest and chromatin types are encircled. ISP, in silico population. **e)** FRiP scores of each scMAbID epitope measurement per K562 cell. FRiP scores are calculated across different ChromHMM domains - Enhancers (EnhA1), LADs and Polycomb group (PcG domain). **f)** UMAP of K562 scMAbID per epitope. Each dot represents one epitope measurement, so each cell is represented six times. Colouring is based on the epitope of interest. Only samples (n=6729) passing the threshold of 150 unique counts per epitope were included. **g)** UMAP of integrated mouse scMAbID samples (mESC, n=674; early NPC, n=849). An integrated dataset was computed with all six epitope measurements, so each cell is represented once. Colouring is based on Leiden algorithm cluster assignments, which were labelled as ‘pluripotent’ or ‘differentiated’. **h)** Barplot showing the number of cells from each cell type assigned to the two clusters of (g). Values indicate the percentage of mESCs or early NPC cells assigned to each cluster. **i)** UMAP as (g), with colours reflecting the assigned inactive X-chromosome allele (Xi) based on the ratio of H3K27me3 counts between the two X-alleles. **j)** Violin plots showing the ratio of unique counts of the Xi (inactive X-allele) versus the Xa (active X-allele) per cell for H3K4me1, H3K27me3 and H3K9me3. Values on the y-axis reflect log_2_(counts Xi/counts Xa). White dot represents the median, boxes indicate the interquartile range.

A total number of 1956 K562, 1424 mEsC and 1424 early NPC cells were sequenced and of these respectively 1248, 674 and 849 cells passed the quality thresholds (Extended Data Fig. 4b). Thus, after filtering we still retained 47% to 63% of all sequenced cells, even though the relative complexity of scMAbID data is lower than for bulk MAbID (Extended Data Fig. 4b). The median number of unique counts per cell after filtering was 2715 for K562, 2281 for mESC and 2842 for early NPCs, with per epitope a median of unique counts ranging from 119 to 706 in each cell (Fig. 4b). These numbers are in a similar range to those reported by other recent methods that measure two or three histone PTMs simultaneously^21,26^ (Extended Data Fig. 4c).

First, to assess the quality and specificity of the data, the unique reads of all K562 cells were combined to generate in silico populations (ISP). The scMAbID ISP profiles display a comparable distribution to their matching bulk MAbID, ChIP-seq or DamID dataset (Fig. 4c). The correspondence between the scMAbID ISP and reference datasets is moderately lower than we previously observed for bulk MAbID. We surmise this to be a consequence of the lower relative read numbers that are obtained from the single-cell measurements. Nevertheless, visualization using UMAP shows that the scMAbID ISP samples group with the respective MAbID 1000-cell counterparts, verifying the genome-wide similarity between these datasets (Fig. 4d). To further assess the specificity of the data at single-cell resolution, we calculated FRiP (Fraction of Reads in Peaks) scores for each epitope in single cells using ChromHMM domains as a reference. High FRiP scores are observed for epitope measurements at the corresponding domain, while these are considerably lower at unrelated chromatin types (Fig. 4e). Thus, scMAbID enables obtaining joint measurements of six epitopes in single cells.

After verifying the overall quality of the scMAbID data, we determined if the epitope-specific information from each individual cell enables separation of samples by chromatin state. We took all epitope-measurements that pass a threshold of 150 unique counts per cell (n=6729) and embedded these within UMAP space. The epitope-specific samples consistently separate based on chromatin type and similar chromatin types measured with different antibodies mix together (Fig. 4f). This is independent of the read depth per cell or epitope, indicating that the information is specific to the used primary antibody-DNA conjugate (Extended Data Fig. 4d). Together, these results validate the ability of scMAbID to distinguish epigenetic profiles at a single-cell resolution in a multiplexed set-up.

### Multifactorial chromatin states can be identified by integrating epigenomic measurements

Next, we explored whether scMAbID can discern closely related mESC and early NPCs based on their multiplexed epigenomic profiles. Combining all scMAbID samples of mESCs into in silico populations (ISP) generates genomic profiles following the expected patterns of enrichment with similarity to ChIP-seq or bulk MAbID profiles (Extended Data Fig. 4e). The FRiP scores of single-cell epitope measurements in both cell types are higher for the corresponding chromatin states when compared to other regions (Extended Data Fig. 4f). Moreover, the single-cell epitope measurements with the highest depth per epitope (mESC, n=1800; early NPC, n=1800) mainly separate based on their respective chromatin type (Extended Data Fig. 4g). Overall, these results indicate that scMAbID generates epitope-specific measurements in mESC and early NPC cells.

We noticed that while the UMAP embedding is strongly driven by chromatin signatures, there is already a slight separation between mESC and early NPCs within the same chromatin types (Extended Data Fig. 4g). We wondered whether integration of all multiplexed epitope measurements per cell would improve differentiating these cell types. To integrate the six modalities, we computed a dataset containing the combined epigenetic information for each cell, based on the theoretical work of Zhu et al. from 2021^16^. Briefly, cell-similarity (Jaccard) matrices were calculated per epitope, whereafter dimensionality reduction was performed on the summed matrices. Upon cluster assignment, 97.8% of the mESCs and 70.2% of early NPCs are assigned to their cellular origin based on their integrated chromatin signatures (Fig. 4g-h and Extended Data Fig. 4h). This confirms that the multifactorial chromatin profiles contain sufficient information to accurately separate closely related cell types. Interestingly, 29.8% of the early NPC cells are annotated as mESCs, presumably because these cells failed to differentiate or maintained a more pluripotent state (Fig. 4h). We subsequently labelled the assigned clusters as ‘pluripotent’ and ‘differentiated’.

We then wanted to assess the contribution of each modality to the assignment of cell states. To examine this, we used the Information Gain metric (IG)^39^ to systematically determine the accuracy of cluster assignments with reduced sets of epitopes. We compared these to the original prediction of mESCs assigned to the pluripotent cluster and early NPCs in the differentiated cluster to calculate the IG per epitope-set. The IG performance-metric improves with the inclusion of additional modalities, underlining the added value of multiplexing epitope measurements (Extended Data Fig. 4i). Unsurprisingly, H3K27me3, H3K4me1 and H3K27ac contributed most to the assignment of the two clusters, as these are reported to be valuable predictors of cell type and developmental stage (Extended Data Fig. 4j)^16,20^.

### scMAbID captures changes in chromatin signatures during X-chromosome inactivation

Female mESCs undergo X-chromosome inactivation (XCI)^40^ upon differentiation. This process involves major changes in the distribution of several histone PTMs over the inactivated X-chromosome. We therefore wondered whether we could identify which cells had undergone XCI and examine corresponding changes in multifactorial chromatin signatures. Since this developmental phenomenon occurs randomly, establishing which allele has been inactivated requires both single-cell information and distinctive features between the two alleles. Our female mESCs originate from a hybrid cross between mice from two distinct genotypes (Cast/EiJ × 129SvJae), enabling paternal and maternal genomes to be assigned from frequent SNPs.

Upon random inactivation, the inactive X-allele (Xi) is associated with a marked increase in H3K27me3 levels compared to the allele that remains active (Xa)^41,42^. Therefore, we calculated the ratio of unique H3K27me3 counts between the two X-alleles to establish whether the cells had undergone XCI and which allele had been inactivated. When projecting this status onto the UMAP, we observe that the cells that have inactivated one of the X-alleles are mostly present in the differentiated cluster as expected (Fig. 4i and Extended Data Fig. 4k). The genomic H3K27me3 ISP profiles over the X-chromosome (as well as several of the Hox clusters) also show an overall increase in H3K27me3 levels in the cells of the differentiated cluster (Extended Data Fig. 4l). Interestingly, of the early NPC cells that were labelled as pluripotent (Fig. 4g) only 4.0% have undergone XCI, compared to 27.0% of the early NPCs assigned to the differentiated cluster. This implies that many of these cells indeed failed to proceed through the differentiation trajectory.

Finally, based on the classification of the Xi-allele by the H3K27me3 ratio, we used the multiplexed measurements to determine the occupancy of H3K4me1 and H3K9me3 on the Xi compared to the Xa-allele. The levels of H3K4me1, marking enhancer regions, are decreased on the inactivated allele as expected during the early stages of XCI^43,44^ (Fig. 4j). At the same time, H3K9me3 levels are increased, which has previously been reported to be deposited upon accumulation of Xist RNA on the inactive X-chromosome^43,45^ (Fig. 4j). These results highlight the potential of MAbID to capture single-cell multifactorial dynamics in chromatin states along differentiation trajectories.

## Discussion

The recent advancement of multi-omics strategies at single-cell resolution has created ample opportunity to obtain a deeper understanding of gene regulatory processes^9–11,16,19^. However, measuring multiple epigenetic modifications in a combined setting remains a challenge. Here, we introduce MAbID, a method that enables robust simultaneous profiling of histone PTMs and chromatin-binding proteins in single cells for both active and inactive chromatin types. We generated joint readouts of an unprecedented six epitopes in single cells and integrated all measurements into one dataset to effectively capture differences in chromatin states.

Several other methods have recently been developed that successfully generate combined measurements of multiple histone PTMs in single cells, by employing the Tn5 transposase to integrate barcodes into the genome at sites of antibody binding^21,23–26^. Even though Tn5 is highly efficient, it remains challenging to profile histone PTMs enriched in constitutive heterochromatin due to the affinity of Tn5 for open chromatin regions^28^. Instead, MAbID uses restriction-digestion and ligation steps to effectively integrate barcodes into the genome. We successfully profiled several epitopes located at inaccessible chromatin, such as Lamin B1 and H3K9me3. At the same time, MAbID can be further improved to reduce background signals that we observed at accessible chromatin regions. This is most likely the result of the locally increased efficiency of either the digestion or ligation steps. The background can be corrected for by normalization over a control sample (Extended Data Fig. 1c), analogous to normalization approaches in ChIP-seq^46^. Additional technical improvements to reduce off-target signal can be achieved by optimizing blocking reagents, performing extensive antibody titrations or by optimizing restriction-digestion and ligation steps of the protocol.

We initially tested and benchmarked MAbID by employing secondary antibody-DNA conjugates, since these are produced at lower costs and are readily available in buffers that are compatible with the conjugation protocol. The ability to also successfully use primary antibody-DNA conjugates in one combined incubation greatly increases the flexibility and multiplexing potential of the method. To our knowledge, MAbID is the first method to profile a combination of more than three epitopes, even though there is no theoretical or technical limitation towards combining more measurements for similar approaches. In our experience, the quality of the antibody as well as the efficiency of the conjugation to the antibody-adapter were critical in obtaining high data quality. Especially when using monoclonal primary antibodies, it is imperative to validate the binding efficiency and specificity towards the epitope after the conjugation procedure.

A common challenge for all the current methods simultaneously profiling epigenetic modifications is the low coverage obtained from single cells^21,23,24,26^, which is also true for MAbID single-cell data. Even though combined profiling inherently creates a rich dataset, the sparsity of reads hampers studying the relationship between epitopes, for example when investigating co-occupancy. A recent study by Gopalan et al.^21^ has tackled this in bulk samples by identifying reads that contain double epitope-specific barcodes, which reveal colocalizations of these epitopes. Such a strategy is currently not integrated in the MAbID procedure, but could be accommodated within the method with minor adaptations. Increasing the efficiency of recovering ligation events as well as implementing combinatorial-indexing strategies^47^ to increase the throughput can be additional optimizations to the single-cell protocol.

With MAbID, we have developed a new strategy to profile a wide range of chromatin states in low-input material and single cells that is technically simple and distinct from other recent advancements. The possibility to study several histone PTMs and chromatin-binding proteins together in one sample could greatly benefit research with limited amounts of material, where serial experiments are technically unattainable. We anticipate that innovations such as MAbID and other methods will enable researchers to study the combined epigenetic landscape of complex biological systems in a single integrated experiment.

## Acknowledgements

We would like to thank all the members of the Kind laboratory for their comments throughout the project and their critical reading of the manuscript. We thank Klaas Mulder for his input on generating antibody-DNA conjugates and Peter Zeller for helpful comments and suggestions during the development of the MAbID technique. This work was funded by an ERC Consolidator grant (EU ERC CoG-101002885 FateID) and an NWO-ENW VIDI grant (161.339). The Oncode Institute is partially funded by the KWF Dutch Cancer Society. The laboratory of H.K. is supported by MEXT/JSPS KAKENHI (JP18H05527 and JP21H04764), Japan Science and Technology Agency (JPMJCR16G1), and Japan Agency for Medical Research and Development (JP22ama121020). In addition, we would like to thank the Hubrecht Sorting Facility, especially Reinier van der Linden, and the Utrecht Sequencing Facility (USEQ) for providing sequencing service and data. USEQ is subsidized by the University Medical Center Utrecht and The Netherlands X-omics Initiative (NWO project 184.034.019).

## Author contributions

S.J.A.L., K.L.d.L. and J.K. designed the method. S.J.A.L. developed the method and performed all experiments. R.E.v.B. assisted with experiments. R.H.v.d.W, T.K., and S.J.A.L. performed preliminary computational analyses. R.H.v.d.W. performed all formal computational analyses and designed software. S.J.A.L and R.H.v.d.W. validated and curated the data. H.K. provided resources. R. H.v.d.W and S.J.A.L. performed data visualization. S. J.A.L. and J.K. wrote the original draft of the manuscript, which R.H.v.d.W. edited. J.K. conceived and supervised the study, was responsible for project administration and acquired funding.

---

**Extended Data Fig1.**
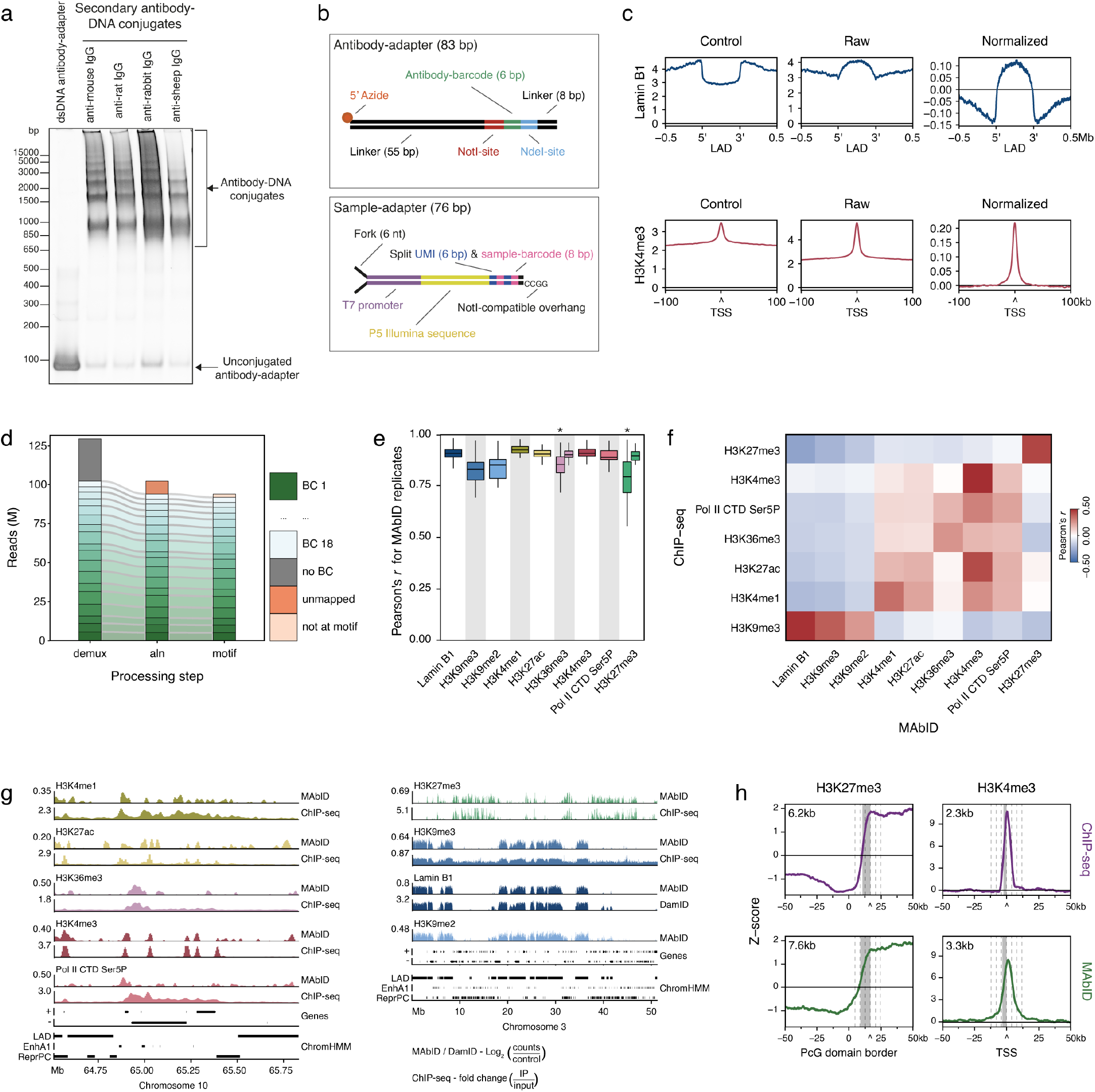
Overview of the MAbID method. **a)** Gel electrophoresis analysis of secondary antibody-DNA conjugates targeting primary IgGs of mouse, rat, rabbit or sheep origin. Conjugates were separated on a native polyacrylamide gel with a 4-12% gradient. Unconjugated antibody-adapter was loaded as a control. **b)** Schematic overview of the designs of the antibody-adapter and sample-adapter. The top strand of the antibody-adapter has a 5’ azide modification (N_3_) for coupling to the antibody. The fork in the double-stranded sample-adapter was created by adding 6 nt non-complementary sequences on the top and bottom strands. UMI, Unique Molecular Identifier; nt, nucleotide; bp, basepair. **c)** Average signal enrichment of MAbID samples using secondary antibody-DNA conjugates, around LAD regions (ENCODE, -/+ 0.5 Mb) or TSS of active genes (-/+ 100 kb). ‘Control’ represents a combination of depth-normalized samples in which the primary antibody was omitted, ‘raw’ is the depth-normalized signal of a MAbID sample using the Lamin B1 or H3K4me3 primary antibody (over LADs or TSS respectively), normalized is the raw sample normalized over the control. Values on the y-axis reflect the counts per million (for control and raw) or log_2_(counts/control) (for normalized). **d)** Barplot showing the number of reads (M, million) retained per computational processing step. Different segments represent separate samples used (BC 1-18), identified based on the combined presence of the sample (SBC) and antibody-barcode (ABBC) within the read. Demux, demultiplexing of reads based on combined barcodes; aln, alignment of reads to the human genome; motif, reads mapping to the TTAA sequence motif. **e)** Boxplots showing the Pearson’s *r* correlation coefficients between corresponding chromosomes of MAbID replicates. Boxplots denoted with an asterisk (*) show the correlation coefficients between corresponding chromosomes of MAbID samples using different primary antibodies against the same epitope. Boxes indicate the interquartile range, center line represents the median. **f)** Heatmap showing the genome-wide Pearson’s *r* correlation coefficient of MAbID samples with different ChIP-seq samples. **g)** Genome browser tracks of MAbID with ChIP-seq or DamID samples for active chromatin types on a narrow genomic scale (left) or inactive chromatin types on a broad genomic scale (right). Genes (+, forward; -, reverse) and ENCODE/ChromHMM domain calls (LAD, Lamina-associated domain; EnhA1, Active enhancer 1; ReprPC, Repressed PolyComb) are indicated. Values on the y-axis reflect positive log_2_(counts/control) values for MAbID and DamID and fold change (IP/input) for ChIP-seq. **h)** Average signal enrichment of MAbID or ChIP-seq samples for H3K27me3 or H3K4me3 around Polycomb-group domain borders (PcG, ChromHMM, -/+ 50 kb, left) or TSS of active genes (-/+ 50 kb, right) respectively. Values on the y-axis reflect Z-score normalized values of log_2_(counts/control) for MAbID and fold change (IP/input) for ChIP-seq. The top of the signal is called at the curve’s inflection point, indicated by ^. Gray box highlights the distance at 50% decay (compared to 100% at the top), which is noted in the top left corner. Dashed lines reflect linear steps of 4 kb distance from the top.

**Extended Data Fig2.**
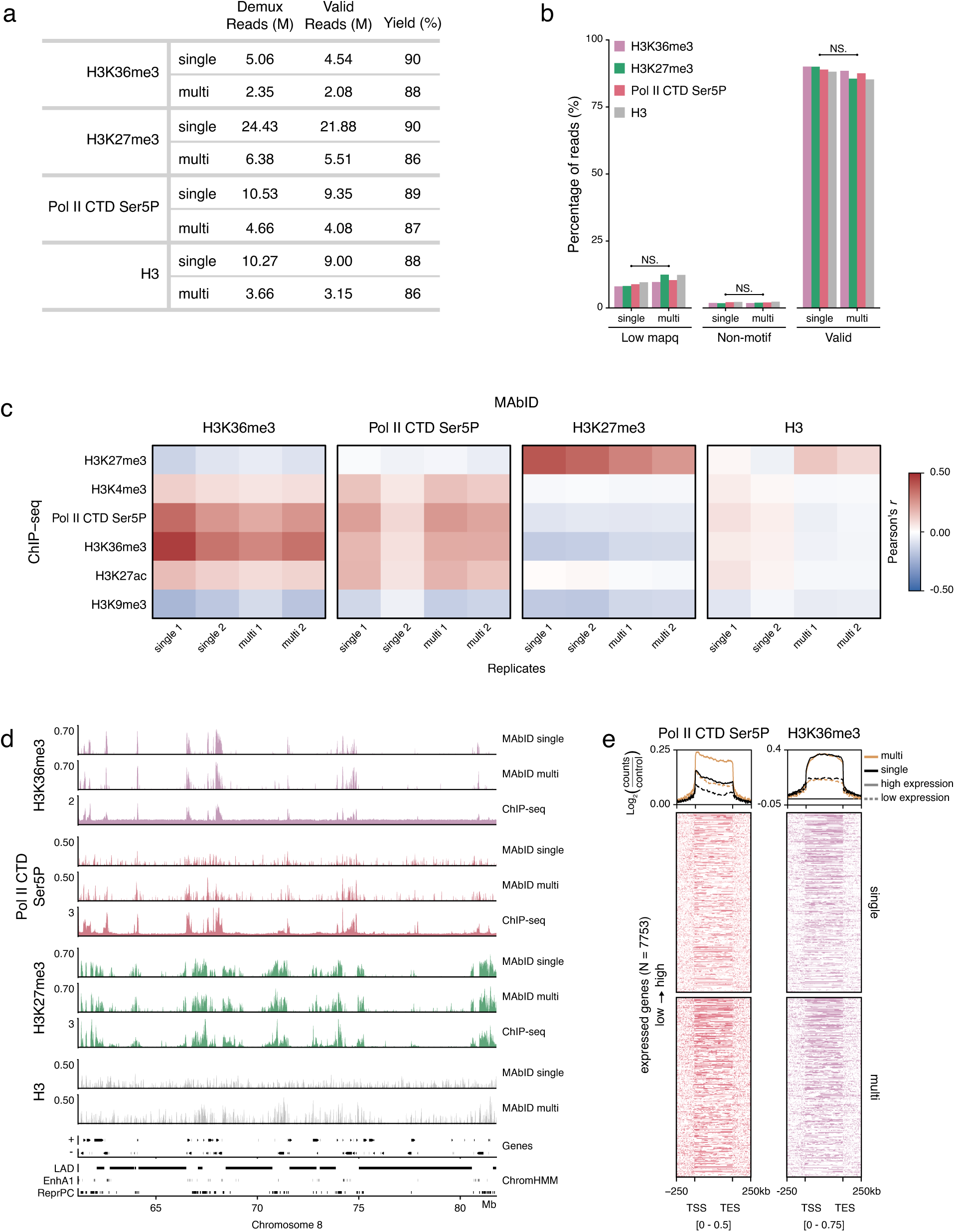
Individual and multiplexed MAbID samples have similar data quality. **a)** Table of the number of demultiplexed reads (demux), valid reads (aligned at TTAA motif) and the resulting yield (% of valid in demux) for individual (single) or combined (multi) samples. M; million. **b)** Barplot showing the percentage of reads lost or retained in the different computational processing steps comparing individual (single) or combined (multi) measurements. Low mapq, low mapping quality; Non-motif, not aligned at TTAA sequence motif; Valid, reads passing quality thresholds aligning at a TTAA sequence motif; NS, non-significant difference based on Wilcoxon signed-rank test. **c)** Heatmap showing the genome-wide Pearson’s *r* correlation coefficient of MAbID replicates of individual (single) or combined (multi) measurements with different ChIP-seq samples. **d)** Genome browser tracks comparing MAbID samples of individual (single) or combined (multi) measurements with ChIP-seq samples. Genes (+, forward; -, reverse) and ENCODE/ChromHMM domain calls (LAD, Lamina-associated domain; EnhA1, Active enhancer 1; ReprPC, Repressed PolyComb) are indicated. Values on the y-axis reflect positive log_2_(counts/control) values for MAbID and fold change (IP/input) for ChIP-seq. **e)** MAbID signal enrichment of individual (single) or combined (multi) measurements of Pol II CTD Ser5P and H3K36me3 around active genes (-/+ 250 kb). Genes were stratified on expression level by setting the top-50% as the “high”-group and the bottom 50% as the “low”-group, line plots indicate the average signal enrichment per group. The heatmap below shows the signal per set of genes based on expression percentiles, ordered from high to low. Heatmaps are split for the individual (top) or combined (bottom) staining; 7753 expressed genes are included per heatmap. The heatmap data range is indicated underneath.

**Extended Data Fig3.**
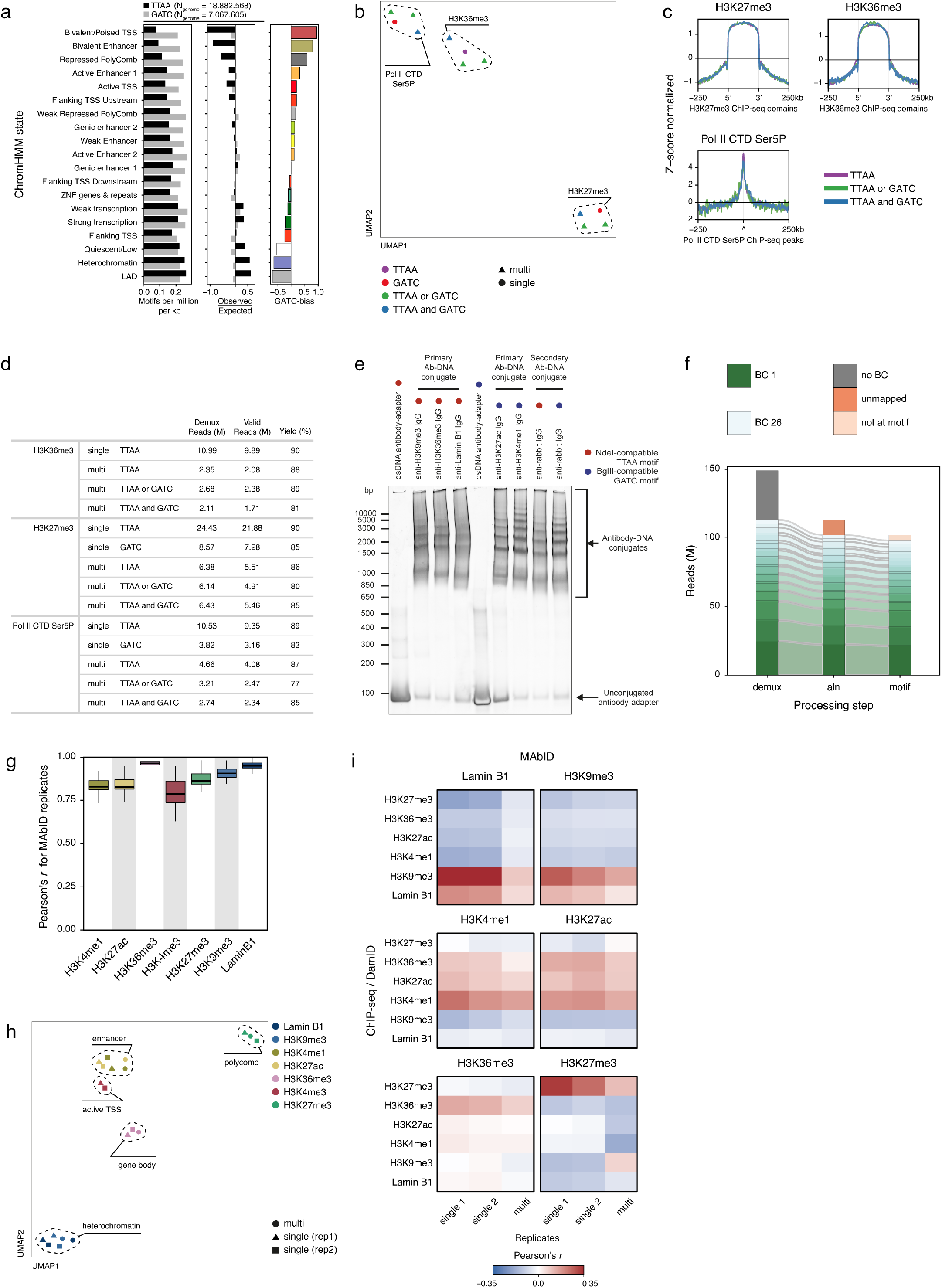
MAbID is customizable to the genomic context of the epitope of interest. **a)** Barplots showing the number of TTAA or GATC sequence motifs in the human genome distributed over ChromHMM states. Observed/Expected (O/E) is the log_2_ transformation of the number of motifs observed compared to the expected number based on the proportion of the genome per state. GATC-bias shows the difference between the O/Es for the GATC versus the TTAA motif. **b)** UMAP embedding of MAbID replicates of individual (single) or combined (multi) measurements using different antibody-adapter types. Colouring is based on the targeted sequence motif by the used antibody-adapter. TTAA, all epitopes are targeted with the NdeI-compatible antibody adapter; GATC, all epitopes are targeted with the BglII-compatible antibody adapter; TTAA or GATC, epitopes are targeted with the NdeI- or BglII compatible adapter; TTAA and GATC, all epitopes are targeted with both types of adapter. The used primary antibodies are encircled. **c)** Average MAbID signal enrichment of H3K27me3, H3K36me3 and Pol II CTD Ser5P around the same domains/peaks called on ChIP-seq data (-/+ 250kb), comparing MAbID samples from combined (multi) or individual (single) measurements using different types of antibody-adapters. Values on the y-axis are Z-score normalized log_2_(counts/control). **d)** Table of the number of demultiplexed reads (demux), valid reads (aligned at TTAA or GATC motif) and the resulting yield (% of valid in demux) for samples using different types of antibody-adapters in individual (single) or combined (multi) samples. M; million. **e)** Gel electrophoresis analysis of primary antibody-DNA conjugates targeting different epitopes. Conjugates were separated on a native polyacrylamide gel with a 4-12% gradient. Unconjugated antibody-adapter was loaded as a control. Red and blue dots indicate the type of antibody-adapter used. **f)** Barplot showing the number of reads (M, million) retained per computational processing step. Different segments represent separate samples used (BC 1-26), identified based on the combined presence of the sample (SBC) and antibody-barcode (ABBC) within the read. Demux, demultiplexing of reads based on combined barcodes; aln, alignment of reads to the human genome; motif, reads mapping to the TTAA or GATC sequence motif. **g)** Boxplots showing the Pearson’s *r* correlation coefficient between corresponding chromosomes of MAbID replicates using primary antibody-DNA conjugates. Boxes indicate the interquartile range, center line represents the median. **h)** UMAP of MAbID samples from combined (multi) or individual (single, two replicates) measurements with primary antibody-DNA conjugates. Colouring is based on the epitope, chromatin types are encircled. **i)** Heatmap showing the genome-wide Pearson’s *r* correlation coefficient of MAbID replicates using primary antibody-DNA conjugates for individual (single) or combined (multi) measurements with different ChIP-seq or DamID (for Lamin B1) samples.

**Extended Data Fig4.**
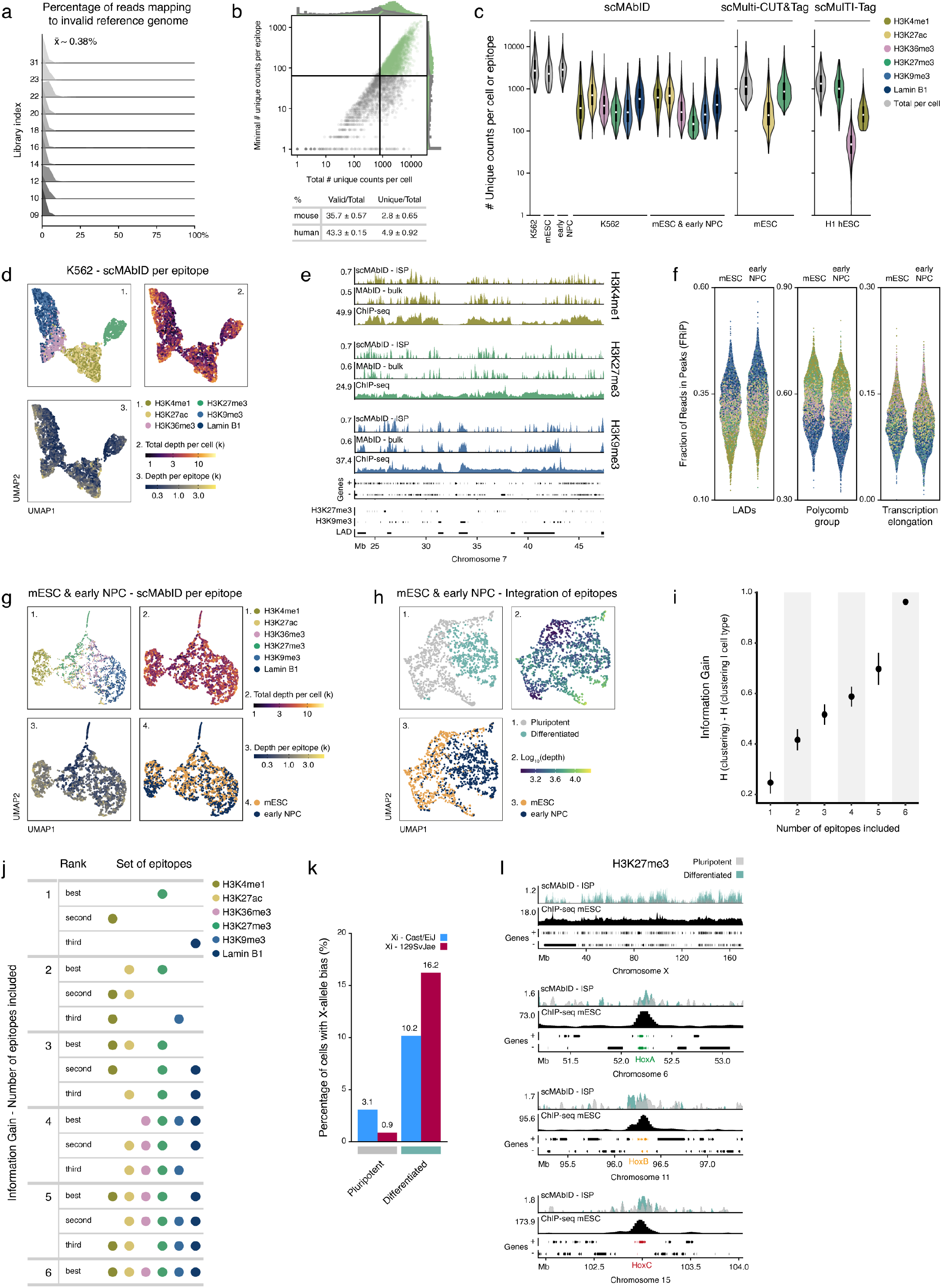
scMAbID data of human and mouse single cells with six multiplexed measurements. **a)** Percentages of reads per well that map to the invalid reference genome for each library index. Calculations are based on wells containing cells of only one origin. The mean percentage across indexes is 0.38%. **b)** Representation of scMAbID cells and reads passing the quality thresholds. The combined dot plot for K562, mESC and early NPC cells shows the total number of unique counts per cell versus the minimal number of unique counts per epitope per cell (the epitope with the lowest count value). Density plots at the top and right sides of the plot indicate the number of samples. Samples passing the quality thresholds are highlighted in green. The table shows the yield (% of reads from total) for valid (at TTAA or GATC motif) or unique (based on UMI) counts for each species. **c)** Violin plots comparing the number of counts or reads between scMAbID, scMulti-CUT&Tag^21^ and scMulTI-Tag^26^. Violins represent the total number of unique counts/reads per cell as well as the number of unique counts/reads per epitope within one cell for the different cell types. White dot represents the median, boxes indicate the interquartile range. **d)** UMAP of K562 scMAbID samples. Each dot represents one epitope measurement from one cell, so each cell is represented six times. Colouring is based on 1) the epitope of interest, 2) total depth per cell or 3) depth per epitope. Only cells passing the threshold of 150 unique counts per epitope were included. k, thousands. **e)** Genome browser tracks comparing mESC scMAbID ISP (n=674) samples of H3K4me1, H3K27me3 and H3K9me3 with bulk MAbID and ChIP-seq. Genes (+, forward; -, reverse) and ENCODE domain calls (H3K27me3 and H3K9me3, ChIP-seq calls; LAD, Lamina-associated domain) are indicated. Values on the y-axis reflect positive log_2_(counts/control) values for MAbID and fold change (IP/input) for ChIP-seq. ISP, in silico population. **f)** FRiP scores of each scMAbID epitope measurement per mESC and early NPC cell. FRiP scores are calculated across different ChromHMM domains - LADs (Lamina-associated domain), Polycomb group (PcG domain) and Transcription Elongation (state 10, TranscriptionElongation). **g)** UMAP of mESC and early NPC scMAbID samples. For each cell type, the top 300 highest depth measurements per epitope are included (mESC, n=1800; early NPC, n=1800). Colouring is based on 1) the epitope of interest, 2) total depth per cell, 3) depth per epitope or 4) cell type of origin. k, thousands. h) UMAP of integrated mouse scMAbID samples (mESC, n=674; early NPC, n=849). An integrated dataset was computed with all six epitope measurements, so each cell is represented once. Colouring is based on 1) Leiden algorithm cluster assignment, 2) total depth per cell (log_10_ transformed) or 3) cell type of origin. i) Information Gain (IG) for different numbers of epitopes included during data integration. The IG metric is calculated by comparing each resulting clustering to the gold standard. The latter is composed of mESC_pluripotent_ and early NPC_differentiated_ based on the original integrated dataset using six epitopes. Whiskers denote the Standard Error of the Mean. j) Table showing the sets of epitopes with the three highest IG values of (i) per number of included epitopes. Colouring is based on epitopes. k) Barplot showing the percentage of cells that have undergone XCI in the pluripotent or differentiated clusters. Bar colours reflect the assigned inactive X-chromosome allele (Xi). Values above bars are percentages of cells inactivating the respective allele. l) Genome browser tracks comparing scMAbID ISP H3K27me3 samples of the pluripotent (n=912) and differentiated (n=611) clusters, along with a mESC ChIP-seq sample of H3K27me3. Genomic regions are the X-chromosome and the regions around Hox clusters A, B and C. Genes (+, forward; -, reverse) are indicated, Hox clusters A, B and C are highlighted with green, orange and red respectively. Values on the y-axis reflect positive log_2_(counts/control) values for MAbID and fold change (IP/input) for ChIP-seq. ISP, in silico population.

## Methods

### Cell culture

All cell lines were grown in a humidified chamber at 37 °C in 5% CO_2_, and were routinely tested for mycoplasma. K562 cells were cultured in suspension on 10-cm dishes in Roswell Park Memorial Institute 1640 (RPMI 1640, Gibco, 61870010) supplemented with 10% FBS (Sigma, F7524, lot BCBW6329) and 1% Pen/Strep (Gibco, 15140122). Cells were passaged every 2-3 days. Mouse F1 hybrid Cast/EiJ (paternal) x 129SvJae (maternal) embryonic stem cells (mESCs; a gift from the Joost Gribnau laboratory) were cultured on 6-well plates with irradiated primary mouse embryonic fibroblasts (MEFs) in mESC culture media (CM) defined as follows: Glasgow’s MEM (G-MEM, Gibco, 11710035) supplemented with 10% FBS, 1% Pen/Strep, 1x GlutaMAX (Gibco, 35050061), 1x MEM non-essential amino acids (Gibco, 11140050), 1 mM sodium pyruvate (Gibco, 11360070), 0.1 mM β-mercaptoethanol (Sigma, M3148) and 1000 U/mL ESGROmLIF (EMD Millipore, ESG1107). mESCs were alternatively cultured in feeder-free conditions on gelatin-coated plates (0.1% gelatin, in house) in 60%-BRL medium, defined as a mix of 40% CM medium (as defined) and 60% conditioned CM medium (incubated 1 week on Buffalo Rat Liver cells), supplemented with 10% FBS, 1% Pen/Strep, 1x GlutaMAX, 1x MEM non-essential amino acids, 0.1 mM β-mercaptoethanol and 1000 U/mL ESGROmLIF. Cells were split every 2-3 days and medium was changed every 1-2 days. This mESC cell line does not contain a Y chromosome. To harvest, cells were washed once with Phosphate Buffered Saline (PBS, in house) and incubated with TrypLE Express Enzyme (Gibco, 12605010) for 3 minutes at 37 °C. Cells were dissociated by pipetting and TrypLE was inactivated by diluting cells five-fold in CM medium, before proceeding with fixation and permeabilization as described.

### Neural differentiation

For differentiation towards the neural lineage (largely following a standard *in vitro* differentiation protocol^38^), mESCs were taken in culture on MEFs and subsequently passaged 3 times in feeder-free conditions in 60%-BRL medium. On day 0 of the differentiation, mESCs were plated on gelatin-coated 6-well plates (0.15% gelatin, Sigma, G1890) at a density of 2.5 x 10^4^ cells per cm^2^ in N2B27 medium defined as follows: 0.5x Dulbecco’s Modified Eagle Medium/Nutrient Mixture F-12 (DMEM-F12, Gibco, 11320033), 0.5x Neurobasal medium (Gibco, 21103049), 15mM HEPES (Gibco, 15630080), 0.5x N-2 supplement (Gibco, 17502048), 0.5x B-27 serum-free supplement (Gibo, 17504044) and 0.1 mM β-mercaptoethanol. From day 3 of differentiation, cells were washed daily with DMEM-F12 medium and refreshed with N2B27 medium. To harvest on day 5 of differentiation, cells were washed once with DMEM-F12 and incubated for 1 minute at room temperature with Accutase Enzyme Detachment Medium (Invitrogen, 00-4555-56). Cells were dissociated by pipetting and Accutase was inactivated by diluting cells ten-fold in DMEM-F12. Cells were centrifuged for 4 minutes at 300 g and resuspended in N2B27 medium, before proceeding with fixation and permeabilization as described.

### Antibodies

For antibodies see Supplementary Table 1.

### ABBC and SBC adapters

The MAbID protocol uses two types of adapters, the antibody-adapter (ABBC) and the sample-adapter (SBC).

#### ABBC adapter

Double-stranded ABBC adapters were conjugated to the antibody via a SPAAC click reaction^30–32^ (see section Antibody-DNA conjugation). The top strand of the double-stranded adapter was produced as HPLC-purified oligo and has a 5’ Azide modification (IDT, /5AzideN/) to allow for antibody conjugation, the bottom strand was produced as standard-desalted oligo. For the NdeI-compatible adapter (TTAA motif, to ligate to MseI-digested genome), the elements in the design were (5’ to 3’) a 55 nt linker, a NotI recognition site, a 6 nt ABBC barcode and a NdeI recognition site. In the BglII-compatible adapter (GATC motif, to ligate to MboI-digested genome), the NdeI recognition site is replaced by a BglII recognition site. Additionally, the adenine of the motif on the bottom strand of the adapter was methylated (IDT, /iN6Me-dA/). The hemi-methylated G^m6^ATC sequence is no longer recognized and digested by MboI, but the modification does not decrease BglII digestion efficiency. The oligo is produced as HPLC-purified. For top and bottom oligo sequences, see Supplementary Table 2. Top and bottom oligos were annealed in a 1:1 ratio at 10 *μ*M final concentration in 1X annealing buffer (10 mM Tris-Cl, pH 7.4, 1 mM EDTA and 100 mM NaCl) in 0.5 mL DNA LoBind tubes (Eppendorf, 0030108400) by incubating in a PCR machine at 95 °C for 5 min, followed by gradual cooling down with 0.5 °C per 15 seconds to 4 °C final.

#### SBC adapters

SBC adapters were designed as forked double-stranded DNA adapters, which can ligate to the ABBC adapters. The bottom adapter has a 5’ Phosphorylation modification (IDT, /5Phos/) and 4 nt GGCC (5’ to 3’) overhang to facilitate ligation to NotI digested DNA. Both top and bottom oligos were produced as standard-desalted oligos. The other elements in the design were (5’ to 3’) a 6 nt non-complementary fork, the T7 promoter, the 5’ Illumina adapter (as used in the Illumina TruSeq Small RNA kit) and a split 2x 3 nt Unique Molecular Identifier (UMI) interspaced with a split 2x 4 nt SBC barcode. For top and bottom oligo sequences, see Supplementary Table 3. Top and bottom oligos were annealed in a 1:1 ratio at 40 *μ*M final concentration in 1X annealing buffer (10 mM Tris-Cl, pH 7.4, 1 mM EDTA and 50 mM NaCl) in a 96-well plate by incubating in a PCR machine at 95 °C for 5 min, followed by gradual cooling down with 0.5 °C per 15 seconds to 4 °C final. Double-stranded SBC adapters were diluted further before use.

### Antibody-DNA conjugation

#### Secondary antibody-conjugates

Secondary antibody-DNA conjugations were performed as described by Harada, A. et al. 2019^25,30^, with minor modifications. Briefly, goat anti-rabbit IgG (Jackson ImmunoResearch, 111-005-114), donkey anti-mouse IgG (Jackson ImmunoResearch, 715-005-150), donkey anti-rat IgG (Jackson Immunoresearch, 712-005-150) or donkey anti-sheep IgG (Jackson ImmunoResearch, 713-005-147) was buffer-exchanged from storage buffer to 100 mM NaHCO3 (pH 8.3) using Zeba™ Spin Desalting columns (40K MWCO, 0.5 mL, ThermoFisher, 87767). Subsequently, 100 *μ*g antibody in 100 *μ*L of 100 mM NaHCO_3_ (pH 8.3) was conjugated with dibenzocyclooctyne (DBCO)- PEG4-NHS ester (Sigma, 764019) by adding 0.25 *μ*L of DBCO-PEG4-NHS (dissolved at 25 mM in dimethylsulfoxide (DMSO, Calbiochem, 317275), 10 times molar ratio to antibody) and incubated for 1 hour at room temperature on a tube roller. The sample was passed through a Zeba™ Spin Desalting column to remove free DBCO-PEG4-NHS and to directly buffer-exchange to PBS. DBCO-PEG4-conjugated antibodies were concentrated using an Amicon Ultra-0.5 NMWL 10-kDa centrifugal filter (Merck Milipore, UFC501024) and measured on a NanoDrop™ 2000. The DBCO-PEG4-conjugated antibody was diluted to 1 *μ*g/*μ*L in PBS. Conjugation of antibody with the ABBC DNA adapter was performed at a molar ratio of 1:2 by mixing 75 *μ*L of DBCO-PEG4-conjugated antibody (75 *μ*g, in PBS) with 100 *μ*L of double-stranded ABBC adapter (10 *μ*M, see section ‘ABBC and SBC adapters’). Samples were incubated at 4 °C for 1 week on a rotor at 8 rpm. Subsequent clean-up of the antibody-DNA conjugate was performed as described by Harada, A. et al. 2019^25,30^, with an average yield of 20-30 *μ*g. The concentration of the antibody-DNA conjugate was measured using the Qubit Protein Assay (Invitrogen, Q33211). Sample quality and conjugation efficiency were assessed using standard agarose gel electrophoresis or Native PAGE with TBE 4-12% gradient gels (Invitrogen, EC62352BOX) stained with SYBR Gold Nucleic Gel stain (Invitrogen, S11494). Gels were imaged using the Amersham Typhoon laser-scanner platform (Cytiva). Antibody-DNA conjugates were stored at 4 °C.

#### Primary antibody-conjugates

Primary antibody-DNA conjugations were performed as described in the previous section ‘Secondary antibody-conjugates’ with minor modifications. Primary antibodies were first cleaned using the Abcam Antibody Purification Kit (Protein A) (Abcam, ab102784) following manufacturer’s instructions (performing overnight incubation at 4 °C in the spin cartridge on a rotor at 8 rpm). All four elution phases were taken along to maximize the yield. Purified antibodies were concentrated using an Amicon Ultra-0.5 NMWL 10-kDa centrifugal filter, after which 350*μ*L 100 mM NaHCO_3_ was added and concentrated again to exchange buffers. The concentrated antibody was measured on the Nanodrop™ 2000. All subsequent steps were performed as described in the section ‘Secondary antibody-conjugates’ from the DBCO-PEG4-NHS incubation onwards.

### Cell harvesting, fixation and permeabilization

Cells were harvested (~10 x 10^6^ cells) and washed once with PBS. All centrifugation steps were at 200 g for 4 minutes at 4 °C. Cells were fixated in 1% formaldehyde (Sigma, F8875) in PBS for 5 minutes, before quenching the reaction with 125 mM final concentration of glycine (Sigma, 50046) and placing the cells on ice. All subsequent steps were performed on ice or at 4 °C. Cells were washed three times with PBS before resuspension in Wash buffer 1 (20mM HEPES pH 7.5 (Gibco, 15630-056), 150 mM NaCl, 66.6 *μ*g/mL Spermidine (Sigma, S2626), 1 X cOmplete™ protease inhibitor cocktail (Roche, 11697498001), 0.05% Saponin (Sigma, 47036), 2mM EDTA) and transferred to a 1.5 mL protein LoBind Eppendorf tube (Eppendorf, EP0030108116-100EA). Cells were permeabilized for 30 minutes at 4 °C on a tube roller. Bovine Serum Albumin (BSA, Sigma, A2153) was added to 5 mg/ mL final concentration and incubated for another 60 minutes at 4 °C on a tube roller. Permeabilized nuclei were used for antibody incubation.

### Antibody incubations

All centrifugation steps were at 200 g for 4 minutes at 4 °C.

#### Primary antibody-conjugates

Permeabilized nuclei (see section Cell harvesting, fixation and permeabilization) were counted on a TC20™ Automated Cell Counter (BioRad, 1450102). Nuclei were diluted to ~2.5 x 10^6^ cells/mL in Wash buffer 1, of which 100 *μ*L (~250,000 nuclei) was used for each primary antibody incubation. Primary antibody conjugated to an ABBC adapter (see Antibody-DNA conjugation section) was added and incubated overnight at 4 °C on a tube roller (see Supplementary Table 1 for antibody concentrations used). For each experiment, a control sample without a primary antibody was taken along. The next morning, the nuclei were washed two times with Wash Buffer 2 (20mM HEPES pH 7.5, 150 mM NaCl, 66.6 *μ*g/mL Spermidine, 1X cOmplete™ protease inhibitor cocktail, 0.05% Saponin) and resuspended in 200 *μ*L Wash Buffer 2 containing Hoechst 34580 (Sigma, 63493) at 1 *μ*g/mL. Nuclei were incubated for 1 hour at 4 °C on a tube roller. Finally, nuclei were washed two times with Wash Buffer 2 and resuspended 500 *μ*L Wash Buffer 2 before proceeding to FACS sorting.

#### Secondary antibody-conjugates

Permeabilized nuclei (see section Cell harvesting, fixation and permeabilization) were counted on a TC20™ Automated Cell Counter. Nuclei were diluted to ~2.5 x 10^6^ cells/mL in Wash buffer 1, of which 200 *μ*L (~500,000 nuclei) is used for each primary antibody incubation. Primary antibody (unconjugated) was added and nuclei were incubated overnight at 4 °C on a tube roller (see Supplementary Table 1 for antibody concentrations). For each experiment, a control sample without a primary antibody was taken along. The next morning, the nuclei were washed two times with Wash Buffer 2 (20mM HEPES pH 7.5, 150 mM NaCl, 66.6 *μ*g/ mL Spermidine, 1X cOmplete™ protease inhibitor cocktail, 0.05% Saponin) and resuspended in 200 *μ*L Wash Buffer 2 containing Hoechst 34580 at 1 *μ*g/mL. Secondary antibody conjugated to an ABBC adapter (see Antibody-DNA conjugation section) was added (2 *μ*g/mL) and incubated for 1 hour at 4 °C on a tube roller. Finally, nuclei were washed two times with Wash Buffer 2 and resuspended in 500 *μ*L Wash Buffer 2 before proceeding to FACS sorting.

### FACS sorting

Nuclei were pipetted through a Cell Strainer Snap Cap into a Falcon 5 mL Round Bottom Polypropylene Test Tube (Fisher Scientific, 10314791) just prior to sorting on a BD Influx or BD FACsJazz Cell sorter. Nuclei were sorted in G1/S cell-cycle phase, based on the Hoechst levels. For 1000-cell samples, nuclei were sorted into a tube of a PCR tube strip containing 5 *μ*L 1X CutSmart buffer (NEB, B7204S) per well. The final volume after sorting was ~7.5 *μ*L per tube. For samples with 100 cells or less, the appropriate number of nuclei was sorted into a 384-well PCR plate (BioRad, HSP3831) containing 200 nL 1X CutSmart buffer and 5 *μ*L mineral oil (Sigma, M8410) per well. Plates were sealed with aluminum covers (Greiner, 676090).

### MAbID procedure

#### Manual preparation of MAbID samples

Samples containing 1000 nuclei were processed in PCR tube strips. Samples were spun briefly in a table-top rotor between incubation steps. 2.5 *μ*L of Digestion-1 mix (MseI (12.5 U, NEB, R0525M) and/ or MboI (12.5 U, NEB, R0147M) in 1X CutSmart buffer) was added to a total volume of 10 *μ*L per tube, including 7.5 *μ*L sorting volume. Samples were incubated in a PCR machine for 3 hours at 37 °C before holding at 4 °C. 5 *μ*L of rSAP mix (rSAP (1 U, NEB, M0371L) in 1X CutSmart buffer (for MseI/NdeI digestions) or 1X NEBuffer 3.1 (NEB, B7203S), for all digestions including MboI/BglII)) was added to a total volume of 15 *μ*L per tube. Samples were incubated for 30 minutes at 37 °C, then 3 minutes at 65 °C before transfer to ice. 5 *μ*L of Digestion-2 mix (NdeI (5 U, NEB, R0111L) and/or BglII (5 U, NEB, R0144L) in 1X CutSmart buffer (for MseI/NdeI digestions) or 1X NEBuffer 3.1 (for all digestions including MboI/ BglII)) was added to a total volume of 20 *μ*L per tube. Samples were incubated for 1 hour at 37 °C, before holding at 4 °C. 6 *μ*L of Ligation-1 mix (3.75 U T4 DNA ligase (Roche, 10799009001), 33.3 mM DTT (Invitrogen, 707265), 3.33 mM ATP (NEB, P0756L) in 1X Ligase Buffer (Roche, 10799009001)) was added to a total volume of 26 *μ*L per tube. Samples were incubated for 16 hours at 16 °C, before holding at 4 °C. 4 *μ*L of Lysis mix (Proteinase K (5.05 mg/mL, Roche, 3115879001), IGEPAL CA-630 (5.05%, Sigma, I8896) in 1X CutSmart buffer) was added to a total volume of 30 *μ*L per tube. Samples were incubated for 4 hours at 56 oC, 6 hours at 65 °C and 20 minutes at 80 °C before holding at 4 °C. 10 *μ*L of Digestion-3 mix (5 U NotI-HF (NEB, R3189L) in 1X CutSmart buffer) was added to a total volume of 40 *μ*L. Samples were incubated for 3 hours at 37 °C before holding at 4 °C. 2.5 *μ*L of uniquely barcoded SBC adapter (550 nM, see section ABBC and SBC adapters) was added to reach a final concentration of ~25 nM during ligation. 12.5 *μ*L of Ligation-2 mix (6.25 U T4 DNA ligase, 34 mM DTT, 3.4 mM ATP in 1X Ligase Buffer) is added to each tube to a final volume of 55 *μ*L during ligation. Samples were incubated for 12 hours at 16 °C and 10 minutes at 65 °C before holding at 4 °C.

#### Robotic preparation of MAbID plates

384-well PCR plates with sorted nuclei were processed using a Nanodrop II robot at 12 psi pressure (BioNex) for adding all mixes. Indicated volumes are per well. Between handling, plates were spun for 2 minutes at 1000 g at 4 °C each time. 200 nl of Digestion-1 mix (MseI (0.5 U) and/or MboI (0.5 U) in 1X CutSmart buffer) was added to a total volume of 400 nL per well. Plates were incubated in a PCR machine for 3 hours at 37 °C before holding at 4 °C. 200 nL of rSAP mix (rSAP (0.04 U) in 1X CutSmart buffer (for MseI/NdeI digestions) or 1X NEBuffer 3.1 (for all digestions including MboI/BglII)) was added to a total volume of 600 nL per well. Plates were incubated for 30 minutes hours at 37 °C, then 3 minutes at 65 °C before directly placing on ice. 200 nL of Digestion-2 mix (NdeI (0.2 U) and/or BglII (0.2 U) in 1X CutSmart buffer (for MseI/NdeI digestions) or 1X NEBuffer 3.1 (for all digestions including MboI/BglII)) was added to a total volume of 800 nL per well. Plates are incubated for 1 hour at 37 °C, before holding at 4 °C. 240 nL of Ligation-1 mix (0.15 U T4 DNA ligase, 33.3 mM DTT, 3.33 mM ATP in 1X Ligase Buffer) was added to a total volume of 1040 nL per well. Plates were incubated for 16 hours at 16 °C, before holding at 4 °C. 160 nL of Lysis mix (Proteinase K (5.05 mg/mL), IGEPAL CA-630 (5.05%) in 1X CutSmart buffer) was added to a total volume of 1200 nL per well. Plates were incubated for 4 hours at 56 °C, 6 hours at 65 °C and 20 minutes at 80 °C before holding at 4 °C. 400 nL of Digestion-3 mix (0.2 U NotI-HF in 1X CutSmart buffer) was added to a total volume of 1600 nL per well. Plates were incubated for 3 hours at 37 °C before holding at 40C. 150 nL of uniquely barcoded SBC adapter (110 nM, see section ABBC and SBC adapters) was added to each well using a Mosquito HTS robot (TTP Labtech) to reach a final concentration of ~7.5 nM during ligation. 450 nL of Ligation-2 mix (0.25 U T4 DNA ligase, 37.8 mM DTT, 3.78 mM ATP in 1X Ligase Buffer) was added to a final volume of 2200 nL during ligation. Plates were incubated for 12 hours at 16 °C and 10 minutes at 65 °C before holding at 4 °C.

### Library preparation

Samples ligated with unique SBC adapters were pooled, either 2-4 1000 nuclei samples or a full 384-well plate were pooled for combined *in vitro* transcription (IVT). To reduce batch effects, controls (secondary antibody-conjugates without primary antibody incubation) were pooled with their corresponding samples whenever possible. For 384-well plates, mineral oil was removed by spinning the sample for 2 minutes at 2000 g and transferring the liquid phase to a clean tube, which was repeated three times. After pooling, samples were incubated for 10 minutes with 1.0 volume CleanNGS magnetic beads (CleanNA, CPCR-0050), diluted 1:4 to 1:10 in bead binding buffer (20% PEG 8000, 2.5 M NaCl, 10 mM Tris-HCl, 1 mM EDTA, 0.05% Tween 20, pH 8.0 at 25 °C). The bead dilution ratio depended on the total volume, 1:4 for 1000 nuclei samples and 1:10 for a full 384-well plate. Samples were placed on a magnetic rack (DynaMag™-2, ThermoFisher, 12321D) to collect beads on the side of the tube. Beads were washed two times with 80% ethanol and briefly allowed to dry before resuspending in 8 *μ*L water. *In vitro* transcription was performed by adding 12 *μ*L IVT mix from the MEGAScript T7 kit (Invitrogen, AM1334) for 14 hours at 37 °C before holding at 4 °C. Library preparation was subsequently performed as described previously^13,48^, using 5 *μ*L of aRNA and 8 to 11 PCR cycles, depending on the aRNA yield. Purified aRNA from different IVT reactions (with unique SBC adapters) can be pooled before proceeding with cDNA synthesis to reduce batch effects. Libraries were run on the Illumina NextSeq500 platform with high output 1×75 bp, the Illumina NextSeq2000 platform with high output 1×100 bp or the Illumina NextSeq2000 platform with high output 2×100 bp.

### Raw data processing

Reads of the raw sequencing output conform to a MAbID specific layout of 5’-[3 nt UMI][4 nt SBC part 1][3 nt UMI][4 nt SBC part 2]AGGGCCGC[8 nt ABBC][genomic sequence]-3’. Raw R1-reads were demultiplexed on the expected barcode-sequences using CutAdapt 3.0^49^, with the following custom settings. First, we only allow matches with at least a 29 nt overlap and only keep reads directly starting with the adapter (i.e., an anchored 5’ adapter). The maximum error rate setting of 2 makes it possible to retain reads with 1) two mismatches in the specified adapter-sequence (ignoring the UMI) and 2) a one nucleotide insertion or deletion (indel) at the start of the read due to digestion-ligation or sequencing(-library) errors. Demultiplexed reads are parsed through a custom script to classify reads on correct adapter-structures on a 7-tiered range. Reads in class 1 adhere perfectly to the barcode-expectations, while class-7 reads only contain the AGGGCCGC-sequence at the expected location. Reads typically fall into class 1 (average for figure 1: 95%). This classification allows fine control over which reads should be retained. For this manuscript, we only allow classes 1 and 2 (1 nucleotide indel in the first UMI) to ensure the highest possible quality. Finally, the script creates a fastq.gz-file, adding the UMI-sequences to the read-ID for downstream processing and removing the adapter sequence.

### Sequence alignments

Demultiplexed and filtered reads were processed in a similar fashion to Rooijers et al.^13,48^, with the additional flexibility to set the selected restriction site motif. Briefly, reads are aligned using Bowtie version 2.4.1^50^ in unpaired mode, using default end-to-end parameters. We used the UCSC hg19 reference genome for K562 samples and the NCBI mm10 reference genome for mESC/early NPC samples (both references were downloaded from https://benlangmead.github.io/aws-indexes). Alignments with a mapping quality lower than 10 or not at the expected ligation site (5’ for the MboI GATC-motif or 5’+ 1 for the MseI TTAA-motif) were discarded. For reads originating from mixed sorting single-cell samples (e.g., K562 and mESCs or early NPC), a new hybrid reference genome was built by concatenating hg19 and mm10. Aligned reads to this reference were subsequently mapped to the individual references for further downstream processing by Bowtie using the --very-sensitive -N 1 parameters. Mouse allele-specific reads were assigned by mapping mm10-reads to 129/Sv and Cast/Eij reference genomes. We designated reads to one of the genotypes if it mapped better (i.e., lower edit-distance or higher alignment score) to one of the references.

### Public data

For the K562 analyses, we downloaded the ChromHMM33 calls and several histone PTM datasets from the ENCODE portal^51,52^ (https://www.encodeproject.org/) with the following identifiers: ENCFF001SWK, ENCFF002CKI, ENCFF002CKJ, ENCFF002CKK, ENCFF002CKN, ENCFF002CKY, ENCFF002CUS, ENCFF002CTX, ENCFF002CUU, ENCFF002CKV, and ENCFF002CUN. Expression values per gene for K562 cells were also downloaded from ENCODE (ENCFF401KET). LAD-annotations for K562 were downloaded from the 4D-nucleome project^53^ and converted to hg19-coordinates with the liftOver utility of UCSC (https://genome.ucsc.edu/cgi-bin/hgLiftOver), while the LAD-annotations of Peric-Hupkes et al.^54,55^ were used for the analyses on mm10 datasets. Data from other recent methods^21,26^ was downloaded in RDS-form from supplied repositories and used as-is.

### MAbID analyses

Aligned reads are UMI-flattened and counted per restriction site, like in the scDam&T-seq protocol^13,48^. We allowed up to 1000 UMIs per site in the bulk analyses and up to 2 UMIs per site in the single-cell analyses. UMI-counts per sample were binned (5kb, 20kb, and 100kb), loaded into R and stored into singleCellExperiment-containers^56^. Counts in bins overlapping regions of known problematic nature (i.e., blacklist-regions^57^) or low mappability are set to zero. scMAbID single-cells (unless otherwise indicated) were filtered for a minimum of 800 UMIs per cell and 64 UMIs per epitope in each cell. Normalization of the data was performed by calculating RPKM-values for both samples and control (see eq. 1) and calculating the fold-change over control with a pseudocount value of 1 (see eq. 2).

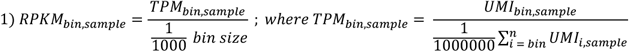

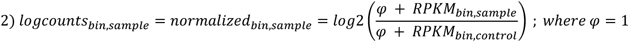

Since the majority of the analyses was performed in R, we created an R-package (mabidR) to load, normalize, and analyze the generated datasets. Genome-wide correlation-analyses were performed on 5kb resolution log_2_(O/E), ignoring blacklisted regions and setting negative values to zero. Replicates were merged after verification that the separate datasets are of high quality and in agreement with each other. Genome browser tracks for bulk MAbID data represent positive log_2_(O/E) values of merged replicate datasets. Enrichment-computations were performed using the computeMatrix tool of Deeptools version 3.5.1^58^ and analyzed in R. Polycomb-group domains were generated by merging 200 bp regions of the ChromHMM states 16 and 17, allowing a gap of 10 kb and filtering the resulting regions on a minimal size of 100 kb. Expression-based stratifications of gene bodies were made by splitting RNA-seq TPM-values on [0,33.4,66.7,100] percentiles, resulting in low/mid/high categories, respectively. Input for the UMAP-analyses was log_2_(O/E) for the bulk- and ISP-approach, and log_1_p(UMI-counts) for single-cell samples. All data was Z-score normalized before PCA. The elbow-method was used to find applicable components as input for UMAP. To limit method-specific accessibility-biases dominating dimensionality reductions (data not shown), only one public dataset (normalized DamID and ChIP-seq) was included per chromatin-state cluster. For the cross-epitope K562 UMAP, we kept epitope-samples with more than 150 UMIs belonging to a cell that has more than 800 UMI counts. Only bins containing UMIs in more than 10 cells were used. For the cross-epitope mouse UMAP, we kept per cell type (mESC and early NPC) the top 300 highest-depth samples for each epitope. Fraction of Reads in Peaks (FRiP) scores of single-cell epitope-measurements are defined as the fraction of counts overlapping a defined region-set. Pan-epitope mouse UMAPs were generated as described in Zhu et al.^16^: for each 100 kb [*bin-cell*] UMI-matrix per epitope, we computed Jaccard-distances (*D_cell i,cell j_* = 1 - *jaccard_cell i,cell j_*). Next, we rescaled values in each D-matrix to be in the 0-1 range and summed the resulting matrices, whereafter PCA and UMAP were performed as above. Information gain was calculated per clustering, by subtracting the weighed entropies of each cluster from the complete entropy. Entropy is defined as -1*Σ*f* · *log2*(*f*), where f is the vector of cluster frequencies.

## Data availability

All relevant data supporting the findings of this study are available within the article and its supplementary information files. All raw sequencing data and processed files will be made available on GEO under accession GSE218476. Any other datasets mentioned in the manuscript were generated using the computational protocols described in the methods.

## Code availability

All relevant code supporting the findings of this study is available on https://github.com/KindLab/MAbID, including the mabidR R-package.

