## Supplementary Table 1 for "Combinatorial single-cell profiling of all major chromatin types with MAbID"

| Primary antibodies used in combination with secondary antibody-DNA conjugates | Target | Host | Type | Clonality | Vendor | Catalog number | LOT numbers | Optimal working concentration |
|---|---|---|---|---|---|---|---|---|
| Anti-Lamin B1 antibody - Nuclear Envelope Marker | Lamin B1 | Rabbit | Primary | Polyclonal | Abcam | ab16048 | GR3398319-7, GR3369248-1 | 10 µg/mL |
| Histone H3K9me2 antibody (pAb) | H3K9me2 | Rabbit | Primary | Polyclonal | Active Motif | 39041 | 39239, 34718002 | 1 in 100 dilution |
| H3K9me3 Recombinant Rabbit Monoclonal Antibody (RM389) | H3K9me3 | Rabbit | Primary | Monoclonal | Invitrogen | MA5-33395 | WI3388337, WH3388337 | 5 µg/mL |
| Tri-methyl-histone-H3 (Lys27) Rabbit mAb | H3K27me3 | Rabbit | Primary | Polyclonal | Cell Signaling Technologies | 9733S | 16,19 | 1 in 200 dilution |
| Recombinant Anti-Histone H3 (tri methyl K27) antibody [EPR18607] - BSA and Azide free | H3K27me3 | Rabbit | Primary | Monoclonal | Abcam | ab222481 | GR3256223-6, GR3256223-1 | 5 µg/mL |
| Anti-Trimethyl-Histone H3 (Lys36) antibody, clone RM155 | H3K36me3 | Rabbit | Primary | Monoclonal | RevMab | 31-1051-00 | T-04-02948 | 0.5-1 µg/mL |
| Histone H3 trimethyl K36 antibody | H3K36me3 | Mouse | Primary | Monoclonal | In-house by Hiroshi Kimura | CM333 | - | 2 µg/mL |
| H3K4me3 Monoclonal Antibody (G.532.8) | H3K4me3 | Rabbit | Primary | Monoclonal | Invitrogen | MA5-11199 | WG3341041, WH334779 | 121-242 ng/mL |
| Histone H3 (mono methyl K4) antibody | H3K4me1 | Rabbit | Primary | Polyclonal | Abcam | ab8895 | GR3402097-1 | 10 µg/mL |
| Recombinant Anti-Histone H3 (acetyl K27) antibody [EP16602] | H3K27ac | Rabbit | Primary | Monoclonal | Abcam | ab177178 | GR3202987-6, GR3202987-19 | 7 µg/mL |
| Recombinant Anti-RNA polymerase II CTD repeat YSPTSPS (phospho S5) antibody [3E8] | Pol II CTD Ser5P | Rat | Primary | Monoclonal | Abcam | ab252852 | GR3302510-1, GR33352497-1 | 5 µg/mL |
| Histone H3 Antibody | Histone H3 | Sheep | Primary | Polyclonal | Novus Biologicals | NB100-747 | p60919 | 25 µg/mL |

| Secondary antibody-DNA conjugates | Target | Host | Type | Clonality | Vendor | Catalog number | LOT numbers | Optimal working concentration |
|---|---|---|---|---|---|---|---|---|
| AffiniPure Goat Anti-Rabbit IgG (H+L) | Rabbit IgG | Goat | Secondary | Polyclonal | Jackson ImmunoResearch | 111-005-144 | 147466 | 2 µg/mL |
| AffiniPure Donkey Anti-Rabbit IgG (H+L) | Rabbit IgG | Donkey | Secondary | Polyclonal | Jackson ImmunoResearch | 711-005-152 | 156033 | 2 µg/mL |
| AffiniPure Donkey Anti-Mouse IgG (H+L) | Mouse IgG | Donkey | Secondary | Polyclonal | Jackson ImmunoResearch | 715-005-150 | 155934 | 2 µg/mL |
| AffiniPure Donkey Anti-Rat IgG (H+L) | Rat IgG | Donkey | Secondary | Polyclonal | Jackson ImmunoResearch | 712-005-150 | 154663 | 2 µg/mL |
| AffiniPure Donkey Anti-Sheep IgG (H+L) | Sheep IgG | Donkey | Secondary | Polyclonal | Jackson ImmunoResearch | 713-005-147 | 150929 | 2 µg/mL |

| Primary antibody-DNA conjugates | Target | Host | Type | Clonality | Vendor | Catalog number | LOT numbers | Optimal working concentration |
|---|---|---|---|---|---|---|---|---|
| Anti-Lamin B1 antibody - Nuclear Envelope Marker | Lamin B1 | Rabbit | Primary | Polyclonal | Abcam | ab16048 | GR3398319-I, GR3417466-I | 20-30 µg/mL |
| H3K9me3 Recombinant Rabbit Monoclonal Antibody (RM389) | H3K9me3 | Rabbit | Primary | Monoclonal | Invitrogen | MA5-33395 | WI3388337, XC3525394 | 10-15 µg/mL |
| Recombinant Anti-Histone H3 (tri methyl K27) antibody [EPR18607] - BSA and Azide free | H3K27me3 | Rabbit | Primary | Monoclonal | Abcam | ab222481 | GR3256223-6 | 20 µg/mL |
| Anti-Trimethyl-Histone H3 (Lys36) antibody, clone RM155 | H3K36me3 | Rabbit | Primary | Monoclonal | RevMab | 31-1051-00 | T-04-02948, T-04-029-48 | 10-20 µg/mL |
| H3K4me3 Monoclonal Antibody (G.532.8) | H3K4me3 | Rabbit | Primary | Monoclonal | Invitrogen | MA5-11199 | WH3347791, WG3341041, WI3395412, XA3475081 | 10 µg/mL |
| Histone H3 (mono methyl K4) antibody | H3K4me1 | Rabbit | Primary | Polyclonal | Abcam | ab8895 | GR3402097-I, GR3426435-2 | 10-15 µg/mL |
| Recombinant Anti-Histone H3 (acetyl K27) antibody [EP16602] | H3K27ac | Rabbit | Primary | Monoclonal | Abcam | ab177178 | GR3202987-6, GR3202987-19 | 20-30 µg/mL |
