## Supplementary Table 2 for "Combinatorial single-cell profiling of all major chromatin types with MAbID"

| Name | ABBC barcode | Sequence (5' - 3') | Restriction site |
|---|---|---|---|
| ABID_ABBC_001_top | CTTCAA | /5AzideN/GTTGGAGTTGAAACGTTCTAATATTCCAATCAGCTTCAACGTGCACCACCGCAGGGCGGCCGCCTTCAACATATGCTAGCATC | NdeI |
| ABID_ABBC_002_top | AGCCAT | /5AzideN/GTTGGAGTTGAAACGTTCTAATATTCCAATCAGCTTCAACGTGCACCACCGCAGGGCGGCCGCAGCCATCATATGCTAGCATC | NdeI |
| ABID_ABBC_003_top | ACACGA | /5AzideN/GTTGGAGTTGAAACGTTCTAATATTCCAATCAGCTTCAACGTGCACCACCGCAGGGCGGCCGCACACGACATATGCTACGTAC | NdeI |
| ABID_ABBC_004_top | GACCTC | /5AzideN/GTTGGAGTTGAAACGTTCTAATATTCCAATCAGCTTCAACGTGCACCACCGCAGGGCGGCCGCGACCTCCATATGCTACGTAC | NdeI |
| ABID_ABBC_005_top | TGGACA | /5AzideN/GTTGGAGTTGAAACGTTCTAATATTCCAATCAGCTTCAACGTGCACCACCGCAGGGCGGCCGCTGGACACATATGCTACGTAC | NdeI |
| ABID_ABBC_006_top | AAGTCC | /5AzideN/GTTGGAGTTGAAACGTTCTAATATTCCAATCAGCTTCAACGTGCACCACCGCAGGGCGGCCGCAAGTCCCATATGCTACGTAC | NdeI |
| ABID_ABBC_007_top | GAAGAA | /5AzideN/GTTGGAGTTGAAACGTTCTAATATTCCAATCAGCTTCAACGTGCACCACCGCAGGGCGGCCGCGAAGAACATATGCTACGTAC | NdeI |
| ABID_ABBC_011_top | TTCGCC | /5AzideN/GTTGGAGTTGAAACGTTCTAATATTCCAATCAGCTTCAACGTGCACCACCGCAGGGCGGCCGCTTCGCCAGATCTCTACGTAC | BglII |
| ABID_ABBC_012_top | ATGGTT | /5AzideN/GTTGGAGTTGAAACGTTCTAATATTCCAATCAGCTTCAACGTGCACCACCGCAGGGCGGCCGCATGGTTAGATCTCTACGTAC | BglII |
| ABID_ABBC_013_top | GTATGC | /5AzideN/GTTGGAGTTGAAACGTTCTAATATTCCAATCAGCTTCAACGTGCACCACCGCAGGGCGGCCGCGTATGCAGATCTCTACGTAC | BglII |
| ABID_ABBC_014_top | CCGAAT | /5AzideN/GTTGGAGTTGAAACGTTCTAATATTCCAATCAGCTTCAACGTGCACCACCGCAGGGCGGCCGCCCGAATAGATCTCTACGTAC | BglII |
| ABID_ABBC_015_top | TATCCA | /5AzideN/GTTGGAGTTGAAACGTTCTAATATTCCAATCAGCTTCAACGTGCACCACCGCAGGGCGGCCGCTATCCAAGATCTCTACGTAC | BglII |
| **Name** | **ABBC barcode** | **Sequence (5' - 3')** | **Restriction site** |
| ABID_ABBC_001_bot | TTGAAG | GATGCTAGCATATGTTGAAGGCGGCCGCCCTGCGGTGGTGCACGTTGAAGCTGATTGGAATATTAGAACGTTTCAACTCCAAC | NdeI |
| ABID_ABBC_002_bot | ATGGCT | GATGCTAGCATATGATGGCTGCGGCCGCCCTGCGGTGGTGCACGTTGAAGCTGATTGGAATATTAGAACGTTTCAACTCCAAC | NdeI |
| ABID_ABBC_003_bot | TCGTGT | GTACGTAGCATATGTCGTGTGCGGCCGCCCTGCGGTGGTGCACGTTGAAGCTGATTGGAATATTAGAACGTTTCAACTCCAAC | NdeI |
| ABID_ABBC_004_bot | GAGGTC | GTACGTAGCATATGGAGGTCGCGGCCGCCCTGCGGTGGTGCACGTTGAAGCTGATTGGAATATTAGAACGTTTCAACTCCAAC | NdeI |
| ABID_ABBC_005_bot | TGTCCA | GTACGTAGCATATGTGTCCAGCGGCCGCCCTGCGGTGGTGCACGTTGAAGCTGATTGGAATATTAGAACGTTTCAACTCCAAC | NdeI |
| ABID_ABBC_006_bot | GGACTT | GTACGTAGCATATGGGACTTGCGGCCGCCCTGCGGTGGTGCACGTTGAAGCTGATTGGAATATTAGAACGTTTCAACTCCAAC | NdeI |
| ABID_ABBC_007_bot | TTCTTC | GTACGTAGCATATGTTCTTCGCGGCCGCCCTGCGGTGGTGCACGTTGAAGCTGATTGGAATATTAGAACGTTTCAACTCCAAC | NdeI |
| ABID_ABBC_011_bot | GGCGAA | GTACGTAGAG/iN6Me-dA/TCTGGCGAAGCGGCCGCCCTGCGGTGGTGCACGTTGAAGCTGATTGGAATATTAGAACGTTTCAACTCCAAC | BglII |
| ABID_ABBC_012_bot | AACCAT | GTACGTAGAG/iN6Me-dA/TCTAACCATGCGGCCGCCCTGCGGTGGTGCACGTTGAAGCTGATTGGAATATTAGAACGTTTCAACTCCAAC | BglII |
| ABID_ABBC_013_bot | GCATAC | GTACGTAGAG/iN6Me-dA/TCTGCATACGCGGCCGCCCTGCGGTGGTGCACGTTGAAGCTGATTGGAATATTAGAACGTTTCAACTCCAAC | BglII |
| ABID_ABBC_014_bot | ATTCGG | GTACGTAGAG/iN6Me-dA/TCTATTCGGGCGGCCGCCCTGCGGTGGTGCACGTTGAAGCTGATTGGAATATTAGAACGTTTCAACTCCAAC | BglII |
| ABID_ABBC_015_bot | TGGATA | GTACGTAGAG/iN6Me-dA/TCTTGGATAGCGGCCGCCCTGCGGTGGTGCACGTTGAAGCTGATTGGAATATTAGAACGTTTCAACTCCAAC | BglII |
