## Supplementary Table 3 for "Combinatorial single-cell profiling of all major chromatin types with MAbID"

| Name | SBC barcode | Sequence (5' - 3') |
|---|---|---|
| ABID_SBC_top_001 | AGGCCATT | GGTGATCCGGTAATACGACTCACTATAGGGGTTCAGAGTTCTACAGTCCGACGATCNNNAGGCNNNCATTAG |
| ABID_SBC_top_002 | TGAGGCTA | GGTGATCCGGTAATACGACTCACTATAGGGGTTCAGAGTTCTACAGTCCGACGATCNNNTGAGNNNGCTAAG |
| ABID_SBC_top_003 | GAGAGCTA | GGTGATCCGGTAATACGACTCACTATAGGGGTTCAGAGTTCTACAGTCCGACGATCNNNGAGANNNGCTAAG |
| ABID_SBC_top_004 | GTTCCGAG | GGTGATCCGGTAATACGACTCACTATAGGGGTTCAGAGTTCTACAGTCCGACGATCNNNGTTCNNNCGAGAG |
| ABID_SBC_top_005 | GTTGTGAC | GGTGATCCGGTAATACGACTCACTATAGGGGTTCAGAGTTCTACAGTCCGACGATCNNNGTTGNNNTGACAG |
| ABID_SBC_top_006 | TACAGACT | GGTGATCCGGTAATACGACTCACTATAGGGGTTCAGAGTTCTACAGTCCGACGATCNNNTACANNNGACTAG |
| ABID_SBC_top_007 | CAGCAGCT | GGTGATCCGGTAATACGACTCACTATAGGGGTTCAGAGTTCTACAGTCCGACGATCNNNCAGCNNNAGCTAG |
| ABID_SBC_top_008 | CTGAGTCA | GGTGATCCGGTAATACGACTCACTATAGGGGTTCAGAGTTCTACAGTCCGACGATCNNNCTGANNNGTCAAG |
| ABID_SBC_top_009 | GATGCTGC | GGTGATCCGGTAATACGACTCACTATAGGGGTTCAGAGTTCTACAGTCCGACGATCNNNGATGNNNCTGCAG |
| ABID_SBC_top_010 | ATGCGCAG | GGTGATCCGGTAATACGACTCACTATAGGGGTTCAGAGTTCTACAGTCCGACGATCNNNATGCNNNGCAGAG |
| ABID_SBC_top_011 | ACCTCTTG | GGTGATCCGGTAATACGACTCACTATAGGGGTTCAGAGTTCTACAGTCCGACGATCNNNACCTNNNCTTGAG |
| ABID_SBC_top_012 | TATCCGGT | GGTGATCCGGTAATACGACTCACTATAGGGGTTCAGAGTTCTACAGTCCGACGATCNNNTATCNNNCGGTAG |
| ABID_SBC_top_013 | TAACACTG | GGTGATCCGGTAATACGACTCACTATAGGGGTTCAGAGTTCTACAGTCCGACGATCNNNTAACNNNACTGAG |
| ABID_SBC_top_014 | GTTGAGTT | GGTGATCCGGTAATACGACTCACTATAGGGGTTCAGAGTTCTACAGTCCGACGATCNNNGTTGNNNAGTTAG |
| ABID_SBC_top_015 | TCCTGTCC | GGTGATCCGGTAATACGACTCACTATAGGGGTTCAGAGTTCTACAGTCCGACGATCNNNTCCTNNNGTCCAG |
| ABID_SBC_top_016 | CTTGGAAG | GGTGATCCGGTAATACGACTCACTATAGGGGTTCAGAGTTCTACAGTCCGACGATCNNNCTTGNNNGAAGAG |
| ABID_SBC_top_017 | CTGTATTG | GGTGATCCGGTAATACGACTCACTATAGGGGTTCAGAGTTCTACAGTCCGACGATCNNNCTGTNNNATTGAG |
| ABID_SBC_top_018 | CGTACTAT | GGTGATCCGGTAATACGACTCACTATAGGGGTTCAGAGTTCTACAGTCCGACGATCNNNCGTANNNCTATAG |
| ABID_SBC_top_019 | GTATCTCC | GGTGATCCGGTAATACGACTCACTATAGGGGTTCAGAGTTCTACAGTCCGACGATCNNNGTATNNNCTCCAG |
| ABID_SBC_top_020 | TGGTCTAA | GGTGATCCGGTAATACGACTCACTATAGGGGTTCAGAGTTCTACAGTCCGACGATCNNNTGGTNNNCTAAAG |
| ABID_SBC_top_021 | CATCGTCC | GGTGATCCGGTAATACGACTCACTATAGGGGTTCAGAGTTCTACAGTCCGACGATCNNNCATCNNNGTCCAG |
| ABID_SBC_top_022 | CTCTATCT | GGTGATCCGGTAATACGACTCACTATAGGGGTTCAGAGTTCTACAGTCCGACGATCNNNCTCTNNNATCTAG |
| ABID_SBC_top_023 | ACTACTGT | GGTGATCCGGTAATACGACTCACTATAGGGGTTCAGAGTTCTACAGTCCGACGATCNNNACTANNNCTGTAG |
| ABID_SBC_top_024 | GACGGTAG | GGTGATCCGGTAATACGACTCACTATAGGGGTTCAGAGTTCTACAGTCCGACGATCNNNGACGNNNGTAGAG |
| ABID_SBC_top_025 | TGATGTCT | GGTGATCCGGTAATACGACTCACTATAGGGGTTCAGAGTTCTACAGTCCGACGATCNNNTGATNNNGTCTAG |
| ABID_SBC_top_026 | TAACGTGT | GGTGATCCGGTAATACGACTCACTATAGGGGTTCAGAGTTCTACAGTCCGACGATCNNNTAACNNNGTGTAG |
| ABID_SBC_top_027 | CGATTCCG | GGTGATCCGGTAATACGACTCACTATAGGGGTTCAGAGTTCTACAGTCCGACGATCNNNCGATNNNTCCGAG |
| ABID_SBC_top_028 | GCTCGTCT | GGTGATCCGGTAATACGACTCACTATAGGGGTTCAGAGTTCTACAGTCCGACGATCNNNGCTCNNNGTCTAG |
| ABID_SBC_top_029 | TGGTGCTC | GGTGATCCGGTAATACGACTCACTATAGGGGTTCAGAGTTCTACAGTCCGACGATCNNNTGGTNNNGCTCAG |
| ABID_SBC_top_030 | CAGTTCGA | GGTGATCCGGTAATACGACTCACTATAGGGGTTCAGAGTTCTACAGTCCGACGATCNNNCAGTNNNTCGAAG |
| ABID_SBC_top_031 | ACTCCGGA | GGTGATCCGGTAATACGACTCACTATAGGGGTTCAGAGTTCTACAGTCCGACGATCNNNACTCNNNCGGAAG |
| ABID_SBC_top_032 | CGCAAGCA | GGTGATCCGGTAATACGACTCACTATAGGGGTTCAGAGTTCTACAGTCCGACGATCNNNCGCANNNAGCAAG |

| | | |
|---|---|---|
| ABID_SBC_top_033 | ATCGGATC | GGTGATCCGGTAATACGACTCACTATAGGGGTTCAGAGTTCTACAGTCCGACGATCNNNATCGNNNGATCAG |
| ABID_SBC_top_034 | ACCGGTAT | GGTGATCCGGTAATACGACTCACTATAGGGGTTCAGAGTTCTACAGTCCGACGATCNNNACCGNNNGTATAG |
| ABID_SBC_top_035 | GAAGGAAT | GGTGATCCGGTAATACGACTCACTATAGGGGTTCAGAGTTCTACAGTCCGACGATCNNNGAAGNNNGAATAG |
| ABID_SBC_top_036 | ACTGGATG | GGTGATCCGGTAATACGACTCACTATAGGGGTTCAGAGTTCTACAGTCCGACGATCNNNACTGNNNGATGAG |
| ABID_SBC_top_037 | CGGATGAA | GGTGATCCGGTAATACGACTCACTATAGGGGTTCAGAGTTCTACAGTCCGACGATCNNNCGGANNNTGAAAG |
| ABID_SBC_top_038 | GTCAACGT | GGTGATCCGGTAATACGACTCACTATAGGGGTTCAGAGTTCTACAGTCCGACGATCNNNGTCANNNACGTAG |
| ABID_SBC_top_039 | TGCTAGTC | GGTGATCCGGTAATACGACTCACTATAGGGGTTCAGAGTTCTACAGTCCGACGATCNNNTGCTNNNAGTCAG |
| ABID_SBC_top_040 | TGGCAGAC | GGTGATCCGGTAATACGACTCACTATAGGGGTTCAGAGTTCTACAGTCCGACGATCNNNTGGCNNNAGACAG |
| ABID_SBC_top_041 | CTCGGACA | GGTGATCCGGTAATACGACTCACTATAGGGGTTCAGAGTTCTACAGTCCGACGATCNNNCTCGNNNGACAAG |
| ABID_SBC_top_042 | TCCGTCTC | GGTGATCCGGTAATACGACTCACTATAGGGGTTCAGAGTTCTACAGTCCGACGATCNNNTCCGNNNTCTCAG |
| ABID_SBC_top_043 | CGGTGATG | GGTGATCCGGTAATACGACTCACTATAGGGGTTCAGAGTTCTACAGTCCGACGATCNNNCGGTNNNGATGAG |
| ABID_SBC_top_044 | ACTCACTA | GGTGATCCGGTAATACGACTCACTATAGGGGTTCAGAGTTCTACAGTCCGACGATCNNNACTCNNNACTAAG |
| ABID_SBC_top_045 | TCGATACC | GGTGATCCGGTAATACGACTCACTATAGGGGTTCAGAGTTCTACAGTCCGACGATCNNNTCGANNNTACCAG |
| ABID_SBC_top_046 | TAACCACA | GGTGATCCGGTAATACGACTCACTATAGGGGTTCAGAGTTCTACAGTCCGACGATCNNNTAACNNNCACAAG |
| ABID_SBC_top_047 | GCTAGCTC | GGTGATCCGGTAATACGACTCACTATAGGGGTTCAGAGTTCTACAGTCCGACGATCNNNGCTANNNGCTCAG |
| ABID_SBC_top_048 | GCTACGTT | GGTGATCCGGTAATACGACTCACTATAGGGGTTCAGAGTTCTACAGTCCGACGATCNNNGCTANNNCGTTAG |
| ABID_SBC_top_049 | GAACGTTA | GGTGATCCGGTAATACGACTCACTATAGGGGTTCAGAGTTCTACAGTCCGACGATCNNNGAACNNNGTTAAG |
| ABID_SBC_top_050 | TCCACTAC | GGTGATCCGGTAATACGACTCACTATAGGGGTTCAGAGTTCTACAGTCCGACGATCNNNTCCANNNCTACAG |
| ABID_SBC_top_051 | TAGTCTTG | GGTGATCCGGTAATACGACTCACTATAGGGGTTCAGAGTTCTACAGTCCGACGATCNNNTAGTNNNCTTGAG |
| ABID_SBC_top_052 | CTAGATGA | GGTGATCCGGTAATACGACTCACTATAGGGGTTCAGAGTTCTACAGTCCGACGATCNNNCTAGNNNATGAAG |
| ABID_SBC_top_053 | GTGTGCAT | GGTGATCCGGTAATACGACTCACTATAGGGGTTCAGAGTTCTACAGTCCGACGATCNNNGTGTNNNGCATAG |
| ABID_SBC_top_054 | CGGAATTC | GGTGATCCGGTAATACGACTCACTATAGGGGTTCAGAGTTCTACAGTCCGACGATCNNNCGGANNNATTCAG |
| ABID_SBC_top_055 | ATGACAGA | GGTGATCCGGTAATACGACTCACTATAGGGGTTCAGAGTTCTACAGTCCGACGATCNNNATGANNNCAGAAG |
| ABID_SBC_top_056 | CGACTACC | GGTGATCCGGTAATACGACTCACTATAGGGGTTCAGAGTTCTACAGTCCGACGATCNNNCGACNNNTACCAG |
| ABID_SBC_top_057 | TCGTAGTT | GGTGATCCGGTAATACGACTCACTATAGGGGTTCAGAGTTCTACAGTCCGACGATCNNNTCGTNNNAGTTAG |
| ABID_SBC_top_058 | GATAGAAC | GGTGATCCGGTAATACGACTCACTATAGGGGTTCAGAGTTCTACAGTCCGACGATCNNNGATANNNGAACAG |
| ABID_SBC_top_059 | TAACTGCC | GGTGATCCGGTAATACGACTCACTATAGGGGTTCAGAGTTCTACAGTCCGACGATCNNNTAACNNNTGCCAG |
| ABID_SBC_top_060 | GTTATACG | GGTGATCCGGTAATACGACTCACTATAGGGGTTCAGAGTTCTACAGTCCGACGATCNNNGTTANNNTACGAG |
| ABID_SBC_top_061 | CAAGTCAC | GGTGATCCGGTAATACGACTCACTATAGGGGTTCAGAGTTCTACAGTCCGACGATCNNNCAAGNNNTCACAG |
| ABID_SBC_top_062 | CTTATCAG | GGTGATCCGGTAATACGACTCACTATAGGGGTTCAGAGTTCTACAGTCCGACGATCNNNCTTANNNTCAGAG |
| ABID_SBC_top_063 | GTTAATGC | GGTGATCCGGTAATACGACTCACTATAGGGGTTCAGAGTTCTACAGTCCGACGATCNNNGTTANNNATGCAG |
| ABID_SBC_top_064 | CACGCATG | GGTGATCCGGTAATACGACTCACTATAGGGGTTCAGAGTTCTACAGTCCGACGATCNNNCACGNNNCATGAG |
| ABID_SBC_top_065 | TCTGGTCG | GGTGATCCGGTAATACGACTCACTATAGGGGTTCAGAGTTCTACAGTCCGACGATCNNNTCTGNNNGTCGAG |

| | | |
|---|---|---|
| ABID_SBC_top_066 | TGCTATAG | GGTGATCCGGTAATACGACTCACTATAGGGGTTCAGAGTTCTACAGTCCGACGATCNNNTGCTNNNATAGAG |
| ABID_SBC_top_067 | TCCGCTGT | GGTGATCCGGTAATACGACTCACTATAGGGGTTCAGAGTTCTACAGTCCGACGATCNNNTCCGNNNCTGTAG |
| ABID_SBC_top_068 | CTTACAAC | GGTGATCCGGTAATACGACTCACTATAGGGGTTCAGAGTTCTACAGTCCGACGATCNNNCTTANNNCAACAG |
| ABID_SBC_top_069 | TCTGCGCT | GGTGATCCGGTAATACGACTCACTATAGGGGTTCAGAGTTCTACAGTCCGACGATCNNNTCTGNNNCGCTAG |
| ABID_SBC_top_070 | TGGCGTGA | GGTGATCCGGTAATACGACTCACTATAGGGGTTCAGAGTTCTACAGTCCGACGATCNNNTGGCNNNGTGAAG |
| ABID_SBC_top_071 | ATATCGGC | GGTGATCCGGTAATACGACTCACTATAGGGGTTCAGAGTTCTACAGTCCGACGATCNNNATATNNNCGGCAG |
| ABID_SBC_top_072 | GTAGATTC | GGTGATCCGGTAATACGACTCACTATAGGGGTTCAGAGTTCTACAGTCCGACGATCNNNGTAGNNNATTCAG |
| ABID_SBC_top_073 | AGAGGCAG | GGTGATCCGGTAATACGACTCACTATAGGGGTTCAGAGTTCTACAGTCCGACGATCNNNAGAGNNNGCAGAG |
| ABID_SBC_top_074 | TATCGTTG | GGTGATCCGGTAATACGACTCACTATAGGGGTTCAGAGTTCTACAGTCCGACGATCNNNTATCNNNGTTGAG |
| ABID_SBC_top_075 | GTATCGAA | GGTGATCCGGTAATACGACTCACTATAGGGGTTCAGAGTTCTACAGTCCGACGATCNNNGTATNNNCGAAAG |
| ABID_SBC_top_076 | TGCAGATC | GGTGATCCGGTAATACGACTCACTATAGGGGTTCAGAGTTCTACAGTCCGACGATCNNNTGCANNNGATCAG |
| ABID_SBC_top_077 | TGGAACGA | GGTGATCCGGTAATACGACTCACTATAGGGGTTCAGAGTTCTACAGTCCGACGATCNNNTGGANNNACGAAG |
| ABID_SBC_top_078 | TACTCACC | GGTGATCCGGTAATACGACTCACTATAGGGGTTCAGAGTTCTACAGTCCGACGATCNNNTACTNNNCACCAG |
| ABID_SBC_top_079 | AGAGAGTC | GGTGATCCGGTAATACGACTCACTATAGGGGTTCAGAGTTCTACAGTCCGACGATCNNNAGAGNNNAGTCAG |
| ABID_SBC_top_080 | ACCAGAGC | GGTGATCCGGTAATACGACTCACTATAGGGGTTCAGAGTTCTACAGTCCGACGATCNNNACCANNNGAGCAG |
| ABID_SBC_top_081 | GCATCGGT | GGTGATCCGGTAATACGACTCACTATAGGGGTTCAGAGTTCTACAGTCCGACGATCNNNGCATNNNCGGTAG |
| ABID_SBC_top_082 | TCGCTCTG | GGTGATCCGGTAATACGACTCACTATAGGGGTTCAGAGTTCTACAGTCCGACGATCNNNTCGCNNNTCTGAG |
| ABID_SBC_top_083 | AGCGTCAT | GGTGATCCGGTAATACGACTCACTATAGGGGTTCAGAGTTCTACAGTCCGACGATCNNNAGCGNNNTCATAG |
| ABID_SBC_top_084 | CGTCATTA | GGTGATCCGGTAATACGACTCACTATAGGGGTTCAGAGTTCTACAGTCCGACGATCNNNCGTCNNNATTAAG |
| ABID_SBC_top_085 | TGATTGAG | GGTGATCCGGTAATACGACTCACTATAGGGGTTCAGAGTTCTACAGTCCGACGATCNNNTGATNNNTGAGAG |
| ABID_SBC_top_086 | CGCTTCTA | GGTGATCCGGTAATACGACTCACTATAGGGGTTCAGAGTTCTACAGTCCGACGATCNNNCGCTNNNTCTAAG |
| ABID_SBC_top_087 | TGTAGCGC | GGTGATCCGGTAATACGACTCACTATAGGGGTTCAGAGTTCTACAGTCCGACGATCNNNTGTANNNGCGCAG |
| ABID_SBC_top_088 | GTGTCACG | GGTGATCCGGTAATACGACTCACTATAGGGGTTCAGAGTTCTACAGTCCGACGATCNNNGTGTNNNCACGAG |
| ABID_SBC_top_089 | TGCGGTCA | GGTGATCCGGTAATACGACTCACTATAGGGGTTCAGAGTTCTACAGTCCGACGATCNNNTGCGNNNGTCAAG |
| ABID_SBC_top_090 | CAATCAGC | GGTGATCCGGTAATACGACTCACTATAGGGGTTCAGAGTTCTACAGTCCGACGATCNNNCAATNNNCAGCAG |
| ABID_SBC_top_091 | GTAGCGTG | GGTGATCCGGTAATACGACTCACTATAGGGGTTCAGAGTTCTACAGTCCGACGATCNNNGTAGNNNCGTGAG |
| ABID_SBC_top_092 | CGATAGTG | GGTGATCCGGTAATACGACTCACTATAGGGGTTCAGAGTTCTACAGTCCGACGATCNNNCGATNNNAGTGAG |
| ABID_SBC_top_093 | ACACTCAC | GGTGATCCGGTAATACGACTCACTATAGGGGTTCAGAGTTCTACAGTCCGACGATCNNNACACNNNTCACAG |
| ABID_SBC_top_094 | ACCTTCGC | GGTGATCCGGTAATACGACTCACTATAGGGGTTCAGAGTTCTACAGTCCGACGATCNNNACCTNNNTCGCAG |
| ABID_SBC_top_095 | ATCGTAGA | GGTGATCCGGTAATACGACTCACTATAGGGGTTCAGAGTTCTACAGTCCGACGATCNNNATCGNNNTAGAAG |
| ABID_SBC_top_096 | CACAGTCG | GGTGATCCGGTAATACGACTCACTATAGGGGTTCAGAGTTCTACAGTCCGACGATCNNNCACANNNGTCGAG |
| ABID_SBC_top_097 | TCCAGCCA | GGTGATCCGGTAATACGACTCACTATAGGGGTTCAGAGTTCTACAGTCCGACGATCNNNTCCANNNGCCAAG |
| ABID_SBC_top_098 | TGCTTACA | GGTGATCCGGTAATACGACTCACTATAGGGGTTCAGAGTTCTACAGTCCGACGATCNNNTGCTNNNTACAAG |

| | | |
|---|---|---|
| ABID_SBC_top_099 | TGGCTGCA | GGTGATCCGGTAATACGACTCACTATAGGGGTTCAGAGTTCTACAGTCCGACGATCNNNTGGCNNNTGCAAG |
| ABID_SBC_top_100 | TGCGTCGA | GGTGATCCGGTAATACGACTCACTATAGGGGTTCAGAGTTCTACAGTCCGACGATCNNNTGCGNNNTCGAAG |
| ABID_SBC_top_101 | AGCAGTAC | GGTGATCCGGTAATACGACTCACTATAGGGGTTCAGAGTTCTACAGTCCGACGATCNNNAGCANNNGTACAG |
| ABID_SBC_top_102 | GCCTAGGA | GGTGATCCGGTAATACGACTCACTATAGGGGTTCAGAGTTCTACAGTCCGACGATCNNNGCCTNNNAGGAAG |
| ABID_SBC_top_103 | TAAGGTCC | GGTGATCCGGTAATACGACTCACTATAGGGGTTCAGAGTTCTACAGTCCGACGATCNNNTAAGNNNGTCCAG |
| ABID_SBC_top_104 | TGTAGACA | GGTGATCCGGTAATACGACTCACTATAGGGGTTCAGAGTTCTACAGTCCGACGATCNNNTGTANNNGACAAG |
| ABID_SBC_top_105 | CGCAGTGA | GGTGATCCGGTAATACGACTCACTATAGGGGTTCAGAGTTCTACAGTCCGACGATCNNNCGCANNNGTGAAG |
| ABID_SBC_top_106 | TCTAAGCC | GGTGATCCGGTAATACGACTCACTATAGGGGTTCAGAGTTCTACAGTCCGACGATCNNNTCTANNNAGCCAG |
| ABID_SBC_top_107 | TGACGACG | GGTGATCCGGTAATACGACTCACTATAGGGGTTCAGAGTTCTACAGTCCGACGATCNNNTGACNNNGACGAG |
| ABID_SBC_top_108 | CTAGTAGT | GGTGATCCGGTAATACGACTCACTATAGGGGTTCAGAGTTCTACAGTCCGACGATCNNNCTAGNNNTAGTAG |
| ABID_SBC_top_109 | CATAAGGC | GGTGATCCGGTAATACGACTCACTATAGGGGTTCAGAGTTCTACAGTCCGACGATCNNNCATANNNAGGCAG |
| ABID_SBC_top_110 | AGACATAC | GGTGATCCGGTAATACGACTCACTATAGGGGTTCAGAGTTCTACAGTCCGACGATCNNNAGACNNNATACAG |
| ABID_SBC_top_111 | CGACGAAT | GGTGATCCGGTAATACGACTCACTATAGGGGTTCAGAGTTCTACAGTCCGACGATCNNNCGACNNNGAATAG |
| ABID_SBC_top_112 | GCAGGACA | GGTGATCCGGTAATACGACTCACTATAGGGGTTCAGAGTTCTACAGTCCGACGATCNNNGCAGNNNGACAAG |
| ABID_SBC_top_113 | CATGACCG | GGTGATCCGGTAATACGACTCACTATAGGGGTTCAGAGTTCTACAGTCCGACGATCNNNCATGNNNACCGAG |
| ABID_SBC_top_114 | CTACGTCG | GGTGATCCGGTAATACGACTCACTATAGGGGTTCAGAGTTCTACAGTCCGACGATCNNNCTACNNNGTCGAG |
| ABID_SBC_top_115 | TCGAGATT | GGTGATCCGGTAATACGACTCACTATAGGGGTTCAGAGTTCTACAGTCCGACGATCNNNTCGANNNGATTAG |
| ABID_SBC_top_116 | GTATTCTG | GGTGATCCGGTAATACGACTCACTATAGGGGTTCAGAGTTCTACAGTCCGACGATCNNNGTATNNNTCTGAG |
| ABID_SBC_top_117 | CGTCTGTC | GGTGATCCGGTAATACGACTCACTATAGGGGTTCAGAGTTCTACAGTCCGACGATCNNNCGTCNNNTGTCAG |
| ABID_SBC_top_118 | CGTCACAG | GGTGATCCGGTAATACGACTCACTATAGGGGTTCAGAGTTCTACAGTCCGACGATCNNNCGTCNNNACAGAG |
| ABID_SBC_top_119 | TCGCACCT | GGTGATCCGGTAATACGACTCACTATAGGGGTTCAGAGTTCTACAGTCCGACGATCNNNTCGCNNNACCTAG |
| ABID_SBC_top_120 | CGACCTTG | GGTGATCCGGTAATACGACTCACTATAGGGGTTCAGAGTTCTACAGTCCGACGATCNNNCGACNNNCTTGAG |
| ABID_SBC_top_121 | ATGTTGGA | GGTGATCCGGTAATACGACTCACTATAGGGGTTCAGAGTTCTACAGTCCGACGATCNNNATGTNNNTGGAAG |
| ABID_SBC_top_122 | GTCGGTCT | GGTGATCCGGTAATACGACTCACTATAGGGGTTCAGAGTTCTACAGTCCGACGATCNNNGTCGNNNGTCTAG |
| ABID_SBC_top_123 | CGATGTTA | GGTGATCCGGTAATACGACTCACTATAGGGGTTCAGAGTTCTACAGTCCGACGATCNNNCGATNNNGTTAAG |
| ABID_SBC_top_124 | TGCTCATT | GGTGATCCGGTAATACGACTCACTATAGGGGTTCAGAGTTCTACAGTCCGACGATCNNNTGCTNNNCATTAG |
| ABID_SBC_top_125 | GCCTATCG | GGTGATCCGGTAATACGACTCACTATAGGGGTTCAGAGTTCTACAGTCCGACGATCNNNGCCTNNNATCGAG |
| ABID_SBC_top_126 | ACTCTAAG | GGTGATCCGGTAATACGACTCACTATAGGGGTTCAGAGTTCTACAGTCCGACGATCNNNACTCNNNTAAGAG |
| ABID_SBC_top_127 | GAAGTGGT | GGTGATCCGGTAATACGACTCACTATAGGGGTTCAGAGTTCTACAGTCCGACGATCNNNGAAGNNNTGGTAG |
| ABID_SBC_top_128 | GAACAGAA | GGTGATCCGGTAATACGACTCACTATAGGGGTTCAGAGTTCTACAGTCCGACGATCNNNGAACNNNAGAAAG |
| ABID_SBC_top_129 | AGGATCTA | GGTGATCCGGTAATACGACTCACTATAGGGGTTCAGAGTTCTACAGTCCGACGATCNNNAGGANNNTCTAAG |
| ABID_SBC_top_130 | TATCCATC | GGTGATCCGGTAATACGACTCACTATAGGGGTTCAGAGTTCTACAGTCCGACGATCNNNTATCNNNCATCAG |
| ABID_SBC_top_131 | GTTCCTTA | GGTGATCCGGTAATACGACTCACTATAGGGGTTCAGAGTTCTACAGTCCGACGATCNNNGTTCNNNCTTAAG |

| | | |
|---|---|---|
| ABID_SBC_top_132 | ACGCAGTC | GGTGATCCGGTAATACGACTCACTATAGGGGTTCAGAGTTCTACAGTCCGACGATCNNNACGCNNNAGTCAG |
| ABID_SBC_top_133 | ATTGGTGT | GGTGATCCGGTAATACGACTCACTATAGGGGTTCAGAGTTCTACAGTCCGACGATCNNNATTGNNNGTGTAG |
| ABID_SBC_top_134 | ACTACGCG | GGTGATCCGGTAATACGACTCACTATAGGGGTTCAGAGTTCTACAGTCCGACGATCNNNACTANNNCGCGAG |
| ABID_SBC_top_135 | GCCAGCAT | GGTGATCCGGTAATACGACTCACTATAGGGGTTCAGAGTTCTACAGTCCGACGATCNNNGCCANNNGCATAG |
| ABID_SBC_top_136 | CTCGCTAG | GGTGATCCGGTAATACGACTCACTATAGGGGTTCAGAGTTCTACAGTCCGACGATCNNNCTCGNNNCTAGAG |
| ABID_SBC_top_137 | CATGGACT | GGTGATCCGGTAATACGACTCACTATAGGGGTTCAGAGTTCTACAGTCCGACGATCNNNCATGNNNGACTAG |
| ABID_SBC_top_138 | ATCAAGTC | GGTGATCCGGTAATACGACTCACTATAGGGGTTCAGAGTTCTACAGTCCGACGATCNNNATCANNNAGTCAG |
| ABID_SBC_top_139 | AGTCCGCT | GGTGATCCGGTAATACGACTCACTATAGGGGTTCAGAGTTCTACAGTCCGACGATCNNNAGTCNNNCGCTAG |
| ABID_SBC_top_140 | CTACACTT | GGTGATCCGGTAATACGACTCACTATAGGGGTTCAGAGTTCTACAGTCCGACGATCNNNCTACNNNACTTAG |
| ABID_SBC_top_141 | CTCTCGTA | GGTGATCCGGTAATACGACTCACTATAGGGGTTCAGAGTTCTACAGTCCGACGATCNNNCTCTNNNCGTAAG |
| ABID_SBC_top_142 | CTGCTGTT | GGTGATCCGGTAATACGACTCACTATAGGGGTTCAGAGTTCTACAGTCCGACGATCNNNCTGCNNNTGTTAG |
| ABID_SBC_top_143 | TACATAGC | GGTGATCCGGTAATACGACTCACTATAGGGGTTCAGAGTTCTACAGTCCGACGATCNNNTACANNNTAGCAG |
| ABID_SBC_top_144 | CAATGTAG | GGTGATCCGGTAATACGACTCACTATAGGGGTTCAGAGTTCTACAGTCCGACGATCNNNCAATNNNGTAGAG |
| ABID_SBC_top_145 | CTTAGTTC | GGTGATCCGGTAATACGACTCACTATAGGGGTTCAGAGTTCTACAGTCCGACGATCNNNCTTANNNGTTCAG |
| ABID_SBC_top_146 | CACTGCCA | GGTGATCCGGTAATACGACTCACTATAGGGGTTCAGAGTTCTACAGTCCGACGATCNNNCACTNNNGCCAAG |
| ABID_SBC_top_147 | ATGCCGTG | GGTGATCCGGTAATACGACTCACTATAGGGGTTCAGAGTTCTACAGTCCGACGATCNNNATGCNNNCGTGAG |
| ABID_SBC_top_148 | AGTGTAAC | GGTGATCCGGTAATACGACTCACTATAGGGGTTCAGAGTTCTACAGTCCGACGATCNNNAGTGNNNTAACAG |
| ABID_SBC_top_149 | ATTGAGCA | GGTGATCCGGTAATACGACTCACTATAGGGGTTCAGAGTTCTACAGTCCGACGATCNNNATTGNNNAGCAAG |
| ABID_SBC_top_150 | TAGTGACG | GGTGATCCGGTAATACGACTCACTATAGGGGTTCAGAGTTCTACAGTCCGACGATCNNNTAGTNNNGACGAG |
| ABID_SBC_top_151 | TGAGCATG | GGTGATCCGGTAATACGACTCACTATAGGGGTTCAGAGTTCTACAGTCCGACGATCNNNTGAGNNNCATGAG |
| ABID_SBC_top_152 | CGCTAGAT | GGTGATCCGGTAATACGACTCACTATAGGGGTTCAGAGTTCTACAGTCCGACGATCNNNCGCTNNNAGATAG |
| ABID_SBC_top_153 | TCCGATTA | GGTGATCCGGTAATACGACTCACTATAGGGGTTCAGAGTTCTACAGTCCGACGATCNNNTCCGNNNATTAAG |
| ABID_SBC_top_154 | TGGCCAAG | GGTGATCCGGTAATACGACTCACTATAGGGGTTCAGAGTTCTACAGTCCGACGATCNNNTGGCNNNCAAGAG |
| ABID_SBC_top_155 | TCGACGGT | GGTGATCCGGTAATACGACTCACTATAGGGGTTCAGAGTTCTACAGTCCGACGATCNNNTCGANNNCGGTAG |
| ABID_SBC_top_156 | ACCGTACT | GGTGATCCGGTAATACGACTCACTATAGGGGTTCAGAGTTCTACAGTCCGACGATCNNNACCGNNNTACTAG |
| ABID_SBC_top_157 | CATCTAAC | GGTGATCCGGTAATACGACTCACTATAGGGGTTCAGAGTTCTACAGTCCGACGATCNNNCATCNNNTAACAG |
| ABID_SBC_top_158 | ATGAATCC | GGTGATCCGGTAATACGACTCACTATAGGGGTTCAGAGTTCTACAGTCCGACGATCNNNATGANNNATCCAG |
| ABID_SBC_top_159 | TCGTGAGA | GGTGATCCGGTAATACGACTCACTATAGGGGTTCAGAGTTCTACAGTCCGACGATCNNNTCGTNNNGAGAAG |
| ABID_SBC_top_160 | GTGAAGAT | GGTGATCCGGTAATACGACTCACTATAGGGGTTCAGAGTTCTACAGTCCGACGATCNNNGTGANNNAGATAG |
| ABID_SBC_top_161 | GAGCGAAG | GGTGATCCGGTAATACGACTCACTATAGGGGTTCAGAGTTCTACAGTCCGACGATCNNNGAGCNNNGAAGAG |
| ABID_SBC_top_162 | AGGTCTCT | GGTGATCCGGTAATACGACTCACTATAGGGGTTCAGAGTTCTACAGTCCGACGATCNNNAGGTNNNCTCTAG |
| ABID_SBC_top_163 | GCACAGTT | GGTGATCCGGTAATACGACTCACTATAGGGGTTCAGAGTTCTACAGTCCGACGATCNNNGCACNNNAGTTAG |
| ABID_SBC_top_164 | GTTGCTAT | GGTGATCCGGTAATACGACTCACTATAGGGGTTCAGAGTTCTACAGTCCGACGATCNNNGTTGNNNCTATAG |

| | | |
|---|---|---|
| ABID_SBC_top_165 | TGCGTGTG | GGTGATCCGGTAATACGACTCACTATAGGGGTTCAGAGTTCTACAGTCCGACGATCNNNTGCGNNNTGTGAG |
| ABID_SBC_top_166 | GTACTGTC | GGTGATCCGGTAATACGACTCACTATAGGGGTTCAGAGTTCTACAGTCCGACGATCNNNGTACNNNTGTCAG |
| ABID_SBC_top_167 | GTGCAGGA | GGTGATCCGGTAATACGACTCACTATAGGGGTTCAGAGTTCTACAGTCCGACGATCNNNGTGCNNNAGGAAG |
| ABID_SBC_top_168 | GTATAGCG | GGTGATCCGGTAATACGACTCACTATAGGGGTTCAGAGTTCTACAGTCCGACGATCNNNGTATNNNAGCGAG |
| ABID_SBC_top_169 | ATTGCTTC | GGTGATCCGGTAATACGACTCACTATAGGGGTTCAGAGTTCTACAGTCCGACGATCNNNATTGNNNCTTCAG |
| ABID_SBC_top_170 | GCTCATGA | GGTGATCCGGTAATACGACTCACTATAGGGGTTCAGAGTTCTACAGTCCGACGATCNNNGCTCNNNATGAAG |
| ABID_SBC_top_171 | GTCGTATT | GGTGATCCGGTAATACGACTCACTATAGGGGTTCAGAGTTCTACAGTCCGACGATCNNNGTCGNNNTATTAG |
| ABID_SBC_top_172 | ACGCATAG | GGTGATCCGGTAATACGACTCACTATAGGGGTTCAGAGTTCTACAGTCCGACGATCNNNACGCNNNATAGAG |
| ABID_SBC_top_173 | GCATTAGA | GGTGATCCGGTAATACGACTCACTATAGGGGTTCAGAGTTCTACAGTCCGACGATCNNNGCATNNNTAGAAG |
| ABID_SBC_top_174 | TCTCTGTA | GGTGATCCGGTAATACGACTCACTATAGGGGTTCAGAGTTCTACAGTCCGACGATCNNNTCTCNNNTGTAAG |
| ABID_SBC_top_175 | TGTCTACT | GGTGATCCGGTAATACGACTCACTATAGGGGTTCAGAGTTCTACAGTCCGACGATCNNNTGTCNNNTACTAG |
| ABID_SBC_top_176 | CGTCTCGA | GGTGATCCGGTAATACGACTCACTATAGGGGTTCAGAGTTCTACAGTCCGACGATCNNNCGTCNNNTCGAAG |
| ABID_SBC_top_177 | CACGGTTA | GGTGATCCGGTAATACGACTCACTATAGGGGTTCAGAGTTCTACAGTCCGACGATCNNNCACGNNNGTTAAG |
| ABID_SBC_top_178 | CGGACATA | GGTGATCCGGTAATACGACTCACTATAGGGGTTCAGAGTTCTACAGTCCGACGATCNNNCGGANNNCATAAG |
| ABID_SBC_top_179 | CTTGTATC | GGTGATCCGGTAATACGACTCACTATAGGGGTTCAGAGTTCTACAGTCCGACGATCNNNCTTGNNNTATCAG |
| ABID_SBC_top_180 | ACTAGCCT | GGTGATCCGGTAATACGACTCACTATAGGGGTTCAGAGTTCTACAGTCCGACGATCNNNACTANNNGCCTAG |
| ABID_SBC_top_181 | TGTGTGAA | GGTGATCCGGTAATACGACTCACTATAGGGGTTCAGAGTTCTACAGTCCGACGATCNNNTGTGNNNTGAAAG |
| ABID_SBC_top_182 | GATGTAAG | GGTGATCCGGTAATACGACTCACTATAGGGGTTCAGAGTTCTACAGTCCGACGATCNNNGATGNNNTAAGAG |
| ABID_SBC_top_183 | CGTGGCAT | GGTGATCCGGTAATACGACTCACTATAGGGGTTCAGAGTTCTACAGTCCGACGATCNNNCGTGNNNGCATAG |
| ABID_SBC_top_184 | TGATGAAC | GGTGATCCGGTAATACGACTCACTATAGGGGTTCAGAGTTCTACAGTCCGACGATCNNNTGATNNNGAACAG |
| ABID_SBC_top_185 | GCGTATAA | GGTGATCCGGTAATACGACTCACTATAGGGGTTCAGAGTTCTACAGTCCGACGATCNNNGCGTNNNATAAAG |
| ABID_SBC_top_186 | ACGAAGCT | GGTGATCCGGTAATACGACTCACTATAGGGGTTCAGAGTTCTACAGTCCGACGATCNNNACGANNNAGCTAG |
| ABID_SBC_top_187 | GTGCGTAC | GGTGATCCGGTAATACGACTCACTATAGGGGTTCAGAGTTCTACAGTCCGACGATCNNNGTGCNNNGTACAG |
| ABID_SBC_top_188 | TCTAGTGA | GGTGATCCGGTAATACGACTCACTATAGGGGTTCAGAGTTCTACAGTCCGACGATCNNNTCTANNNGTGAAG |
| ABID_SBC_top_189 | TCGAGCAC | GGTGATCCGGTAATACGACTCACTATAGGGGTTCAGAGTTCTACAGTCCGACGATCNNNTCGANNNGCACAG |
| ABID_SBC_top_190 | TGTACGTC | GGTGATCCGGTAATACGACTCACTATAGGGGTTCAGAGTTCTACAGTCCGACGATCNNNTGTANNNCGTCAG |
| ABID_SBC_top_191 | GAGTCTAC | GGTGATCCGGTAATACGACTCACTATAGGGGTTCAGAGTTCTACAGTCCGACGATCNNNGAGTNNNCTACAG |
| ABID_SBC_top_192 | GAGTGTCA | GGTGATCCGGTAATACGACTCACTATAGGGGTTCAGAGTTCTACAGTCCGACGATCNNNGAGTNNNGTCAAG |
| ABID_SBC_top_193 | GTGAGACT | GGTGATCCGGTAATACGACTCACTATAGGGGTTCAGAGTTCTACAGTCCGACGATCNNNGTGANNNGACTAG |
| ABID_SBC_top_194 | GCTGCAAC | GGTGATCCGGTAATACGACTCACTATAGGGGTTCAGAGTTCTACAGTCCGACGATCNNNGCTGNNNCAACAG |
| ABID_SBC_top_195 | TGCAGCAG | GGTGATCCGGTAATACGACTCACTATAGGGGTTCAGAGTTCTACAGTCCGACGATCNNNTGCANNNGCAGAG |
| ABID_SBC_top_196 | GTAGACCA | GGTGATCCGGTAATACGACTCACTATAGGGGTTCAGAGTTCTACAGTCCGACGATCNNNGTAGNNNACCAAG |
| ABID_SBC_top_197 | GTACGCCT | GGTGATCCGGTAATACGACTCACTATAGGGGTTCAGAGTTCTACAGTCCGACGATCNNNGTACNNNGCCTAG |

| | | |
|---|---|---|
| ABID_SBC_top_198 | GAAGGCTC | GGTGATCCGGTAATACGACTCACTATAGGGGTTCAGAGTTCTACAGTCCGACGATCNNNGAAGNNNGCTCAG |
| ABID_SBC_top_199 | TCTCGACC | GGTGATCCGGTAATACGACTCACTATAGGGGTTCAGAGTTCTACAGTCCGACGATCNNNTCTCNNNGACCAG |
| ABID_SBC_top_200 | CTCTACGA | GGTGATCCGGTAATACGACTCACTATAGGGGTTCAGAGTTCTACAGTCCGACGATCNNNCTCTNNNACGAAG |
| ABID_SBC_top_201 | GTCACGAC | GGTGATCCGGTAATACGACTCACTATAGGGGTTCAGAGTTCTACAGTCCGACGATCNNNGTCANNNCGACAG |
| ABID_SBC_top_202 | TGTCGCTT | GGTGATCCGGTAATACGACTCACTATAGGGGTTCAGAGTTCTACAGTCCGACGATCNNNTGTCNNNGCTTAG |
| ABID_SBC_top_203 | CATGCTCA | GGTGATCCGGTAATACGACTCACTATAGGGGTTCAGAGTTCTACAGTCCGACGATCNNNCATGNNNCTCAAG |
| ABID_SBC_top_204 | GACTCGCA | GGTGATCCGGTAATACGACTCACTATAGGGGTTCAGAGTTCTACAGTCCGACGATCNNNGACTNNNCGCAAG |
| ABID_SBC_top_205 | TGTGCACC | GGTGATCCGGTAATACGACTCACTATAGGGGTTCAGAGTTCTACAGTCCGACGATCNNNTGTGNNNCACCAG |
| ABID_SBC_top_206 | GTAGCAGA | GGTGATCCGGTAATACGACTCACTATAGGGGTTCAGAGTTCTACAGTCCGACGATCNNNGTAGNNNCAGAAG |
| ABID_SBC_top_207 | CTGTACAC | GGTGATCCGGTAATACGACTCACTATAGGGGTTCAGAGTTCTACAGTCCGACGATCNNNCTGTNNNACACAG |
| ABID_SBC_top_208 | CTTGACGC | GGTGATCCGGTAATACGACTCACTATAGGGGTTCAGAGTTCTACAGTCCGACGATCNNNCTTGNNNACGCAG |
| ABID_SBC_top_209 | AGCGCGTT | GGTGATCCGGTAATACGACTCACTATAGGGGTTCAGAGTTCTACAGTCCGACGATCNNNAGCGNNNCGTTAG |
| ABID_SBC_top_210 | ACACCGAG | GGTGATCCGGTAATACGACTCACTATAGGGGTTCAGAGTTCTACAGTCCGACGATCNNNACACNNNCGAGAG |
| ABID_SBC_top_211 | TCTGGAGT | GGTGATCCGGTAATACGACTCACTATAGGGGTTCAGAGTTCTACAGTCCGACGATCNNNTCTGNNNGAGTAG |
| ABID_SBC_top_212 | CGCTCAAG | GGTGATCCGGTAATACGACTCACTATAGGGGTTCAGAGTTCTACAGTCCGACGATCNNNCGCTNNNCAAGAG |
| ABID_SBC_top_213 | AGGAGTGT | GGTGATCCGGTAATACGACTCACTATAGGGGTTCAGAGTTCTACAGTCCGACGATCNNNAGGANNNGTGTAG |
| ABID_SBC_top_214 | GCACCTTC | GGTGATCCGGTAATACGACTCACTATAGGGGTTCAGAGTTCTACAGTCCGACGATCNNNGCACNNNCTTCAG |
| ABID_SBC_top_215 | AGATTCGT | GGTGATCCGGTAATACGACTCACTATAGGGGTTCAGAGTTCTACAGTCCGACGATCNNNAGATNNNTCGTAG |
| ABID_SBC_top_216 | TGTGATGT | GGTGATCCGGTAATACGACTCACTATAGGGGTTCAGAGTTCTACAGTCCGACGATCNNNTGTGNNNATGTAG |
| ABID_SBC_top_217 | CTGCGAGA | GGTGATCCGGTAATACGACTCACTATAGGGGTTCAGAGTTCTACAGTCCGACGATCNNNCTGCNNNGAGAAG |
| ABID_SBC_top_218 | TGCGAGCT | GGTGATCCGGTAATACGACTCACTATAGGGGTTCAGAGTTCTACAGTCCGACGATCNNNTGCGNNNAGCTAG |
| ABID_SBC_top_219 | ATAGAGAG | GGTGATCCGGTAATACGACTCACTATAGGGGTTCAGAGTTCTACAGTCCGACGATCNNNATAGNNNAGAGAG |
| ABID_SBC_top_220 | ATGCTAGC | GGTGATCCGGTAATACGACTCACTATAGGGGTTCAGAGTTCTACAGTCCGACGATCNNNATGCNNNTAGCAG |
| ABID_SBC_top_221 | GCGTACTG | GGTGATCCGGTAATACGACTCACTATAGGGGTTCAGAGTTCTACAGTCCGACGATCNNNGCGTNNNACTGAG |
| ABID_SBC_top_222 | CGTGTGGT | GGTGATCCGGTAATACGACTCACTATAGGGGTTCAGAGTTCTACAGTCCGACGATCNNNCGTGNNNTGGTAG |
| ABID_SBC_top_223 | ATCGACTA | GGTGATCCGGTAATACGACTCACTATAGGGGTTCAGAGTTCTACAGTCCGACGATCNNNATCGNNNACTAAG |
| ABID_SBC_top_224 | ACAGGTTA | GGTGATCCGGTAATACGACTCACTATAGGGGTTCAGAGTTCTACAGTCCGACGATCNNNACAGNNNGTTAAG |
| ABID_SBC_top_225 | CGCTTAGC | GGTGATCCGGTAATACGACTCACTATAGGGGTTCAGAGTTCTACAGTCCGACGATCNNNCGCTNNNTAGCAG |
| ABID_SBC_top_226 | CTTCGCCA | GGTGATCCGGTAATACGACTCACTATAGGGGTTCAGAGTTCTACAGTCCGACGATCNNNCTTCNNNGCCAAG |
| ABID_SBC_top_227 | TCGAATCA | GGTGATCCGGTAATACGACTCACTATAGGGGTTCAGAGTTCTACAGTCCGACGATCNNNTCGANNNATCAAG |
| ABID_SBC_top_228 | AGGACGAT | GGTGATCCGGTAATACGACTCACTATAGGGGTTCAGAGTTCTACAGTCCGACGATCNNNAGGANNNCGATAG |
| ABID_SBC_top_229 | AGCGATGC | GGTGATCCGGTAATACGACTCACTATAGGGGTTCAGAGTTCTACAGTCCGACGATCNNNAGCGNNNATGCAG |
| ABID_SBC_top_230 | TCTCTCGC | GGTGATCCGGTAATACGACTCACTATAGGGGTTCAGAGTTCTACAGTCCGACGATCNNNTCTCNNNTCGCAG |

| | | |
|---|---|---|
| ABID_SBC_top_231 | GCTGTCCA | GGTGATCCGGTAATACGACTCACTATAGGGGTTCAGAGTTCTACAGTCCGACGATCNNNGCTGNNNTCCAAG |
| ABID_SBC_top_232 | GAGCTGAC | GGTGATCCGGTAATACGACTCACTATAGGGGTTCAGAGTTCTACAGTCCGACGATCNNNGAGCNNNTGACAG |
| ABID_SBC_top_233 | CTTCATGT | GGTGATCCGGTAATACGACTCACTATAGGGGTTCAGAGTTCTACAGTCCGACGATCNNNCTTCNNNATGTAG |
| ABID_SBC_top_234 | TATACTCG | GGTGATCCGGTAATACGACTCACTATAGGGGTTCAGAGTTCTACAGTCCGACGATCNNNTATANNNCTCGAG |
| ABID_SBC_top_235 | TCCTTATG | GGTGATCCGGTAATACGACTCACTATAGGGGTTCAGAGTTCTACAGTCCGACGATCNNNTCCTNNNTATGAG |
| ABID_SBC_top_236 | AGGTTAAG | GGTGATCCGGTAATACGACTCACTATAGGGGTTCAGAGTTCTACAGTCCGACGATCNNNAGGTNNNTAAGAG |
| ABID_SBC_top_237 | GACAAGTA | GGTGATCCGGTAATACGACTCACTATAGGGGTTCAGAGTTCTACAGTCCGACGATCNNNGACANNNAGTAAG |
| ABID_SBC_top_238 | TGACGTTC | GGTGATCCGGTAATACGACTCACTATAGGGGTTCAGAGTTCTACAGTCCGACGATCNNNTGACNNNGTTCAG |
| ABID_SBC_top_239 | GTGATATC | GGTGATCCGGTAATACGACTCACTATAGGGGTTCAGAGTTCTACAGTCCGACGATCNNNGTGANNNTATCAG |
| ABID_SBC_top_240 | GTTAGCAA | GGTGATCCGGTAATACGACTCACTATAGGGGTTCAGAGTTCTACAGTCCGACGATCNNNGTTANNNGCAAAG |
| ABID_SBC_top_241 | TCATTGTC | GGTGATCCGGTAATACGACTCACTATAGGGGTTCAGAGTTCTACAGTCCGACGATCNNNTCATNNNTGTCAG |
| ABID_SBC_top_242 | TGGTTCAT | GGTGATCCGGTAATACGACTCACTATAGGGGTTCAGAGTTCTACAGTCCGACGATCNNNTGGTNNNTCATAG |
| ABID_SBC_top_243 | CAATCGTT | GGTGATCCGGTAATACGACTCACTATAGGGGTTCAGAGTTCTACAGTCCGACGATCNNNCAATNNNCGTTAG |
| ABID_SBC_top_244 | ATACGATG | GGTGATCCGGTAATACGACTCACTATAGGGGTTCAGAGTTCTACAGTCCGACGATCNNNATACNNNGATGAG |
| ABID_SBC_top_245 | TAGTAGAG | GGTGATCCGGTAATACGACTCACTATAGGGGTTCAGAGTTCTACAGTCCGACGATCNNNTAGTNNNAGAGAG |
| ABID_SBC_top_246 | ATCAGCGA | GGTGATCCGGTAATACGACTCACTATAGGGGTTCAGAGTTCTACAGTCCGACGATCNNNATCANNNGCGAAG |
| ABID_SBC_top_247 | TCCGGAAG | GGTGATCCGGTAATACGACTCACTATAGGGGTTCAGAGTTCTACAGTCCGACGATCNNNTCCGNNNGAAGAG |
| ABID_SBC_top_248 | ACGTGCAA | GGTGATCCGGTAATACGACTCACTATAGGGGTTCAGAGTTCTACAGTCCGACGATCNNNACGTNNNGCAAAG |
| ABID_SBC_top_249 | TACGTCCT | GGTGATCCGGTAATACGACTCACTATAGGGGTTCAGAGTTCTACAGTCCGACGATCNNNTACGNNNTCCTAG |
| ABID_SBC_top_250 | TCCTCGAG | GGTGATCCGGTAATACGACTCACTATAGGGGTTCAGAGTTCTACAGTCCGACGATCNNNTCCTNNNCGAGAG |
| ABID_SBC_top_251 | CTCTGCAG | GGTGATCCGGTAATACGACTCACTATAGGGGTTCAGAGTTCTACAGTCCGACGATCNNNCTCTNNNGCAGAG |
| ABID_SBC_top_252 | ACGTCTGA | GGTGATCCGGTAATACGACTCACTATAGGGGTTCAGAGTTCTACAGTCCGACGATCNNNACGTNNNCTGAAG |
| ABID_SBC_top_253 | CGACAGGA | GGTGATCCGGTAATACGACTCACTATAGGGGTTCAGAGTTCTACAGTCCGACGATCNNNCGACNNNAGGAAG |
| ABID_SBC_top_254 | GTCGAGAA | GGTGATCCGGTAATACGACTCACTATAGGGGTTCAGAGTTCTACAGTCCGACGATCNNNGTCGNNNAGAAAG |
| ABID_SBC_top_255 | ACTGATCT | GGTGATCCGGTAATACGACTCACTATAGGGGTTCAGAGTTCTACAGTCCGACGATCNNNACTGNNNATCTAG |
| ABID_SBC_top_256 | ACCAGCTG | GGTGATCCGGTAATACGACTCACTATAGGGGTTCAGAGTTCTACAGTCCGACGATCNNNACCANNNGCTGAG |
| ABID_SBC_top_257 | CTATGATC | GGTGATCCGGTAATACGACTCACTATAGGGGTTCAGAGTTCTACAGTCCGACGATCNNNCTATNNNGATCAG |
| ABID_SBC_top_258 | ATCTGTGC | GGTGATCCGGTAATACGACTCACTATAGGGGTTCAGAGTTCTACAGTCCGACGATCNNNATCTNNNGTGCAG |
| ABID_SBC_top_259 | GATAGCCG | GGTGATCCGGTAATACGACTCACTATAGGGGTTCAGAGTTCTACAGTCCGACGATCNNNGATANNNGCCGAG |
| ABID_SBC_top_260 | GATCACCT | GGTGATCCGGTAATACGACTCACTATAGGGGTTCAGAGTTCTACAGTCCGACGATCNNNGATCNNNACCTAG |
| ABID_SBC_top_261 | GACGATGT | GGTGATCCGGTAATACGACTCACTATAGGGGTTCAGAGTTCTACAGTCCGACGATCNNNGACGNNNATGTAG |
| ABID_SBC_top_262 | ATCACTCT | GGTGATCCGGTAATACGACTCACTATAGGGGTTCAGAGTTCTACAGTCCGACGATCNNNATCANNNCTCTAG |
| ABID_SBC_top_263 | CAGTGAAT | GGTGATCCGGTAATACGACTCACTATAGGGGTTCAGAGTTCTACAGTCCGACGATCNNNCAGTNNNGAATAG |

| | | |
|---|---|---|
| ABID_SBC_top_264 | GATCGACA | GGTGATCCGGTAATACGACTCACTATAGGGGTTCAGAGTTCTACAGTCCGACGATCNNNGATCNNNGACAAG |
| ABID_SBC_top_265 | CAGACAAG | GGTGATCCGGTAATACGACTCACTATAGGGGTTCAGAGTTCTACAGTCCGACGATCNNNCAGANNNCAAGAG |
| ABID_SBC_top_266 | TCGCCTTA | GGTGATCCGGTAATACGACTCACTATAGGGGTTCAGAGTTCTACAGTCCGACGATCNNNTCGCNNNCTTAAG |
| ABID_SBC_top_267 | GTGCATCG | GGTGATCCGGTAATACGACTCACTATAGGGGTTCAGAGTTCTACAGTCCGACGATCNNNGTGCNNNATCGAG |
| ABID_SBC_top_268 | GTAGGTAA | GGTGATCCGGTAATACGACTCACTATAGGGGTTCAGAGTTCTACAGTCCGACGATCNNNGTAGNNNGTAAAG |
| ABID_SBC_top_269 | TGGCTATC | GGTGATCCGGTAATACGACTCACTATAGGGGTTCAGAGTTCTACAGTCCGACGATCNNNTGGCNNNTATCAG |
| ABID_SBC_top_270 | TCGTCACT | GGTGATCCGGTAATACGACTCACTATAGGGGTTCAGAGTTCTACAGTCCGACGATCNNNTCGTNNNCACTAG |
| ABID_SBC_top_271 | ATCTTCCT | GGTGATCCGGTAATACGACTCACTATAGGGGTTCAGAGTTCTACAGTCCGACGATCNNNATCTNNNTCCTAG |
| ABID_SBC_top_272 | GTGTCGTC | GGTGATCCGGTAATACGACTCACTATAGGGGTTCAGAGTTCTACAGTCCGACGATCNNNGTGTNNNCGTCAG |
| ABID_SBC_top_273 | TGAGTGGC | GGTGATCCGGTAATACGACTCACTATAGGGGTTCAGAGTTCTACAGTCCGACGATCNNNTGAGNNNTGGCAG |
| ABID_SBC_top_274 | CGAGCTGT | GGTGATCCGGTAATACGACTCACTATAGGGGTTCAGAGTTCTACAGTCCGACGATCNNNCGAGNNNCTGTAG |
| ABID_SBC_top_275 | ACGATATG | GGTGATCCGGTAATACGACTCACTATAGGGGTTCAGAGTTCTACAGTCCGACGATCNNNACGANNNTATGAG |
| ABID_SBC_top_276 | CGGTTGCT | GGTGATCCGGTAATACGACTCACTATAGGGGTTCAGAGTTCTACAGTCCGACGATCNNNCGGTNNNTGCTAG |
| ABID_SBC_top_277 | GCCTTGAC | GGTGATCCGGTAATACGACTCACTATAGGGGTTCAGAGTTCTACAGTCCGACGATCNNNGCCTNNNTGACAG |
| ABID_SBC_top_278 | TCTCGCAG | GGTGATCCGGTAATACGACTCACTATAGGGGTTCAGAGTTCTACAGTCCGACGATCNNNTCTCNNNGCAGAG |
| ABID_SBC_top_279 | AGGCTCCT | GGTGATCCGGTAATACGACTCACTATAGGGGTTCAGAGTTCTACAGTCCGACGATCNNNAGGCNNNTCCTAG |
| ABID_SBC_top_280 | TATGCAGA | GGTGATCCGGTAATACGACTCACTATAGGGGTTCAGAGTTCTACAGTCCGACGATCNNNTATGNNNCAGAAG |
| ABID_SBC_top_281 | GAGTCATT | GGTGATCCGGTAATACGACTCACTATAGGGGTTCAGAGTTCTACAGTCCGACGATCNNNGAGTNNNCATTAG |
| ABID_SBC_top_282 | GCGATCTT | GGTGATCCGGTAATACGACTCACTATAGGGGTTCAGAGTTCTACAGTCCGACGATCNNNGCGANNNTCTTAG |
| ABID_SBC_top_283 | AGATCTTC | GGTGATCCGGTAATACGACTCACTATAGGGGTTCAGAGTTCTACAGTCCGACGATCNNNAGATNNNCTTCAG |
| ABID_SBC_top_284 | AGCATCCG | GGTGATCCGGTAATACGACTCACTATAGGGGTTCAGAGTTCTACAGTCCGACGATCNNNAGCANNNTCCGAG |
| ABID_SBC_top_285 | TCTGTCAT | GGTGATCCGGTAATACGACTCACTATAGGGGTTCAGAGTTCTACAGTCCGACGATCNNNTCTGNNNTCATAG |
| ABID_SBC_top_286 | CGGTCAGT | GGTGATCCGGTAATACGACTCACTATAGGGGTTCAGAGTTCTACAGTCCGACGATCNNNCGGTNNNCAGTAG |
| ABID_SBC_top_287 | CTAGTCTA | GGTGATCCGGTAATACGACTCACTATAGGGGTTCAGAGTTCTACAGTCCGACGATCNNNCTAGNNNTCTAAG |
| ABID_SBC_top_288 | GCCGATAC | GGTGATCCGGTAATACGACTCACTATAGGGGTTCAGAGTTCTACAGTCCGACGATCNNNGCCGNNNATACAG |
| ABID_SBC_top_289 | GACTATTC | GGTGATCCGGTAATACGACTCACTATAGGGGTTCAGAGTTCTACAGTCCGACGATCNNNGACTNNNATTCAG |
| ABID_SBC_top_290 | TGTCAGTG | GGTGATCCGGTAATACGACTCACTATAGGGGTTCAGAGTTCTACAGTCCGACGATCNNNTGTCNNNAGTGAG |
| ABID_SBC_top_291 | ATGTGACA | GGTGATCCGGTAATACGACTCACTATAGGGGTTCAGAGTTCTACAGTCCGACGATCNNNATGTNNNGACAAG |
| ABID_SBC_top_292 | AGGTAGTA | GGTGATCCGGTAATACGACTCACTATAGGGGTTCAGAGTTCTACAGTCCGACGATCNNNAGGTNNNAGTAAG |
| ABID_SBC_top_293 | TACGAGGC | GGTGATCCGGTAATACGACTCACTATAGGGGTTCAGAGTTCTACAGTCCGACGATCNNNTACGNNNAGGCAG |
| ABID_SBC_top_294 | CTCAACTG | GGTGATCCGGTAATACGACTCACTATAGGGGTTCAGAGTTCTACAGTCCGACGATCNNNCTCANNNACTGAG |
| ABID_SBC_top_295 | CATATGCT | GGTGATCCGGTAATACGACTCACTATAGGGGTTCAGAGTTCTACAGTCCGACGATCNNNCATANNNTGCTAG |
| ABID_SBC_top_296 | TCATTCCT | GGTGATCCGGTAATACGACTCACTATAGGGGTTCAGAGTTCTACAGTCCGACGATCNNNTCATNNNTCCTAG |

| | | |
|---|---|---|
| ABID_SBC_top_297 | TAAGTACG | GGTGATCCGGTAATACGACTCACTATAGGGGTTCAGAGTTCTACAGTCCGACGATCNNNTAAGNNNTACGAG |
| ABID_SBC_top_298 | CTAGCGAC | GGTGATCCGGTAATACGACTCACTATAGGGGTTCAGAGTTCTACAGTCCGACGATCNNNCTAGNNNCGACAG |
| ABID_SBC_top_299 | CGGTGTAC | GGTGATCCGGTAATACGACTCACTATAGGGGTTCAGAGTTCTACAGTCCGACGATCNNNCGGTNNNGTACAG |
| ABID_SBC_top_300 | TCAGACGT | GGTGATCCGGTAATACGACTCACTATAGGGGTTCAGAGTTCTACAGTCCGACGATCNNNTCAGNNNACGTAG |
| ABID_SBC_top_301 | TACACGTT | GGTGATCCGGTAATACGACTCACTATAGGGGTTCAGAGTTCTACAGTCCGACGATCNNNTACANNNCGTTAG |
| ABID_SBC_top_302 | TATGCGTG | GGTGATCCGGTAATACGACTCACTATAGGGGTTCAGAGTTCTACAGTCCGACGATCNNNTATGNNNCGTGAG |
| ABID_SBC_top_303 | TCCATCGT | GGTGATCCGGTAATACGACTCACTATAGGGGTTCAGAGTTCTACAGTCCGACGATCNNNTCCANNNTCGTAG |
| ABID_SBC_top_304 | TATCAGCA | GGTGATCCGGTAATACGACTCACTATAGGGGTTCAGAGTTCTACAGTCCGACGATCNNNTATCNNNAGCAAG |
| ABID_SBC_top_305 | TGTGGTAC | GGTGATCCGGTAATACGACTCACTATAGGGGTTCAGAGTTCTACAGTCCGACGATCNNNTGTGNNNGTACAG |
| ABID_SBC_top_306 | GAGTTCCT | GGTGATCCGGTAATACGACTCACTATAGGGGTTCAGAGTTCTACAGTCCGACGATCNNNGAGTNNNTCCTAG |
| ABID_SBC_top_307 | GAGAACAG | GGTGATCCGGTAATACGACTCACTATAGGGGTTCAGAGTTCTACAGTCCGACGATCNNNGAGANNNACAGAG |
| ABID_SBC_top_308 | AGTGCTGA | GGTGATCCGGTAATACGACTCACTATAGGGGTTCAGAGTTCTACAGTCCGACGATCNNNAGTGNNNCTGAAG |
| ABID_SBC_top_309 | GACGCGAT | GGTGATCCGGTAATACGACTCACTATAGGGGTTCAGAGTTCTACAGTCCGACGATCNNNGACGNNNCGATAG |
| ABID_SBC_top_310 | GATACATG | GGTGATCCGGTAATACGACTCACTATAGGGGTTCAGAGTTCTACAGTCCGACGATCNNNGATANNNCATGAG |
| ABID_SBC_top_311 | AGGAACAC | GGTGATCCGGTAATACGACTCACTATAGGGGTTCAGAGTTCTACAGTCCGACGATCNNNAGGANNNACACAG |
| ABID_SBC_top_312 | CACTTCAT | GGTGATCCGGTAATACGACTCACTATAGGGGTTCAGAGTTCTACAGTCCGACGATCNNNCACTNNNTCATAG |
| ABID_SBC_top_313 | TAGACGAA | GGTGATCCGGTAATACGACTCACTATAGGGGTTCAGAGTTCTACAGTCCGACGATCNNNTAGANNNCGAAAG |
| ABID_SBC_top_314 | TGATATGC | GGTGATCCGGTAATACGACTCACTATAGGGGTTCAGAGTTCTACAGTCCGACGATCNNNTGATNNNATGCAG |
| ABID_SBC_top_315 | CTATTGCA | GGTGATCCGGTAATACGACTCACTATAGGGGTTCAGAGTTCTACAGTCCGACGATCNNNCTATNNNTGCAAG |
| ABID_SBC_top_316 | GCGTTGTA | GGTGATCCGGTAATACGACTCACTATAGGGGTTCAGAGTTCTACAGTCCGACGATCNNNGCGTNNNTGTAAG |
| ABID_SBC_top_317 | CAGAGACC | GGTGATCCGGTAATACGACTCACTATAGGGGTTCAGAGTTCTACAGTCCGACGATCNNNCAGANNNGACCAG |
| ABID_SBC_top_318 | ACCTACCA | GGTGATCCGGTAATACGACTCACTATAGGGGTTCAGAGTTCTACAGTCCGACGATCNNNACCTNNNACCAAG |
| ABID_SBC_top_319 | CTTCGATT | GGTGATCCGGTAATACGACTCACTATAGGGGTTCAGAGTTCTACAGTCCGACGATCNNNCTTCNNNGATTAG |
| ABID_SBC_top_320 | CGAGATCC | GGTGATCCGGTAATACGACTCACTATAGGGGTTCAGAGTTCTACAGTCCGACGATCNNNCGAGNNNATCCAG |
| ABID_SBC_top_321 | CTGTGCTA | GGTGATCCGGTAATACGACTCACTATAGGGGTTCAGAGTTCTACAGTCCGACGATCNNNCTGTNNNGCTAAG |
| ABID_SBC_top_322 | TAGCGATA | GGTGATCCGGTAATACGACTCACTATAGGGGTTCAGAGTTCTACAGTCCGACGATCNNNTAGCNNNGATAAG |
| ABID_SBC_top_323 | GTTCTAGA | GGTGATCCGGTAATACGACTCACTATAGGGGTTCAGAGTTCTACAGTCCGACGATCNNNGTTCNNNTAGAAG |
| ABID_SBC_top_324 | CTCGGCTT | GGTGATCCGGTAATACGACTCACTATAGGGGTTCAGAGTTCTACAGTCCGACGATCNNNCTCGNNNGCTTAG |
| ABID_SBC_top_325 | TAGTATCC | GGTGATCCGGTAATACGACTCACTATAGGGGTTCAGAGTTCTACAGTCCGACGATCNNNTAGTNNNATCCAG |
| ABID_SBC_top_326 | TCAGCAAT | GGTGATCCGGTAATACGACTCACTATAGGGGTTCAGAGTTCTACAGTCCGACGATCNNNTCAGNNNCAATAG |
| ABID_SBC_top_327 | TCTGCTAA | GGTGATCCGGTAATACGACTCACTATAGGGGTTCAGAGTTCTACAGTCCGACGATCNNNTCTGNNNCTAAAG |
| ABID_SBC_top_328 | CGGATACG | GGTGATCCGGTAATACGACTCACTATAGGGGTTCAGAGTTCTACAGTCCGACGATCNNNCGGANNNTACGAG |
| ABID_SBC_top_329 | CGTGGAGA | GGTGATCCGGTAATACGACTCACTATAGGGGTTCAGAGTTCTACAGTCCGACGATCNNNCGTGNNNGAGAAG |

| | | |
|---|---|---|
| ABID_SBC_top_330 | TCATACAC | GGTGATCCGGTAATACGACTCACTATAGGGGTTCAGAGTTCTACAGTCCGACGATCNNNTCATNNNACACAG |
| ABID_SBC_top_331 | CGGAAGGT | GGTGATCCGGTAATACGACTCACTATAGGGGTTCAGAGTTCTACAGTCCGACGATCNNNCGGANNNAGGTAG |
| ABID_SBC_top_332 | GTTACAGT | GGTGATCCGGTAATACGACTCACTATAGGGGTTCAGAGTTCTACAGTCCGACGATCNNNGTTANNNCAGTAG |
| ABID_SBC_top_333 | GTCTGACC | GGTGATCCGGTAATACGACTCACTATAGGGGTTCAGAGTTCTACAGTCCGACGATCNNNGTCTNNNGACCAG |
| ABID_SBC_top_334 | GTCTTCAA | GGTGATCCGGTAATACGACTCACTATAGGGGTTCAGAGTTCTACAGTCCGACGATCNNNGTCTNNNTCAAAG |
| ABID_SBC_top_335 | GTTCTGCT | GGTGATCCGGTAATACGACTCACTATAGGGGTTCAGAGTTCTACAGTCCGACGATCNNNGTTCNNNTGCTAG |
| ABID_SBC_top_336 | TCCGCACA | GGTGATCCGGTAATACGACTCACTATAGGGGTTCAGAGTTCTACAGTCCGACGATCNNNTCCGNNNCACAAG |
| ABID_SBC_top_337 | ATTCTCGT | GGTGATCCGGTAATACGACTCACTATAGGGGTTCAGAGTTCTACAGTCCGACGATCNNNATTCNNNTCGTAG |
| ABID_SBC_top_338 | TGTGTCCG | GGTGATCCGGTAATACGACTCACTATAGGGGTTCAGAGTTCTACAGTCCGACGATCNNNTGTGNNNTCCGAG |
| ABID_SBC_top_339 | GTCATGTG | GGTGATCCGGTAATACGACTCACTATAGGGGTTCAGAGTTCTACAGTCCGACGATCNNNGTCANNNTGTGAG |
| ABID_SBC_top_340 | GTGCTAAT | GGTGATCCGGTAATACGACTCACTATAGGGGTTCAGAGTTCTACAGTCCGACGATCNNNGTGCNNNTAATAG |
| ABID_SBC_top_341 | GAGATCGC | GGTGATCCGGTAATACGACTCACTATAGGGGTTCAGAGTTCTACAGTCCGACGATCNNNGAGANNNTCGCAG |
| ABID_SBC_top_342 | GCCTGCTA | GGTGATCCGGTAATACGACTCACTATAGGGGTTCAGAGTTCTACAGTCCGACGATCNNNGCCTNNNGCTAAG |
| ABID_SBC_top_343 | GAGAATCT | GGTGATCCGGTAATACGACTCACTATAGGGGTTCAGAGTTCTACAGTCCGACGATCNNNGAGANNNATCTAG |
| ABID_SBC_top_344 | CAGCCTGT | GGTGATCCGGTAATACGACTCACTATAGGGGTTCAGAGTTCTACAGTCCGACGATCNNNCAGCNNNCTGTAG |
| ABID_SBC_top_345 | GTGTATGT | GGTGATCCGGTAATACGACTCACTATAGGGGTTCAGAGTTCTACAGTCCGACGATCNNNGTGTNNNATGTAG |
| ABID_SBC_top_346 | AGACTGTG | GGTGATCCGGTAATACGACTCACTATAGGGGTTCAGAGTTCTACAGTCCGACGATCNNNAGACNNNTGTGAG |
| ABID_SBC_top_347 | GATATGTC | GGTGATCCGGTAATACGACTCACTATAGGGGTTCAGAGTTCTACAGTCCGACGATCNNNGATANNNTGTCAG |
| ABID_SBC_top_348 | TGGACTCC | GGTGATCCGGTAATACGACTCACTATAGGGGTTCAGAGTTCTACAGTCCGACGATCNNNTGGANNNCTCCAG |
| ABID_SBC_top_349 | TGATCGTA | GGTGATCCGGTAATACGACTCACTATAGGGGTTCAGAGTTCTACAGTCCGACGATCNNNTGATNNNCGTAAG |
| ABID_SBC_top_350 | TAGTCGGC | GGTGATCCGGTAATACGACTCACTATAGGGGTTCAGAGTTCTACAGTCCGACGATCNNNTAGTNNNCGGCAG |
| ABID_SBC_top_351 | ACCATGTA | GGTGATCCGGTAATACGACTCACTATAGGGGTTCAGAGTTCTACAGTCCGACGATCNNNACCANNNTGTAAG |
| ABID_SBC_top_352 | TCTACAAG | GGTGATCCGGTAATACGACTCACTATAGGGGTTCAGAGTTCTACAGTCCGACGATCNNNTCTANNNCAAGAG |
| ABID_SBC_top_353 | AGAGTGCT | GGTGATCCGGTAATACGACTCACTATAGGGGTTCAGAGTTCTACAGTCCGACGATCNNNAGAGNNNTGCTAG |
| ABID_SBC_top_354 | CTTGTGCG | GGTGATCCGGTAATACGACTCACTATAGGGGTTCAGAGTTCTACAGTCCGACGATCNNNCTTGNNNTGCGAG |
| ABID_SBC_top_355 | TAATGCGC | GGTGATCCGGTAATACGACTCACTATAGGGGTTCAGAGTTCTACAGTCCGACGATCNNNTAATNNNGCGCAG |
| ABID_SBC_top_356 | TGCTGCGT | GGTGATCCGGTAATACGACTCACTATAGGGGTTCAGAGTTCTACAGTCCGACGATCNNNTGCTNNNGCGTAG |
| ABID_SBC_top_357 | GAGTGAGC | GGTGATCCGGTAATACGACTCACTATAGGGGTTCAGAGTTCTACAGTCCGACGATCNNNGAGTNNNGAGCAG |
| ABID_SBC_top_358 | CGCGATAA | GGTGATCCGGTAATACGACTCACTATAGGGGTTCAGAGTTCTACAGTCCGACGATCNNNCGCGNNNATAAAG |
| ABID_SBC_top_359 | ATGTTCTC | GGTGATCCGGTAATACGACTCACTATAGGGGTTCAGAGTTCTACAGTCCGACGATCNNNATGTNNNTCTCAG |
| ABID_SBC_top_360 | ACTCAGAT | GGTGATCCGGTAATACGACTCACTATAGGGGTTCAGAGTTCTACAGTCCGACGATCNNNACTCNNNAGATAG |
| ABID_SBC_top_361 | GCCATAAG | GGTGATCCGGTAATACGACTCACTATAGGGGTTCAGAGTTCTACAGTCCGACGATCNNNGCCANNNTAAGAG |
| ABID_SBC_top_362 | TGTAACCT | GGTGATCCGGTAATACGACTCACTATAGGGGTTCAGAGTTCTACAGTCCGACGATCNNNTGTANNNACCTAG |

| | | |
|---|---|---|
| ABID_SBC_top_363 | ACAGCTAC | GGTGATCCGGTAATACGACTCACTATAGGGGTTCAGAGTTCTACAGTCCGACGATCNNNACAGNNNCTACAG |
| ABID_SBC_top_364 | GTTGGAGC | GGTGATCCGGTAATACGACTCACTATAGGGGTTCAGAGTTCTACAGTCCGACGATCNNNGTTGNNNGAGCAG |
| ABID_SBC_top_365 | GAATAGGC | GGTGATCCGGTAATACGACTCACTATAGGGGTTCAGAGTTCTACAGTCCGACGATCNNNGAATNNNAGGCAG |
| ABID_SBC_top_366 | TCGTGTAT | GGTGATCCGGTAATACGACTCACTATAGGGGTTCAGAGTTCTACAGTCCGACGATCNNNTCGTNNNGTATAG |
| ABID_SBC_top_367 | ATGCGTTA | GGTGATCCGGTAATACGACTCACTATAGGGGTTCAGAGTTCTACAGTCCGACGATCNNNATGCNNNGTTAAG |
| ABID_SBC_top_368 | ACTGGCGA | GGTGATCCGGTAATACGACTCACTATAGGGGTTCAGAGTTCTACAGTCCGACGATCNNNACTGNNNGCGAAG |
| ABID_SBC_top_369 | GCCTCAAT | GGTGATCCGGTAATACGACTCACTATAGGGGTTCAGAGTTCTACAGTCCGACGATCNNNGCCTNNNCAATAG |
| ABID_SBC_top_370 | GATACGGA | GGTGATCCGGTAATACGACTCACTATAGGGGTTCAGAGTTCTACAGTCCGACGATCNNNGATANNNCGGAAG |
| ABID_SBC_top_371 | CTCAGTAT | GGTGATCCGGTAATACGACTCACTATAGGGGTTCAGAGTTCTACAGTCCGACGATCNNNCTCANNNGTATAG |
| ABID_SBC_top_372 | CAAGATTG | GGTGATCCGGTAATACGACTCACTATAGGGGTTCAGAGTTCTACAGTCCGACGATCNNNCAAGNNNATTGAG |
| ABID_SBC_top_373 | TAGATGTG | GGTGATCCGGTAATACGACTCACTATAGGGGTTCAGAGTTCTACAGTCCGACGATCNNNTAGANNNTGTGAG |
| ABID_SBC_top_374 | CATGAGAT | GGTGATCCGGTAATACGACTCACTATAGGGGTTCAGAGTTCTACAGTCCGACGATCNNNCATGNNNAGATAG |
| ABID_SBC_top_375 | CTTAATCG | GGTGATCCGGTAATACGACTCACTATAGGGGTTCAGAGTTCTACAGTCCGACGATCNNNCTTANNNATCGAG |
| ABID_SBC_top_376 | ATTATGCC | GGTGATCCGGTAATACGACTCACTATAGGGGTTCAGAGTTCTACAGTCCGACGATCNNNATTANNNTGCCAG |
| ABID_SBC_top_377 | AGTGGCTC | GGTGATCCGGTAATACGACTCACTATAGGGGTTCAGAGTTCTACAGTCCGACGATCNNNAGTGNNNGCTCAG |
| ABID_SBC_top_378 | ACACGTGC | GGTGATCCGGTAATACGACTCACTATAGGGGTTCAGAGTTCTACAGTCCGACGATCNNNACACNNNGTGCAG |
| ABID_SBC_top_379 | CTTACGTG | GGTGATCCGGTAATACGACTCACTATAGGGGTTCAGAGTTCTACAGTCCGACGATCNNNCTTANNNCGTGAG |
| ABID_SBC_top_380 | CGTGGTTG | GGTGATCCGGTAATACGACTCACTATAGGGGTTCAGAGTTCTACAGTCCGACGATCNNNCGTGNNNGTTGAG |
| ABID_SBC_top_381 | TGGATAGT | GGTGATCCGGTAATACGACTCACTATAGGGGTTCAGAGTTCTACAGTCCGACGATCNNNTGGANNNTAGTAG |
| ABID_SBC_top_382 | GATCTCAA | GGTGATCCGGTAATACGACTCACTATAGGGGTTCAGAGTTCTACAGTCCGACGATCNNNGATCNNNTCAAAG |
| ABID_SBC_top_383 | GCCGACTT | GGTGATCCGGTAATACGACTCACTATAGGGGTTCAGAGTTCTACAGTCCGACGATCNNNGCCGNNNACTTAG |
| ABID_SBC_top_384 | ATCTCGAT | GGTGATCCGGTAATACGACTCACTATAGGGGTTCAGAGTTCTACAGTCCGACGATCNNNATCTNNNCGATAG |
| **Name** | **SBC barcode** | **Sequence (5' - 3')** |
| ABID_SBC_bot_001 | AATGGCCT | /5Phos/GGCCCTAATGNNNGCCTNNNGATCGTCGGACTGTAGAACTCTGAACCCCTATAGTGAGTCGTATTACCGGGAGCTT |
| ABID_SBC_bot_002 | TAGCCTCA | /5Phos/GGCCCTTAGCNNNCTCANNNGATCGTCGGACTGTAGAACTCTGAACCCCTATAGTGAGTCGTATTACCGGGAGCTT |
| ABID_SBC_bot_003 | TAGCTCTC | /5Phos/GGCCCTTAGCNNNTCTCNNNGATCGTCGGACTGTAGAACTCTGAACCCCTATAGTGAGTCGTATTACCGGGAGCTT |
| ABID_SBC_bot_004 | CTCGGAAC | /5Phos/GGCCCTCTCGNNNGAACNNNGATCGTCGGACTGTAGAACTCTGAACCCCTATAGTGAGTCGTATTACCGGGAGCTT |
| ABID_SBC_bot_005 | GTCACAAC | /5Phos/GGCCCTGTCANNNCAACNNNGATCGTCGGACTGTAGAACTCTGAACCCCTATAGTGAGTCGTATTACCGGGAGCTT |
| ABID_SBC_bot_006 | AGTCTGTA | /5Phos/GGCCCTAGTCNNNTGTANNNGATCGTCGGACTGTAGAACTCTGAACCCCTATAGTGAGTCGTATTACCGGGAGCTT |
| ABID_SBC_bot_007 | AGCTGCTG | /5Phos/GGCCCTAGCTNNNGCTGNNNGATCGTCGGACTGTAGAACTCTGAACCCCTATAGTGAGTCGTATTACCGGGAGCTT |
| ABID_SBC_bot_008 | TGACTCAG | /5Phos/GGCCCTTGACNNNTCAGNNNGATCGTCGGACTGTAGAACTCTGAACCCCTATAGTGAGTCGTATTACCGGGAGCTT |
| ABID_SBC_bot_009 | GCAGCATC | /5Phos/GGCCCTGCAGNNNCATCNNNGATCGTCGGACTGTAGAACTCTGAACCCCTATAGTGAGTCGTATTACCGGGAGCTT |
| ABID_SBC_bot_010 | CTGCGCAT | /5Phos/GGCCCTCTGCNNNGCATNNNGATCGTCGGACTGTAGAACTCTGAACCCCTATAGTGAGTCGTATTACCGGGAGCTT |

| | | |
|---|---|---|
| ABID_SBC_bot_011 | CAAGAGGT | /5Phos/GGCCCTCAAGNNNAGGTNNNGATCGTCGGACTGTAGAACTCTGAACCCCTATAGTGAGTCGTATTACCGGGAGCTT |
| ABID_SBC_bot_012 | ACCGGATA | /5Phos/GGCCCTACCGNNNGATANNNGATCGTCGGACTGTAGAACTCTGAACCCCTATAGTGAGTCGTATTACCGGGAGCTT |
| ABID_SBC_bot_013 | CAGTGTTA | /5Phos/GGCCCTCAGTNNNGTTANNNGATCGTCGGACTGTAGAACTCTGAACCCCTATAGTGAGTCGTATTACCGGGAGCTT |
| ABID_SBC_bot_014 | AACTCAAC | /5Phos/GGCCCTAACTNNNCAACNNNGATCGTCGGACTGTAGAACTCTGAACCCCTATAGTGAGTCGTATTACCGGGAGCTT |
| ABID_SBC_bot_015 | GGACAGGA | /5Phos/GGCCCTGGACNNNAGGANNNGATCGTCGGACTGTAGAACTCTGAACCCCTATAGTGAGTCGTATTACCGGGAGCTT |
| ABID_SBC_bot_016 | CTTCCAAG | /5Phos/GGCCCTCTTCNNNCAAGNNNGATCGTCGGACTGTAGAACTCTGAACCCCTATAGTGAGTCGTATTACCGGGAGCTT |
| ABID_SBC_bot_017 | CAATACAG | /5Phos/GGCCCTCAATNNNACAGNNNGATCGTCGGACTGTAGAACTCTGAACCCCTATAGTGAGTCGTATTACCGGGAGCTT |
| ABID_SBC_bot_018 | ATAGTACG | /5Phos/GGCCCTATAGNNNTACGNNNGATCGTCGGACTGTAGAACTCTGAACCCCTATAGTGAGTCGTATTACCGGGAGCTT |
| ABID_SBC_bot_019 | GGAGATAC | /5Phos/GGCCCTGGAGNNNATACNNNGATCGTCGGACTGTAGAACTCTGAACCCCTATAGTGAGTCGTATTACCGGGAGCTT |
| ABID_SBC_bot_020 | TTAGACCA | /5Phos/GGCCCTTTAGNNNACCANNNGATCGTCGGACTGTAGAACTCTGAACCCCTATAGTGAGTCGTATTACCGGGAGCTT |
| ABID_SBC_bot_021 | GGACGATG | /5Phos/GGCCCTGGACNNNGATGNNNGATCGTCGGACTGTAGAACTCTGAACCCCTATAGTGAGTCGTATTACCGGGAGCTT |
| ABID_SBC_bot_022 | AGATAGAG | /5Phos/GGCCCTAGATNNNAGAGNNNGATCGTCGGACTGTAGAACTCTGAACCCCTATAGTGAGTCGTATTACCGGGAGCTT |
| ABID_SBC_bot_023 | ACAGTAGT | /5Phos/GGCCCTACAGNNNTAGTNNNGATCGTCGGACTGTAGAACTCTGAACCCCTATAGTGAGTCGTATTACCGGGAGCTT |
| ABID_SBC_bot_024 | CTACCGTC | /5Phos/GGCCCTCTACNNNCGTCNNNGATCGTCGGACTGTAGAACTCTGAACCCCTATAGTGAGTCGTATTACCGGGAGCTT |
| ABID_SBC_bot_025 | AGACATCA | /5Phos/GGCCCTAGACNNNATCANNNGATCGTCGGACTGTAGAACTCTGAACCCCTATAGTGAGTCGTATTACCGGGAGCTT |
| ABID_SBC_bot_026 | ACACGTTA | /5Phos/GGCCCTACACNNNGTTANNNGATCGTCGGACTGTAGAACTCTGAACCCCTATAGTGAGTCGTATTACCGGGAGCTT |
| ABID_SBC_bot_027 | CGGAATCG | /5Phos/GGCCCTCGGANNNATCGNNNGATCGTCGGACTGTAGAACTCTGAACCCCTATAGTGAGTCGTATTACCGGGAGCTT |
| ABID_SBC_bot_028 | AGACGAGC | /5Phos/GGCCCTAGACNNNGAGCNNNGATCGTCGGACTGTAGAACTCTGAACCCCTATAGTGAGTCGTATTACCGGGAGCTT |
| ABID_SBC_bot_029 | GAGCACCA | /5Phos/GGCCCTGAGCNNNACCANNNGATCGTCGGACTGTAGAACTCTGAACCCCTATAGTGAGTCGTATTACCGGGAGCTT |
| ABID_SBC_bot_030 | TCGAACTG | /5Phos/GGCCCTTCGANNNACTGNNNGATCGTCGGACTGTAGAACTCTGAACCCCTATAGTGAGTCGTATTACCGGGAGCTT |
| ABID_SBC_bot_031 | TCCGGAGT | /5Phos/GGCCCTTCCGNNNGAGTNNNGATCGTCGGACTGTAGAACTCTGAACCCCTATAGTGAGTCGTATTACCGGGAGCTT |
| ABID_SBC_bot_032 | TGCTTGCG | /5Phos/GGCCCTTGCTNNNTGCGNNNGATCGTCGGACTGTAGAACTCTGAACCCCTATAGTGAGTCGTATTACCGGGAGCTT |
| ABID_SBC_bot_033 | GATCCGAT | /5Phos/GGCCCTGATCNNNCGATNNNGATCGTCGGACTGTAGAACTCTGAACCCCTATAGTGAGTCGTATTACCGGGAGCTT |
| ABID_SBC_bot_034 | ATACCGGT | /5Phos/GGCCCTATACNNNCGGTNNNGATCGTCGGACTGTAGAACTCTGAACCCCTATAGTGAGTCGTATTACCGGGAGCTT |
| ABID_SBC_bot_035 | ATTCCTTC | /5Phos/GGCCCTATTCNNNCTTCNNNGATCGTCGGACTGTAGAACTCTGAACCCCTATAGTGAGTCGTATTACCGGGAGCTT |
| ABID_SBC_bot_036 | CATCCAGT | /5Phos/GGCCCTCATCNNNCAGTNNNGATCGTCGGACTGTAGAACTCTGAACCCCTATAGTGAGTCGTATTACCGGGAGCTT |
| ABID_SBC_bot_037 | TTCATCCG | /5Phos/GGCCCTTTCANNNTCCGNNNGATCGTCGGACTGTAGAACTCTGAACCCCTATAGTGAGTCGTATTACCGGGAGCTT |
| ABID_SBC_bot_038 | ACGTTGAC | /5Phos/GGCCCTACGTNNNTGACNNNGATCGTCGGACTGTAGAACTCTGAACCCCTATAGTGAGTCGTATTACCGGGAGCTT |
| ABID_SBC_bot_039 | GACTAGCA | /5Phos/GGCCCTGACTNNNAGCANNNGATCGTCGGACTGTAGAACTCTGAACCCCTATAGTGAGTCGTATTACCGGGAGCTT |
| ABID_SBC_bot_040 | GTCTGCCA | /5Phos/GGCCCTGTCTNNNGCCANNNGATCGTCGGACTGTAGAACTCTGAACCCCTATAGTGAGTCGTATTACCGGGAGCTT |
| ABID_SBC_bot_041 | TGTCCGAG | /5Phos/GGCCCTTGTCNNNCGAGNNNGATCGTCGGACTGTAGAACTCTGAACCCCTATAGTGAGTCGTATTACCGGGAGCTT |
| ABID_SBC_bot_042 | GAGACGGA | /5Phos/GGCCCTGAGANNNCGGANNNGATCGTCGGACTGTAGAACTCTGAACCCCTATAGTGAGTCGTATTACCGGGAGCTT |
| ABID_SBC_bot_043 | CATCACCG | /5Phos/GGCCCTCATCNNNACCGNNNGATCGTCGGACTGTAGAACTCTGAACCCCTATAGTGAGTCGTATTACCGGGAGCTT |

| | | |
|---|---|---|
| ABID_SBC_bot_044 | TAGTGAGT | /5Phos/GGCCCTTAGTNNNGAGTNNNGATCGTCGGACTGTAGAACTCTGAACCCCTATAGTGAGTCGTATTACCGGGAGCTT |
| ABID_SBC_bot_045 | GGTATCGA | /5Phos/GGCCCTGGTANNNTCGANNNGATCGTCGGACTGTAGAACTCTGAACCCCTATAGTGAGTCGTATTACCGGGAGCTT |
| ABID_SBC_bot_046 | TGTGGTTA | /5Phos/GGCCCTTGTGNNNGTTANNNGATCGTCGGACTGTAGAACTCTGAACCCCTATAGTGAGTCGTATTACCGGGAGCTT |
| ABID_SBC_bot_047 | GAGCTAGC | /5Phos/GGCCCTGAGCNNNTAGCNNNGATCGTCGGACTGTAGAACTCTGAACCCCTATAGTGAGTCGTATTACCGGGAGCTT |
| ABID_SBC_bot_048 | AACGTAGC | /5Phos/GGCCCTAACGNNNTAGCNNNGATCGTCGGACTGTAGAACTCTGAACCCCTATAGTGAGTCGTATTACCGGGAGCTT |
| ABID_SBC_bot_049 | TAACGTTC | /5Phos/GGCCCTTAACNNNGTTCNNNGATCGTCGGACTGTAGAACTCTGAACCCCTATAGTGAGTCGTATTACCGGGAGCTT |
| ABID_SBC_bot_050 | GTAGTGGA | /5Phos/GGCCCTGTAGNNNTGGANNNGATCGTCGGACTGTAGAACTCTGAACCCCTATAGTGAGTCGTATTACCGGGAGCTT |
| ABID_SBC_bot_051 | CAAGACTA | /5Phos/GGCCCTCAAGNNNACTANNNGATCGTCGGACTGTAGAACTCTGAACCCCTATAGTGAGTCGTATTACCGGGAGCTT |
| ABID_SBC_bot_052 | TCATCTAG | /5Phos/GGCCCTTCATNNNCTAGNNNGATCGTCGGACTGTAGAACTCTGAACCCCTATAGTGAGTCGTATTACCGGGAGCTT |
| ABID_SBC_bot_053 | ATGCACAC | /5Phos/GGCCCTATGCNNNACACNNNGATCGTCGGACTGTAGAACTCTGAACCCCTATAGTGAGTCGTATTACCGGGAGCTT |
| ABID_SBC_bot_054 | GAATTCCG | /5Phos/GGCCCTGAATNNNTCCGNNNGATCGTCGGACTGTAGAACTCTGAACCCCTATAGTGAGTCGTATTACCGGGAGCTT |
| ABID_SBC_bot_055 | TCTGTCAT | /5Phos/GGCCCTTCTGNNNTCATNNNGATCGTCGGACTGTAGAACTCTGAACCCCTATAGTGAGTCGTATTACCGGGAGCTT |
| ABID_SBC_bot_056 | GGTAGTCG | /5Phos/GGCCCTGGTANNNGTCGNNNGATCGTCGGACTGTAGAACTCTGAACCCCTATAGTGAGTCGTATTACCGGGAGCTT |
| ABID_SBC_bot_057 | AACTACGA | /5Phos/GGCCCTAACTNNNACGANNNGATCGTCGGACTGTAGAACTCTGAACCCCTATAGTGAGTCGTATTACCGGGAGCTT |
| ABID_SBC_bot_058 | GTTCTATC | /5Phos/GGCCCTGTTCNNNTATCNNNGATCGTCGGACTGTAGAACTCTGAACCCCTATAGTGAGTCGTATTACCGGGAGCTT |
| ABID_SBC_bot_059 | GGCAGTTA | /5Phos/GGCCCTGGCANNNGTTANNNGATCGTCGGACTGTAGAACTCTGAACCCCTATAGTGAGTCGTATTACCGGGAGCTT |
| ABID_SBC_bot_060 | CGTATAAC | /5Phos/GGCCCTCGTANNNTAACNNNGATCGTCGGACTGTAGAACTCTGAACCCCTATAGTGAGTCGTATTACCGGGAGCTT |
| ABID_SBC_bot_061 | GTGACTTG | /5Phos/GGCCCTGTGANNNCTTGNNNGATCGTCGGACTGTAGAACTCTGAACCCCTATAGTGAGTCGTATTACCGGGAGCTT |
| ABID_SBC_bot_062 | CTGATAAG | /5Phos/GGCCCTCTGANNNTAAGNNNGATCGTCGGACTGTAGAACTCTGAACCCCTATAGTGAGTCGTATTACCGGGAGCTT |
| ABID_SBC_bot_063 | GCATTAAC | /5Phos/GGCCCTGCATNNNTAACNNNGATCGTCGGACTGTAGAACTCTGAACCCCTATAGTGAGTCGTATTACCGGGAGCTT |
| ABID_SBC_bot_064 | CATGCGTG | /5Phos/GGCCCTCATGNNNCGTGNNNGATCGTCGGACTGTAGAACTCTGAACCCCTATAGTGAGTCGTATTACCGGGAGCTT |
| ABID_SBC_bot_065 | CGACCAGA | /5Phos/GGCCCTCGACNNNCAGANNNGATCGTCGGACTGTAGAACTCTGAACCCCTATAGTGAGTCGTATTACCGGGAGCTT |
| ABID_SBC_bot_066 | CTATAGCA | /5Phos/GGCCCTCTATNNNAGCANNNGATCGTCGGACTGTAGAACTCTGAACCCCTATAGTGAGTCGTATTACCGGGAGCTT |
| ABID_SBC_bot_067 | ACAGCGGA | /5Phos/GGCCCTACAGNNNCGGANNNGATCGTCGGACTGTAGAACTCTGAACCCCTATAGTGAGTCGTATTACCGGGAGCTT |
| ABID_SBC_bot_068 | GTTGTAAG | /5Phos/GGCCCTGTTGNNNTAAGNNNGATCGTCGGACTGTAGAACTCTGAACCCCTATAGTGAGTCGTATTACCGGGAGCTT |
| ABID_SBC_bot_069 | AGCGCAGA | /5Phos/GGCCCTAGCGNNNCAGANNNGATCGTCGGACTGTAGAACTCTGAACCCCTATAGTGAGTCGTATTACCGGGAGCTT |
| ABID_SBC_bot_070 | TCACGCCA | /5Phos/GGCCCTTCACNNNGCCANNNGATCGTCGGACTGTAGAACTCTGAACCCCTATAGTGAGTCGTATTACCGGGAGCTT |
| ABID_SBC_bot_071 | GCCGATAT | /5Phos/GGCCCTGCCGNNNATATNNNGATCGTCGGACTGTAGAACTCTGAACCCCTATAGTGAGTCGTATTACCGGGAGCTT |
| ABID_SBC_bot_072 | GAATCTAC | /5Phos/GGCCCTGAATNNNCTACNNNGATCGTCGGACTGTAGAACTCTGAACCCCTATAGTGAGTCGTATTACCGGGAGCTT |
| ABID_SBC_bot_073 | CTGCCTCT | /5Phos/GGCCCTCTGCNNNCTCTNNNGATCGTCGGACTGTAGAACTCTGAACCCCTATAGTGAGTCGTATTACCGGGAGCTT |
| ABID_SBC_bot_074 | CAACGATA | /5Phos/GGCCCTCAACNNNGATANNNGATCGTCGGACTGTAGAACTCTGAACCCCTATAGTGAGTCGTATTACCGGGAGCTT |
| ABID_SBC_bot_075 | TTCGATAC | /5Phos/GGCCCTTTCGNNNATACNNNGATCGTCGGACTGTAGAACTCTGAACCCCTATAGTGAGTCGTATTACCGGGAGCTT |
| ABID_SBC_bot_076 | GATCTGCA | /5Phos/GGCCCTGATCNNNTGCANNNGATCGTCGGACTGTAGAACTCTGAACCCCTATAGTGAGTCGTATTACCGGGAGCTT |

| | | |
|---|---|---|
| ABID_SBC_bot_077 | TCGTTCCA | /5Phos/GGCCCTTCGTNNNTCCANNNGATCGTCGGACTGTAGAACTCTGAACCCCTATAGTGAGTCGTATTACCGGGAGCTT |
| ABID_SBC_bot_078 | GGTGAGTA | /5Phos/GGCCCTGGTGNNNAGTANNNGATCGTCGGACTGTAGAACTCTGAACCCCTATAGTGAGTCGTATTACCGGGAGCTT |
| ABID_SBC_bot_079 | GACTCTCT | /5Phos/GGCCCTGACTNNNCTCTNNNGATCGTCGGACTGTAGAACTCTGAACCCCTATAGTGAGTCGTATTACCGGGAGCTT |
| ABID_SBC_bot_080 | GCTCTGGT | /5Phos/GGCCCTGCTCNNNTGGTNNNGATCGTCGGACTGTAGAACTCTGAACCCCTATAGTGAGTCGTATTACCGGGAGCTT |
| ABID_SBC_bot_081 | ACCGATGC | /5Phos/GGCCCTACCGNNNATGCNNNGATCGTCGGACTGTAGAACTCTGAACCCCTATAGTGAGTCGTATTACCGGGAGCTT |
| ABID_SBC_bot_082 | CAGAGCGA | /5Phos/GGCCCTCAGANNNGCGANNNGATCGTCGGACTGTAGAACTCTGAACCCCTATAGTGAGTCGTATTACCGGGAGCTT |
| ABID_SBC_bot_083 | ATGACGCT | /5Phos/GGCCCTATGANNNCGCTNNNGATCGTCGGACTGTAGAACTCTGAACCCCTATAGTGAGTCGTATTACCGGGAGCTT |
| ABID_SBC_bot_084 | TAATGACG | /5Phos/GGCCCTTAATNNNGACGNNNGATCGTCGGACTGTAGAACTCTGAACCCCTATAGTGAGTCGTATTACCGGGAGCTT |
| ABID_SBC_bot_085 | CTCAATCA | /5Phos/GGCCCTCTCANNNATCANNNGATCGTCGGACTGTAGAACTCTGAACCCCTATAGTGAGTCGTATTACCGGGAGCTT |
| ABID_SBC_bot_086 | TAGAAGCG | /5Phos/GGCCCTTAGANNNAGCGNNNGATCGTCGGACTGTAGAACTCTGAACCCCTATAGTGAGTCGTATTACCGGGAGCTT |
| ABID_SBC_bot_087 | GCGCTACA | /5Phos/GGCCCTGCGCNNNTACANNNGATCGTCGGACTGTAGAACTCTGAACCCCTATAGTGAGTCGTATTACCGGGAGCTT |
| ABID_SBC_bot_088 | CGTGACAC | /5Phos/GGCCCTCGTGNNNACACNNNGATCGTCGGACTGTAGAACTCTGAACCCCTATAGTGAGTCGTATTACCGGGAGCTT |
| ABID_SBC_bot_089 | TGACCGCA | /5Phos/GGCCCTTGACNNNCGCANNNGATCGTCGGACTGTAGAACTCTGAACCCCTATAGTGAGTCGTATTACCGGGAGCTT |
| ABID_SBC_bot_090 | GCTGATTG | /5Phos/GGCCCTGCTGNNNATTGNNNGATCGTCGGACTGTAGAACTCTGAACCCCTATAGTGAGTCGTATTACCGGGAGCTT |
| ABID_SBC_bot_091 | CACGCTAC | /5Phos/GGCCCTCACGNNNCTACNNNGATCGTCGGACTGTAGAACTCTGAACCCCTATAGTGAGTCGTATTACCGGGAGCTT |
| ABID_SBC_bot_092 | CACTATCG | /5Phos/GGCCCTCACTNNNATCGNNNGATCGTCGGACTGTAGAACTCTGAACCCCTATAGTGAGTCGTATTACCGGGAGCTT |
| ABID_SBC_bot_093 | GTGAGTGT | /5Phos/GGCCCTGTGANNNGTGTNNNGATCGTCGGACTGTAGAACTCTGAACCCCTATAGTGAGTCGTATTACCGGGAGCTT |
| ABID_SBC_bot_094 | GCGAAGGT | /5Phos/GGCCCTGCGANNNAGGTNNNGATCGTCGGACTGTAGAACTCTGAACCCCTATAGTGAGTCGTATTACCGGGAGCTT |
| ABID_SBC_bot_095 | TCTACGAT | /5Phos/GGCCCTTCTANNNCGATNNNGATCGTCGGACTGTAGAACTCTGAACCCCTATAGTGAGTCGTATTACCGGGAGCTT |
| ABID_SBC_bot_096 | CGACTGTG | /5Phos/GGCCCTCGACNNNTGTGNNNGATCGTCGGACTGTAGAACTCTGAACCCCTATAGTGAGTCGTATTACCGGGAGCTT |
| ABID_SBC_bot_097 | TGGCTGGA | /5Phos/GGCCCTTGGCNNNTGGANNNGATCGTCGGACTGTAGAACTCTGAACCCCTATAGTGAGTCGTATTACCGGGAGCTT |
| ABID_SBC_bot_098 | TGTAAGCA | /5Phos/GGCCCTTGTANNNAGCANNNGATCGTCGGACTGTAGAACTCTGAACCCCTATAGTGAGTCGTATTACCGGGAGCTT |
| ABID_SBC_bot_099 | TGCAGCCA | /5Phos/GGCCCTTGCANNNGCCANNNGATCGTCGGACTGTAGAACTCTGAACCCCTATAGTGAGTCGTATTACCGGGAGCTT |
| ABID_SBC_bot_100 | TCGACGCA | /5Phos/GGCCCTTCGANNNCGCANNNGATCGTCGGACTGTAGAACTCTGAACCCCTATAGTGAGTCGTATTACCGGGAGCTT |
| ABID_SBC_bot_101 | GTACTGCT | /5Phos/GGCCCTGTACNNNTGCTNNNGATCGTCGGACTGTAGAACTCTGAACCCCTATAGTGAGTCGTATTACCGGGAGCTT |
| ABID_SBC_bot_102 | TCCTAGGC | /5Phos/GGCCCTTCCTNNNAGGCNNNGATCGTCGGACTGTAGAACTCTGAACCCCTATAGTGAGTCGTATTACCGGGAGCTT |
| ABID_SBC_bot_103 | GGACCTTA | /5Phos/GGCCCTGGACNNNCTTANNNGATCGTCGGACTGTAGAACTCTGAACCCCTATAGTGAGTCGTATTACCGGGAGCTT |
| ABID_SBC_bot_104 | TGTCTACA | /5Phos/GGCCCTTGTCNNNTACANNNGATCGTCGGACTGTAGAACTCTGAACCCCTATAGTGAGTCGTATTACCGGGAGCTT |
| ABID_SBC_bot_105 | TCACTGCG | /5Phos/GGCCCTTCACNNNTGCGNNNGATCGTCGGACTGTAGAACTCTGAACCCCTATAGTGAGTCGTATTACCGGGAGCTT |
| ABID_SBC_bot_106 | GGCTTAGA | /5Phos/GGCCCTGGCTNNNTAGANNNGATCGTCGGACTGTAGAACTCTGAACCCCTATAGTGAGTCGTATTACCGGGAGCTT |
| ABID_SBC_bot_107 | CGTCGTCA | /5Phos/GGCCCTCGTCNNNGTCANNNGATCGTCGGACTGTAGAACTCTGAACCCCTATAGTGAGTCGTATTACCGGGAGCTT |
| ABID_SBC_bot_108 | ACTACTAG | /5Phos/GGCCCTACTANNNCTAGNNNGATCGTCGGACTGTAGAACTCTGAACCCCTATAGTGAGTCGTATTACCGGGAGCTT |
| ABID_SBC_bot_109 | GCCTTATG | /5Phos/GGCCCTGCCTNNNTATGNNNGATCGTCGGACTGTAGAACTCTGAACCCCTATAGTGAGTCGTATTACCGGGAGCTT |

| | | |
|---|---|---|
| ABID_SBC_bot_110 | GTATGTCT | /5Phos/GGCCCTGTATNNNGTCTNNNGATCGTCGGACTGTAGAACTCTGAACCCCTATAGTGAGTCGTATTACCGGGAGCTT |
| ABID_SBC_bot_111 | ATTCGTCG | /5Phos/GGCCCTATTCNNNGTCGNNNGATCGTCGGACTGTAGAACTCTGAACCCCTATAGTGAGTCGTATTACCGGGAGCTT |
| ABID_SBC_bot_112 | TGTCCTGC | /5Phos/GGCCCTTGTCNNNCTGCNNNGATCGTCGGACTGTAGAACTCTGAACCCCTATAGTGAGTCGTATTACCGGGAGCTT |
| ABID_SBC_bot_113 | CGGTCATG | /5Phos/GGCCCTCGGTNNNCATGNNNGATCGTCGGACTGTAGAACTCTGAACCCCTATAGTGAGTCGTATTACCGGGAGCTT |
| ABID_SBC_bot_114 | CGACGTAG | /5Phos/GGCCCTCGACNNNGTAGNNNGATCGTCGGACTGTAGAACTCTGAACCCCTATAGTGAGTCGTATTACCGGGAGCTT |
| ABID_SBC_bot_115 | AATCTCGA | /5Phos/GGCCCTAATCNNNTCGANNNGATCGTCGGACTGTAGAACTCTGAACCCCTATAGTGAGTCGTATTACCGGGAGCTT |
| ABID_SBC_bot_116 | CAGAATAC | /5Phos/GGCCCTCAGANNNATACNNNGATCGTCGGACTGTAGAACTCTGAACCCCTATAGTGAGTCGTATTACCGGGAGCTT |
| ABID_SBC_bot_117 | GACAGACG | /5Phos/GGCCCTGACANNNGACGNNNGATCGTCGGACTGTAGAACTCTGAACCCCTATAGTGAGTCGTATTACCGGGAGCTT |
| ABID_SBC_bot_118 | CTGTGACG | /5Phos/GGCCCTCTGTNNNGACGNNNGATCGTCGGACTGTAGAACTCTGAACCCCTATAGTGAGTCGTATTACCGGGAGCTT |
| ABID_SBC_bot_119 | AGGTGCGA | /5Phos/GGCCCTAGGTNNNGCGANNNGATCGTCGGACTGTAGAACTCTGAACCCCTATAGTGAGTCGTATTACCGGGAGCTT |
| ABID_SBC_bot_120 | CAAGGTCG | /5Phos/GGCCCTCAAGNNNGTCGNNNGATCGTCGGACTGTAGAACTCTGAACCCCTATAGTGAGTCGTATTACCGGGAGCTT |
| ABID_SBC_bot_121 | TCCAACAT | /5Phos/GGCCCTTCCANNNACATNNNGATCGTCGGACTGTAGAACTCTGAACCCCTATAGTGAGTCGTATTACCGGGAGCTT |
| ABID_SBC_bot_122 | AGACCGAC | /5Phos/GGCCCTAGACNNNCGACNNNGATCGTCGGACTGTAGAACTCTGAACCCCTATAGTGAGTCGTATTACCGGGAGCTT |
| ABID_SBC_bot_123 | TAACATCG | /5Phos/GGCCCTTAACNNNATCGNNNGATCGTCGGACTGTAGAACTCTGAACCCCTATAGTGAGTCGTATTACCGGGAGCTT |
| ABID_SBC_bot_124 | AATGAGCA | /5Phos/GGCCCTAATGNNNAGCANNNGATCGTCGGACTGTAGAACTCTGAACCCCTATAGTGAGTCGTATTACCGGGAGCTT |
| ABID_SBC_bot_125 | CGATAGGC | /5Phos/GGCCCTCGATNNNAGGCNNNGATCGTCGGACTGTAGAACTCTGAACCCCTATAGTGAGTCGTATTACCGGGAGCTT |
| ABID_SBC_bot_126 | CTTAGAGT | /5Phos/GGCCCTCTTANNNGAGTNNNGATCGTCGGACTGTAGAACTCTGAACCCCTATAGTGAGTCGTATTACCGGGAGCTT |
| ABID_SBC_bot_127 | ACCACTTC | /5Phos/GGCCCTACCANNNCTTCNNNGATCGTCGGACTGTAGAACTCTGAACCCCTATAGTGAGTCGTATTACCGGGAGCTT |
| ABID_SBC_bot_128 | TTCTGTTC | /5Phos/GGCCCTTTCTNNNGTTCNNNGATCGTCGGACTGTAGAACTCTGAACCCCTATAGTGAGTCGTATTACCGGGAGCTT |
| ABID_SBC_bot_129 | TAGATCCT | /5Phos/GGCCCTTAGANNNTCCTNNNGATCGTCGGACTGTAGAACTCTGAACCCCTATAGTGAGTCGTATTACCGGGAGCTT |
| ABID_SBC_bot_130 | GATGGATA | /5Phos/GGCCCTGATGNNNGATANNNGATCGTCGGACTGTAGAACTCTGAACCCCTATAGTGAGTCGTATTACCGGGAGCTT |
| ABID_SBC_bot_131 | TAAGGAAC | /5Phos/GGCCCTTAAGNNNGAACNNNGATCGTCGGACTGTAGAACTCTGAACCCCTATAGTGAGTCGTATTACCGGGAGCTT |
| ABID_SBC_bot_132 | GACTGCGT | /5Phos/GGCCCTGACTNNNGCGTNNNGATCGTCGGACTGTAGAACTCTGAACCCCTATAGTGAGTCGTATTACCGGGAGCTT |
| ABID_SBC_bot_133 | ACACCAAT | /5Phos/GGCCCTACACNNNCAATNNNGATCGTCGGACTGTAGAACTCTGAACCCCTATAGTGAGTCGTATTACCGGGAGCTT |
| ABID_SBC_bot_134 | CGCGTAGT | /5Phos/GGCCCTCGCGNNNTAGTNNNGATCGTCGGACTGTAGAACTCTGAACCCCTATAGTGAGTCGTATTACCGGGAGCTT |
| ABID_SBC_bot_135 | ATGCTGGC | /5Phos/GGCCCTATGCNNNTGGCNNNGATCGTCGGACTGTAGAACTCTGAACCCCTATAGTGAGTCGTATTACCGGGAGCTT |
| ABID_SBC_bot_136 | CTAGCGAG | /5Phos/GGCCCTCTAGNNNCGAGNNNGATCGTCGGACTGTAGAACTCTGAACCCCTATAGTGAGTCGTATTACCGGGAGCTT |
| ABID_SBC_bot_137 | AGTCCATG | /5Phos/GGCCCTAGTCNNNCATGNNNGATCGTCGGACTGTAGAACTCTGAACCCCTATAGTGAGTCGTATTACCGGGAGCTT |
| ABID_SBC_bot_138 | GACTTGAT | /5Phos/GGCCCTGACTNNNTGATNNNGATCGTCGGACTGTAGAACTCTGAACCCCTATAGTGAGTCGTATTACCGGGAGCTT |
| ABID_SBC_bot_139 | AGCGGACT | /5Phos/GGCCCTAGCGNNNGACTNNNGATCGTCGGACTGTAGAACTCTGAACCCCTATAGTGAGTCGTATTACCGGGAGCTT |
| ABID_SBC_bot_140 | AAGTGTAG | /5Phos/GGCCCTAAGTNNNGTAGNNNGATCGTCGGACTGTAGAACTCTGAACCCCTATAGTGAGTCGTATTACCGGGAGCTT |
| ABID_SBC_bot_141 | TACGAGAG | /5Phos/GGCCCTTACGNNNAGAGNNNGATCGTCGGACTGTAGAACTCTGAACCCCTATAGTGAGTCGTATTACCGGGAGCTT |
| ABID_SBC_bot_142 | AACAGCAG | /5Phos/GGCCCTAACANNNGCAGNNNGATCGTCGGACTGTAGAACTCTGAACCCCTATAGTGAGTCGTATTACCGGGAGCTT |

| | | |
|---|---|---|
| ABID_SBC_bot_143 | GCTATGTA | /5Phos/GGCCCTGCTANNNTGTANNNGATCGTCGGACTGTAGAACTCTGAACCCCTATAGTGAGTCGTATTACCGGGAGCTT |
| ABID_SBC_bot_144 | CTACATTG | /5Phos/GGCCCTCTACNNNATTGNNNGATCGTCGGACTGTAGAACTCTGAACCCCTATAGTGAGTCGTATTACCGGGAGCTT |
| ABID_SBC_bot_145 | GAACTAAG | /5Phos/GGCCCTGAACNNNTAAGNNNGATCGTCGGACTGTAGAACTCTGAACCCCTATAGTGAGTCGTATTACCGGGAGCTT |
| ABID_SBC_bot_146 | TGGCAGTG | /5Phos/GGCCCTTGGCNNNAGTGNNNGATCGTCGGACTGTAGAACTCTGAACCCCTATAGTGAGTCGTATTACCGGGAGCTT |
| ABID_SBC_bot_147 | CACGGCAT | /5Phos/GGCCCTCACGNNNGCATNNNGATCGTCGGACTGTAGAACTCTGAACCCCTATAGTGAGTCGTATTACCGGGAGCTT |
| ABID_SBC_bot_148 | GTTACACT | /5Phos/GGCCCTGTTANNNCACTNNNGATCGTCGGACTGTAGAACTCTGAACCCCTATAGTGAGTCGTATTACCGGGAGCTT |
| ABID_SBC_bot_149 | TGCTCAAT | /5Phos/GGCCCTTGCTNNNCAATNNNGATCGTCGGACTGTAGAACTCTGAACCCCTATAGTGAGTCGTATTACCGGGAGCTT |
| ABID_SBC_bot_150 | CGTCACTA | /5Phos/GGCCCTCGTCNNNACTANNNGATCGTCGGACTGTAGAACTCTGAACCCCTATAGTGAGTCGTATTACCGGGAGCTT |
| ABID_SBC_bot_151 | CATGCTCA | /5Phos/GGCCCTCATGNNNCTCANNNGATCGTCGGACTGTAGAACTCTGAACCCCTATAGTGAGTCGTATTACCGGGAGCTT |
| ABID_SBC_bot_152 | ATCTAGCG | /5Phos/GGCCCTATCTNNNAGCGNNNGATCGTCGGACTGTAGAACTCTGAACCCCTATAGTGAGTCGTATTACCGGGAGCTT |
| ABID_SBC_bot_153 | TAATCGGA | /5Phos/GGCCCTTAATNNNCGGANNNGATCGTCGGACTGTAGAACTCTGAACCCCTATAGTGAGTCGTATTACCGGGAGCTT |
| ABID_SBC_bot_154 | CTTGGCCA | /5Phos/GGCCCTCTTGNNNGCCANNNGATCGTCGGACTGTAGAACTCTGAACCCCTATAGTGAGTCGTATTACCGGGAGCTT |
| ABID_SBC_bot_155 | ACCGTCGA | /5Phos/GGCCCTACCGNNNTCGANNNGATCGTCGGACTGTAGAACTCTGAACCCCTATAGTGAGTCGTATTACCGGGAGCTT |
| ABID_SBC_bot_156 | AGTACGGT | /5Phos/GGCCCTAGTANNNCGGTNNNGATCGTCGGACTGTAGAACTCTGAACCCCTATAGTGAGTCGTATTACCGGGAGCTT |
| ABID_SBC_bot_157 | GTTAGATG | /5Phos/GGCCCTGTTANNNGATGNNNGATCGTCGGACTGTAGAACTCTGAACCCCTATAGTGAGTCGTATTACCGGGAGCTT |
| ABID_SBC_bot_158 | GGATTCAT | /5Phos/GGCCCTGGATNNNTCATNNNGATCGTCGGACTGTAGAACTCTGAACCCCTATAGTGAGTCGTATTACCGGGAGCTT |
| ABID_SBC_bot_159 | TCTCACGA | /5Phos/GGCCCTTCTCNNNACGANNNGATCGTCGGACTGTAGAACTCTGAACCCCTATAGTGAGTCGTATTACCGGGAGCTT |
| ABID_SBC_bot_160 | ATCTTCAC | /5Phos/GGCCCTATCTNNNTCACNNNGATCGTCGGACTGTAGAACTCTGAACCCCTATAGTGAGTCGTATTACCGGGAGCTT |
| ABID_SBC_bot_161 | CTTCGCTC | /5Phos/GGCCCTCTTCNNNGCTCNNNGATCGTCGGACTGTAGAACTCTGAACCCCTATAGTGAGTCGTATTACCGGGAGCTT |
| ABID_SBC_bot_162 | AGAGACCT | /5Phos/GGCCCTAGAGNNNACCTNNNGATCGTCGGACTGTAGAACTCTGAACCCCTATAGTGAGTCGTATTACCGGGAGCTT |
| ABID_SBC_bot_163 | AACTGTGC | /5Phos/GGCCCTAACTNNNGTGCNNNGATCGTCGGACTGTAGAACTCTGAACCCCTATAGTGAGTCGTATTACCGGGAGCTT |
| ABID_SBC_bot_164 | ATAGCAAC | /5Phos/GGCCCTATAGNNNCAACNNNGATCGTCGGACTGTAGAACTCTGAACCCCTATAGTGAGTCGTATTACCGGGAGCTT |
| ABID_SBC_bot_165 | CACACGCA | /5Phos/GGCCCTCACANNNCGCANNNGATCGTCGGACTGTAGAACTCTGAACCCCTATAGTGAGTCGTATTACCGGGAGCTT |
| ABID_SBC_bot_166 | GACAGTAC | /5Phos/GGCCCTGACANNNGTACNNNGATCGTCGGACTGTAGAACTCTGAACCCCTATAGTGAGTCGTATTACCGGGAGCTT |
| ABID_SBC_bot_167 | TCCTGCAC | /5Phos/GGCCCTTCCTNNNGCACNNNGATCGTCGGACTGTAGAACTCTGAACCCCTATAGTGAGTCGTATTACCGGGAGCTT |
| ABID_SBC_bot_168 | CGCTATAC | /5Phos/GGCCCTCGCTNNNATACNNNGATCGTCGGACTGTAGAACTCTGAACCCCTATAGTGAGTCGTATTACCGGGAGCTT |
| ABID_SBC_bot_169 | GAAGCAAT | /5Phos/GGCCCTGAAGNNNCAATNNNGATCGTCGGACTGTAGAACTCTGAACCCCTATAGTGAGTCGTATTACCGGGAGCTT |
| ABID_SBC_bot_170 | TCATGAGC | /5Phos/GGCCCTTCATNNNGAGCNNNGATCGTCGGACTGTAGAACTCTGAACCCCTATAGTGAGTCGTATTACCGGGAGCTT |
| ABID_SBC_bot_171 | AATACGAC | /5Phos/GGCCCTAATANNNCGACNNNGATCGTCGGACTGTAGAACTCTGAACCCCTATAGTGAGTCGTATTACCGGGAGCTT |
| ABID_SBC_bot_172 | CTATGCGT | /5Phos/GGCCCTCTATNNNGCGTNNNGATCGTCGGACTGTAGAACTCTGAACCCCTATAGTGAGTCGTATTACCGGGAGCTT |
| ABID_SBC_bot_173 | TCTAATGC | /5Phos/GGCCCTTCTANNNATGCNNNGATCGTCGGACTGTAGAACTCTGAACCCCTATAGTGAGTCGTATTACCGGGAGCTT |
| ABID_SBC_bot_174 | TACAGAGA | /5Phos/GGCCCTTACANNNGAGANNNGATCGTCGGACTGTAGAACTCTGAACCCCTATAGTGAGTCGTATTACCGGGAGCTT |
| ABID_SBC_bot_175 | AGTAGACA | /5Phos/GGCCCTAGTANNNGACANNNGATCGTCGGACTGTAGAACTCTGAACCCCTATAGTGAGTCGTATTACCGGGAGCTT |

| | | |
|---|---|---|
| ABID_SBC_bot_176 | TCGAGACG | /5Phos/GGCCCTTCGANNNGACGNNNGATCGTCGGACTGTAGAACTCTGAACCCCTATAGTGAGTCGTATTACCGGGAGCTT |
| ABID_SBC_bot_177 | TAACCGTG | /5Phos/GGCCCTTAACNNNCGTGNNNGATCGTCGGACTGTAGAACTCTGAACCCCTATAGTGAGTCGTATTACCGGGAGCTT |
| ABID_SBC_bot_178 | TATGTCCG | /5Phos/GGCCCTTATGNNNTCCGNNNGATCGTCGGACTGTAGAACTCTGAACCCCTATAGTGAGTCGTATTACCGGGAGCTT |
| ABID_SBC_bot_179 | GATACAAG | /5Phos/GGCCCTGATANNNCAAGNNNGATCGTCGGACTGTAGAACTCTGAACCCCTATAGTGAGTCGTATTACCGGGAGCTT |
| ABID_SBC_bot_180 | AGGCTAGT | /5Phos/GGCCCTAGGCNNNTAGTNNNGATCGTCGGACTGTAGAACTCTGAACCCCTATAGTGAGTCGTATTACCGGGAGCTT |
| ABID_SBC_bot_181 | TTCACACA | /5Phos/GGCCCTTTCANNNCACANNNGATCGTCGGACTGTAGAACTCTGAACCCCTATAGTGAGTCGTATTACCGGGAGCTT |
| ABID_SBC_bot_182 | CTTACATC | /5Phos/GGCCCTCTTANNNCATCNNNGATCGTCGGACTGTAGAACTCTGAACCCCTATAGTGAGTCGTATTACCGGGAGCTT |
| ABID_SBC_bot_183 | ATGCCACG | /5Phos/GGCCCTATGCNNNCACGNNNGATCGTCGGACTGTAGAACTCTGAACCCCTATAGTGAGTCGTATTACCGGGAGCTT |
| ABID_SBC_bot_184 | GTTCATCA | /5Phos/GGCCCTGTTCNNNATCANNNGATCGTCGGACTGTAGAACTCTGAACCCCTATAGTGAGTCGTATTACCGGGAGCTT |
| ABID_SBC_bot_185 | TTATACGC | /5Phos/GGCCCTTTATNNNACGCNNNGATCGTCGGACTGTAGAACTCTGAACCCCTATAGTGAGTCGTATTACCGGGAGCTT |
| ABID_SBC_bot_186 | AGCTTCGT | /5Phos/GGCCCTAGCTNNNTCGTNNNGATCGTCGGACTGTAGAACTCTGAACCCCTATAGTGAGTCGTATTACCGGGAGCTT |
| ABID_SBC_bot_187 | GTACGCAC | /5Phos/GGCCCTGTACNNNGCACNNNGATCGTCGGACTGTAGAACTCTGAACCCCTATAGTGAGTCGTATTACCGGGAGCTT |
| ABID_SBC_bot_188 | TCACTAGA | /5Phos/GGCCCTTCACNNNTAGANNNGATCGTCGGACTGTAGAACTCTGAACCCCTATAGTGAGTCGTATTACCGGGAGCTT |
| ABID_SBC_bot_189 | GTGCTCGA | /5Phos/GGCCCTGTGCNNNTCGANNNGATCGTCGGACTGTAGAACTCTGAACCCCTATAGTGAGTCGTATTACCGGGAGCTT |
| ABID_SBC_bot_190 | GACGTACA | /5Phos/GGCCCTGACGNNNTACANNNGATCGTCGGACTGTAGAACTCTGAACCCCTATAGTGAGTCGTATTACCGGGAGCTT |
| ABID_SBC_bot_191 | GTAGACTC | /5Phos/GGCCCTGTAGNNNACTCNNNGATCGTCGGACTGTAGAACTCTGAACCCCTATAGTGAGTCGTATTACCGGGAGCTT |
| ABID_SBC_bot_192 | TGACACTC | /5Phos/GGCCCTTGACNNNACTCNNNGATCGTCGGACTGTAGAACTCTGAACCCCTATAGTGAGTCGTATTACCGGGAGCTT |
| ABID_SBC_bot_193 | AGTCTCAC | /5Phos/GGCCCTAGTCNNNTCACNNNGATCGTCGGACTGTAGAACTCTGAACCCCTATAGTGAGTCGTATTACCGGGAGCTT |
| ABID_SBC_bot_194 | GTTGCAGC | /5Phos/GGCCCTGTTGNNNCAGCNNNGATCGTCGGACTGTAGAACTCTGAACCCCTATAGTGAGTCGTATTACCGGGAGCTT |
| ABID_SBC_bot_195 | CTGCTGCA | /5Phos/GGCCCTCTGCNNNTGCANNNGATCGTCGGACTGTAGAACTCTGAACCCCTATAGTGAGTCGTATTACCGGGAGCTT |
| ABID_SBC_bot_196 | TGGTCTAC | /5Phos/GGCCCTTGGTNNNCTACNNNGATCGTCGGACTGTAGAACTCTGAACCCCTATAGTGAGTCGTATTACCGGGAGCTT |
| ABID_SBC_bot_197 | AGGCGTAC | /5Phos/GGCCCTAGGCNNNGTACNNNGATCGTCGGACTGTAGAACTCTGAACCCCTATAGTGAGTCGTATTACCGGGAGCTT |
| ABID_SBC_bot_198 | GAGCCTTC | /5Phos/GGCCCTGAGCNNNCTTCNNNGATCGTCGGACTGTAGAACTCTGAACCCCTATAGTGAGTCGTATTACCGGGAGCTT |
| ABID_SBC_bot_199 | GGTCGAGA | /5Phos/GGCCCTGGTCNNNGAGANNNGATCGTCGGACTGTAGAACTCTGAACCCCTATAGTGAGTCGTATTACCGGGAGCTT |
| ABID_SBC_bot_200 | TCGTAGAG | /5Phos/GGCCCTTCGTNNNAGAGNNNGATCGTCGGACTGTAGAACTCTGAACCCCTATAGTGAGTCGTATTACCGGGAGCTT |
| ABID_SBC_bot_201 | GTCGTGAC | /5Phos/GGCCCTGTCGNNNTGACNNNGATCGTCGGACTGTAGAACTCTGAACCCCTATAGTGAGTCGTATTACCGGGAGCTT |
| ABID_SBC_bot_202 | AAGCGACA | /5Phos/GGCCCTAAGCNNNGACANNNGATCGTCGGACTGTAGAACTCTGAACCCCTATAGTGAGTCGTATTACCGGGAGCTT |
| ABID_SBC_bot_203 | TGAGCATG | /5Phos/GGCCCTTGAGNNNCATGNNNGATCGTCGGACTGTAGAACTCTGAACCCCTATAGTGAGTCGTATTACCGGGAGCTT |
| ABID_SBC_bot_204 | TGCGAGTC | /5Phos/GGCCCTTGCGNNNAGTCNNNGATCGTCGGACTGTAGAACTCTGAACCCCTATAGTGAGTCGTATTACCGGGAGCTT |
| ABID_SBC_bot_205 | GGTGCACA | /5Phos/GGCCCTGGTGNNNCACANNNGATCGTCGGACTGTAGAACTCTGAACCCCTATAGTGAGTCGTATTACCGGGAGCTT |
| ABID_SBC_bot_206 | TCTGCTAC | /5Phos/GGCCCTTCTGNNNCTACNNNGATCGTCGGACTGTAGAACTCTGAACCCCTATAGTGAGTCGTATTACCGGGAGCTT |
| ABID_SBC_bot_207 | GTGTACAG | /5Phos/GGCCCTGTGTNNNACAGNNNGATCGTCGGACTGTAGAACTCTGAACCCCTATAGTGAGTCGTATTACCGGGAGCTT |
| ABID_SBC_bot_208 | GCGTCAAG | /5Phos/GGCCCTGCGTNNNCAAGNNNGATCGTCGGACTGTAGAACTCTGAACCCCTATAGTGAGTCGTATTACCGGGAGCTT |

| | | |
|---|---|---|
| ABID_SBC_bot_209 | AACGCGCT | /5Phos/GGCCCTAACGNNNCGCTNNNGATCGTCGGACTGTAGAACTCTGAACCCCTATAGTGAGTCGTATTACCGGGAGCTT |
| ABID_SBC_bot_210 | CTCGGTGT | /5Phos/GGCCCTCTCGNNNGTGTNNNGATCGTCGGACTGTAGAACTCTGAACCCCTATAGTGAGTCGTATTACCGGGAGCTT |
| ABID_SBC_bot_211 | ACTCCAGA | /5Phos/GGCCCTACTCNNNCAGANNNGATCGTCGGACTGTAGAACTCTGAACCCCTATAGTGAGTCGTATTACCGGGAGCTT |
| ABID_SBC_bot_212 | CTTGAGCG | /5Phos/GGCCCTCTTGNNNAGCGNNNGATCGTCGGACTGTAGAACTCTGAACCCCTATAGTGAGTCGTATTACCGGGAGCTT |
| ABID_SBC_bot_213 | ACACTCCT | /5Phos/GGCCCTACACNNNTCCTNNNGATCGTCGGACTGTAGAACTCTGAACCCCTATAGTGAGTCGTATTACCGGGAGCTT |
| ABID_SBC_bot_214 | GAAGGTGC | /5Phos/GGCCCTGAAGNNNGTGCNNNGATCGTCGGACTGTAGAACTCTGAACCCCTATAGTGAGTCGTATTACCGGGAGCTT |
| ABID_SBC_bot_215 | ACGAATCT | /5Phos/GGCCCTACGANNNATCTNNNGATCGTCGGACTGTAGAACTCTGAACCCCTATAGTGAGTCGTATTACCGGGAGCTT |
| ABID_SBC_bot_216 | ACATCACA | /5Phos/GGCCCTACATNNNCACANNNGATCGTCGGACTGTAGAACTCTGAACCCCTATAGTGAGTCGTATTACCGGGAGCTT |
| ABID_SBC_bot_217 | TCTCGCAG | /5Phos/GGCCCTTCTCNNNGCAGNNNGATCGTCGGACTGTAGAACTCTGAACCCCTATAGTGAGTCGTATTACCGGGAGCTT |
| ABID_SBC_bot_218 | AGCTCGCA | /5Phos/GGCCCTAGCTNNNCGCANNNGATCGTCGGACTGTAGAACTCTGAACCCCTATAGTGAGTCGTATTACCGGGAGCTT |
| ABID_SBC_bot_219 | CTCTCTAT | /5Phos/GGCCCTCTCTNNNCTATNNNGATCGTCGGACTGTAGAACTCTGAACCCCTATAGTGAGTCGTATTACCGGGAGCTT |
| ABID_SBC_bot_220 | GCTAGCAT | /5Phos/GGCCCTGCTANNNGCATNNNGATCGTCGGACTGTAGAACTCTGAACCCCTATAGTGAGTCGTATTACCGGGAGCTT |
| ABID_SBC_bot_221 | CAGTACGC | /5Phos/GGCCCTCAGTNNNACGCNNNGATCGTCGGACTGTAGAACTCTGAACCCCTATAGTGAGTCGTATTACCGGGAGCTT |
| ABID_SBC_bot_222 | ACCACACG | /5Phos/GGCCCTACCANNNCACGNNNGATCGTCGGACTGTAGAACTCTGAACCCCTATAGTGAGTCGTATTACCGGGAGCTT |
| ABID_SBC_bot_223 | TAGTCGAT | /5Phos/GGCCCTTAGTNNNCGATNNNGATCGTCGGACTGTAGAACTCTGAACCCCTATAGTGAGTCGTATTACCGGGAGCTT |
| ABID_SBC_bot_224 | TAACCTGT | /5Phos/GGCCCTTAACNNNCTGTNNNGATCGTCGGACTGTAGAACTCTGAACCCCTATAGTGAGTCGTATTACCGGGAGCTT |
| ABID_SBC_bot_225 | GCTAAGCG | /5Phos/GGCCCTGCTANNNAGCGNNNGATCGTCGGACTGTAGAACTCTGAACCCCTATAGTGAGTCGTATTACCGGGAGCTT |
| ABID_SBC_bot_226 | TGGCGAAG | /5Phos/GGCCCTTGGCNNNGAAGNNNGATCGTCGGACTGTAGAACTCTGAACCCCTATAGTGAGTCGTATTACCGGGAGCTT |
| ABID_SBC_bot_227 | TGATTCGA | /5Phos/GGCCCTTGATNNNTCGANNNGATCGTCGGACTGTAGAACTCTGAACCCCTATAGTGAGTCGTATTACCGGGAGCTT |
| ABID_SBC_bot_228 | ATCGTCCT | /5Phos/GGCCCTATCGNNNTCCTNNNGATCGTCGGACTGTAGAACTCTGAACCCCTATAGTGAGTCGTATTACCGGGAGCTT |
| ABID_SBC_bot_229 | GCATCGCT | /5Phos/GGCCCTGCATNNNCGCTNNNGATCGTCGGACTGTAGAACTCTGAACCCCTATAGTGAGTCGTATTACCGGGAGCTT |
| ABID_SBC_bot_230 | GCGAGAGA | /5Phos/GGCCCTGCGANNNGAGANNNGATCGTCGGACTGTAGAACTCTGAACCCCTATAGTGAGTCGTATTACCGGGAGCTT |
| ABID_SBC_bot_231 | TGGACAGC | /5Phos/GGCCCTTGGANNNCAGCNNNGATCGTCGGACTGTAGAACTCTGAACCCCTATAGTGAGTCGTATTACCGGGAGCTT |
| ABID_SBC_bot_232 | GTCAGCTC | /5Phos/GGCCCTGTCANNNGCTCNNNGATCGTCGGACTGTAGAACTCTGAACCCCTATAGTGAGTCGTATTACCGGGAGCTT |
| ABID_SBC_bot_233 | ACATGAAG | /5Phos/GGCCCTACATNNNGAAGNNNGATCGTCGGACTGTAGAACTCTGAACCCCTATAGTGAGTCGTATTACCGGGAGCTT |
| ABID_SBC_bot_234 | CGAGTATA | /5Phos/GGCCCTCGAGNNNTATANNNGATCGTCGGACTGTAGAACTCTGAACCCCTATAGTGAGTCGTATTACCGGGAGCTT |
| ABID_SBC_bot_235 | CATAAGGA | /5Phos/GGCCCTCATANNNAGGANNNGATCGTCGGACTGTAGAACTCTGAACCCCTATAGTGAGTCGTATTACCGGGAGCTT |
| ABID_SBC_bot_236 | CTTAACCT | /5Phos/GGCCCTCTTANNNACCTNNNGATCGTCGGACTGTAGAACTCTGAACCCCTATAGTGAGTCGTATTACCGGGAGCTT |
| ABID_SBC_bot_237 | TACTTGTC | /5Phos/GGCCCTTACTNNNTGTCNNNGATCGTCGGACTGTAGAACTCTGAACCCCTATAGTGAGTCGTATTACCGGGAGCTT |
| ABID_SBC_bot_238 | GAACGTCA | /5Phos/GGCCCTGAACNNNGTCANNNGATCGTCGGACTGTAGAACTCTGAACCCCTATAGTGAGTCGTATTACCGGGAGCTT |
| ABID_SBC_bot_239 | GATATCAC | /5Phos/GGCCCTGATANNNTCACNNNGATCGTCGGACTGTAGAACTCTGAACCCCTATAGTGAGTCGTATTACCGGGAGCTT |
| ABID_SBC_bot_240 | TTGCTAAC | /5Phos/GGCCCTTTGCNNNTAACNNNGATCGTCGGACTGTAGAACTCTGAACCCCTATAGTGAGTCGTATTACCGGGAGCTT |
| ABID_SBC_bot_241 | GACAATGA | /5Phos/GGCCCTGACANNNATGANNNGATCGTCGGACTGTAGAACTCTGAACCCCTATAGTGAGTCGTATTACCGGGAGCTT |

| | | |
|---|---|---|
| ABID_SBC_bot_242 | ATGAACCA | /5Phos/GGCCCTATGANNNACCANNNGATCGTCGGACTGTAGAACTCTGAACCCCTATAGTGAGTCGTATTACCGGGAGCTT |
| ABID_SBC_bot_243 | AACGATTG | /5Phos/GGCCCTAACGNNNATTGNNNGATCGTCGGACTGTAGAACTCTGAACCCCTATAGTGAGTCGTATTACCGGGAGCTT |
| ABID_SBC_bot_244 | CATCGTAT | /5Phos/GGCCCTCATCNNNGTATNNNGATCGTCGGACTGTAGAACTCTGAACCCCTATAGTGAGTCGTATTACCGGGAGCTT |
| ABID_SBC_bot_245 | CTCTACTA | /5Phos/GGCCCTCTCTNNNACTANNNGATCGTCGGACTGTAGAACTCTGAACCCCTATAGTGAGTCGTATTACCGGGAGCTT |
| ABID_SBC_bot_246 | TCGCTGAT | /5Phos/GGCCCTTCGCNNNTGATNNNGATCGTCGGACTGTAGAACTCTGAACCCCTATAGTGAGTCGTATTACCGGGAGCTT |
| ABID_SBC_bot_247 | CTTCCGGA | /5Phos/GGCCCTCTTCNNNCGGANNNGATCGTCGGACTGTAGAACTCTGAACCCCTATAGTGAGTCGTATTACCGGGAGCTT |
| ABID_SBC_bot_248 | TTGCACGT | /5Phos/GGCCCTTTGCNNNACGTNNNGATCGTCGGACTGTAGAACTCTGAACCCCTATAGTGAGTCGTATTACCGGGAGCTT |
| ABID_SBC_bot_249 | AGGACGTA | /5Phos/GGCCCTAGGANNNCGTANNNGATCGTCGGACTGTAGAACTCTGAACCCCTATAGTGAGTCGTATTACCGGGAGCTT |
| ABID_SBC_bot_250 | CTCGAGGA | /5Phos/GGCCCTCTCGNNNAGGANNNGATCGTCGGACTGTAGAACTCTGAACCCCTATAGTGAGTCGTATTACCGGGAGCTT |
| ABID_SBC_bot_251 | CTGCAGAG | /5Phos/GGCCCTCTGCNNNAGAGNNNGATCGTCGGACTGTAGAACTCTGAACCCCTATAGTGAGTCGTATTACCGGGAGCTT |
| ABID_SBC_bot_252 | TCAGACGT | /5Phos/GGCCCTTCAGNNNACGTNNNGATCGTCGGACTGTAGAACTCTGAACCCCTATAGTGAGTCGTATTACCGGGAGCTT |
| ABID_SBC_bot_253 | TCCTGTCG | /5Phos/GGCCCTTCCTNNNGTCGNNNGATCGTCGGACTGTAGAACTCTGAACCCCTATAGTGAGTCGTATTACCGGGAGCTT |
| ABID_SBC_bot_254 | TTCTCGAC | /5Phos/GGCCCTTTCTNNNCGACNNNGATCGTCGGACTGTAGAACTCTGAACCCCTATAGTGAGTCGTATTACCGGGAGCTT |
| ABID_SBC_bot_255 | AGATCAGT | /5Phos/GGCCCTAGATNNNCAGTNNNGATCGTCGGACTGTAGAACTCTGAACCCCTATAGTGAGTCGTATTACCGGGAGCTT |
| ABID_SBC_bot_256 | CAGCTGGT | /5Phos/GGCCCTCAGCNNNTGGTNNNGATCGTCGGACTGTAGAACTCTGAACCCCTATAGTGAGTCGTATTACCGGGAGCTT |
| ABID_SBC_bot_257 | GATCATAG | /5Phos/GGCCCTGATCNNNATAGNNNGATCGTCGGACTGTAGAACTCTGAACCCCTATAGTGAGTCGTATTACCGGGAGCTT |
| ABID_SBC_bot_258 | GCACAGAT | /5Phos/GGCCCTGCACNNNAGATNNNGATCGTCGGACTGTAGAACTCTGAACCCCTATAGTGAGTCGTATTACCGGGAGCTT |
| ABID_SBC_bot_259 | CGGCTATC | /5Phos/GGCCCTCGGCNNNTATCNNNGATCGTCGGACTGTAGAACTCTGAACCCCTATAGTGAGTCGTATTACCGGGAGCTT |
| ABID_SBC_bot_260 | AGGTGATC | /5Phos/GGCCCTAGGTNNNGATCNNNGATCGTCGGACTGTAGAACTCTGAACCCCTATAGTGAGTCGTATTACCGGGAGCTT |
| ABID_SBC_bot_261 | ACATCGTC | /5Phos/GGCCCTACATNNNCGTCNNNGATCGTCGGACTGTAGAACTCTGAACCCCTATAGTGAGTCGTATTACCGGGAGCTT |
| ABID_SBC_bot_262 | AGAGTGAT | /5Phos/GGCCCTAGAGNNNTGATNNNGATCGTCGGACTGTAGAACTCTGAACCCCTATAGTGAGTCGTATTACCGGGAGCTT |
| ABID_SBC_bot_263 | ATTCACTG | /5Phos/GGCCCTATTCNNNACTGNNNGATCGTCGGACTGTAGAACTCTGAACCCCTATAGTGAGTCGTATTACCGGGAGCTT |
| ABID_SBC_bot_264 | TGTCGATC | /5Phos/GGCCCTTGTCNNNGATCNNNGATCGTCGGACTGTAGAACTCTGAACCCCTATAGTGAGTCGTATTACCGGGAGCTT |
| ABID_SBC_bot_265 | CTTGTCTG | /5Phos/GGCCCTCTTGNNNTCTGNNNGATCGTCGGACTGTAGAACTCTGAACCCCTATAGTGAGTCGTATTACCGGGAGCTT |
| ABID_SBC_bot_266 | TAAGGCGA | /5Phos/GGCCCTTAAGNNNGCGANNNGATCGTCGGACTGTAGAACTCTGAACCCCTATAGTGAGTCGTATTACCGGGAGCTT |
| ABID_SBC_bot_267 | CGATGCAC | /5Phos/GGCCCTCGATNNNGCACNNNGATCGTCGGACTGTAGAACTCTGAACCCCTATAGTGAGTCGTATTACCGGGAGCTT |
| ABID_SBC_bot_268 | TTACCTAC | /5Phos/GGCCCTTTACNNNCTACNNNGATCGTCGGACTGTAGAACTCTGAACCCCTATAGTGAGTCGTATTACCGGGAGCTT |
| ABID_SBC_bot_269 | GATAGCCA | /5Phos/GGCCCTGATANNNGCCANNNGATCGTCGGACTGTAGAACTCTGAACCCCTATAGTGAGTCGTATTACCGGGAGCTT |
| ABID_SBC_bot_270 | AGTGACGA | /5Phos/GGCCCTAGTGNNNACGANNNGATCGTCGGACTGTAGAACTCTGAACCCCTATAGTGAGTCGTATTACCGGGAGCTT |
| ABID_SBC_bot_271 | AGGAAGAT | /5Phos/GGCCCTAGGANNNAGATNNNGATCGTCGGACTGTAGAACTCTGAACCCCTATAGTGAGTCGTATTACCGGGAGCTT |
| ABID_SBC_bot_272 | GACGACAC | /5Phos/GGCCCTGACGNNNACACNNNGATCGTCGGACTGTAGAACTCTGAACCCCTATAGTGAGTCGTATTACCGGGAGCTT |
| ABID_SBC_bot_273 | GCCACTCA | /5Phos/GGCCCTGCCANNNCTCANNNGATCGTCGGACTGTAGAACTCTGAACCCCTATAGTGAGTCGTATTACCGGGAGCTT |
| ABID_SBC_bot_274 | ACAGCTCG | /5Phos/GGCCCTACAGNNNCTCGNNNGATCGTCGGACTGTAGAACTCTGAACCCCTATAGTGAGTCGTATTACCGGGAGCTT |

| | | |
|---|---|---|
| ABID_SBC_bot_275 | CATATCGT | /5Phos/GGCCCTCATANNNTCGTNNNGATCGTCGGACTGTAGAACTCTGAACCCCTATAGTGAGTCGTATTACCGGGAGCTT |
| ABID_SBC_bot_276 | AGCAACCG | /5Phos/GGCCCTAGCANNNACCGNNNGATCGTCGGACTGTAGAACTCTGAACCCCTATAGTGAGTCGTATTACCGGGAGCTT |
| ABID_SBC_bot_277 | GTCAAGGC | /5Phos/GGCCCTGTCANNNAGGCNNNGATCGTCGGACTGTAGAACTCTGAACCCCTATAGTGAGTCGTATTACCGGGAGCTT |
| ABID_SBC_bot_278 | CTGCGAGA | /5Phos/GGCCCTCTGCNNNGAGANNNGATCGTCGGACTGTAGAACTCTGAACCCCTATAGTGAGTCGTATTACCGGGAGCTT |
| ABID_SBC_bot_279 | AGGAGCCT | /5Phos/GGCCCTAGGANNNGCCTNNNGATCGTCGGACTGTAGAACTCTGAACCCCTATAGTGAGTCGTATTACCGGGAGCTT |
| ABID_SBC_bot_280 | TCTGCATA | /5Phos/GGCCCTTCTGNNNCATANNNGATCGTCGGACTGTAGAACTCTGAACCCCTATAGTGAGTCGTATTACCGGGAGCTT |
| ABID_SBC_bot_281 | AATGACTC | /5Phos/GGCCCTAATGNNNACTCNNNGATCGTCGGACTGTAGAACTCTGAACCCCTATAGTGAGTCGTATTACCGGGAGCTT |
| ABID_SBC_bot_282 | AAGATCGC | /5Phos/GGCCCTAAGANNNTCGCNNNGATCGTCGGACTGTAGAACTCTGAACCCCTATAGTGAGTCGTATTACCGGGAGCTT |
| ABID_SBC_bot_283 | GAAGATCT | /5Phos/GGCCCTGAAGNNNATCTNNNGATCGTCGGACTGTAGAACTCTGAACCCCTATAGTGAGTCGTATTACCGGGAGCTT |
| ABID_SBC_bot_284 | CGGATGCT | /5Phos/GGCCCTCGGANNNTGCTNNNGATCGTCGGACTGTAGAACTCTGAACCCCTATAGTGAGTCGTATTACCGGGAGCTT |
| ABID_SBC_bot_285 | ATGACAGA | /5Phos/GGCCCTATGANNNCAGANNNGATCGTCGGACTGTAGAACTCTGAACCCCTATAGTGAGTCGTATTACCGGGAGCTT |
| ABID_SBC_bot_286 | ACTGACCG | /5Phos/GGCCCTACTGNNNACCGNNNGATCGTCGGACTGTAGAACTCTGAACCCCTATAGTGAGTCGTATTACCGGGAGCTT |
| ABID_SBC_bot_287 | TAGACTAG | /5Phos/GGCCCTTAGANNNCTAGNNNGATCGTCGGACTGTAGAACTCTGAACCCCTATAGTGAGTCGTATTACCGGGAGCTT |
| ABID_SBC_bot_288 | GTATCGGC | /5Phos/GGCCCTGTATNNNCGGCNNNGATCGTCGGACTGTAGAACTCTGAACCCCTATAGTGAGTCGTATTACCGGGAGCTT |
| ABID_SBC_bot_289 | GAATAGTC | /5Phos/GGCCCTGAATNNNAGTCNNNGATCGTCGGACTGTAGAACTCTGAACCCCTATAGTGAGTCGTATTACCGGGAGCTT |
| ABID_SBC_bot_290 | CACTGACA | /5Phos/GGCCCTCACTNNNGACANNNGATCGTCGGACTGTAGAACTCTGAACCCCTATAGTGAGTCGTATTACCGGGAGCTT |
| ABID_SBC_bot_291 | TGTCACAT | /5Phos/GGCCCTTGTCNNNACATNNNGATCGTCGGACTGTAGAACTCTGAACCCCTATAGTGAGTCGTATTACCGGGAGCTT |
| ABID_SBC_bot_292 | TACTACCT | /5Phos/GGCCCTTACTNNNACCTNNNGATCGTCGGACTGTAGAACTCTGAACCCCTATAGTGAGTCGTATTACCGGGAGCTT |
| ABID_SBC_bot_293 | GCCTCGTA | /5Phos/GGCCCTGCCTNNNCGTANNNGATCGTCGGACTGTAGAACTCTGAACCCCTATAGTGAGTCGTATTACCGGGAGCTT |
| ABID_SBC_bot_294 | CAGTTGAG | /5Phos/GGCCCTCAGTNNNTGAGNNNGATCGTCGGACTGTAGAACTCTGAACCCCTATAGTGAGTCGTATTACCGGGAGCTT |
| ABID_SBC_bot_295 | AGCATATG | /5Phos/GGCCCTAGCANNNTATGNNNGATCGTCGGACTGTAGAACTCTGAACCCCTATAGTGAGTCGTATTACCGGGAGCTT |
| ABID_SBC_bot_296 | AGGAATGA | /5Phos/GGCCCTAGGANNNATGANNNGATCGTCGGACTGTAGAACTCTGAACCCCTATAGTGAGTCGTATTACCGGGAGCTT |
| ABID_SBC_bot_297 | CGTACTTA | /5Phos/GGCCCTCGTANNNCTTANNNGATCGTCGGACTGTAGAACTCTGAACCCCTATAGTGAGTCGTATTACCGGGAGCTT |
| ABID_SBC_bot_298 | GTCGCTAG | /5Phos/GGCCCTGTCGNNNCTAGNNNGATCGTCGGACTGTAGAACTCTGAACCCCTATAGTGAGTCGTATTACCGGGAGCTT |
| ABID_SBC_bot_299 | GTACACCG | /5Phos/GGCCCTGTACNNNACCGNNNGATCGTCGGACTGTAGAACTCTGAACCCCTATAGTGAGTCGTATTACCGGGAGCTT |
| ABID_SBC_bot_300 | ACGTCTGA | /5Phos/GGCCCTACGTNNNCTGANNNGATCGTCGGACTGTAGAACTCTGAACCCCTATAGTGAGTCGTATTACCGGGAGCTT |
| ABID_SBC_bot_301 | AACGTGTA | /5Phos/GGCCCTAACGNNNTGTANNNGATCGTCGGACTGTAGAACTCTGAACCCCTATAGTGAGTCGTATTACCGGGAGCTT |
| ABID_SBC_bot_302 | CACGCATA | /5Phos/GGCCCTCACGNNNCATANNNGATCGTCGGACTGTAGAACTCTGAACCCCTATAGTGAGTCGTATTACCGGGAGCTT |
| ABID_SBC_bot_303 | ACGATGGA | /5Phos/GGCCCTACGANNNTGGANNNGATCGTCGGACTGTAGAACTCTGAACCCCTATAGTGAGTCGTATTACCGGGAGCTT |
| ABID_SBC_bot_304 | TGCTGATA | /5Phos/GGCCCTTGCTNNNGATANNNGATCGTCGGACTGTAGAACTCTGAACCCCTATAGTGAGTCGTATTACCGGGAGCTT |
| ABID_SBC_bot_305 | GTACCACA | /5Phos/GGCCCTGTACNNNCACANNNGATCGTCGGACTGTAGAACTCTGAACCCCTATAGTGAGTCGTATTACCGGGAGCTT |
| ABID_SBC_bot_306 | AGGAACTC | /5Phos/GGCCCTAGGANNNACTCNNNGATCGTCGGACTGTAGAACTCTGAACCCCTATAGTGAGTCGTATTACCGGGAGCTT |
| ABID_SBC_bot_307 | CTGTTCTC | /5Phos/GGCCCTCTGTNNNTCTCNNNGATCGTCGGACTGTAGAACTCTGAACCCCTATAGTGAGTCGTATTACCGGGAGCTT |

| | | |
|---|---|---|
| ABID_SBC_bot_308 | TCAGCACT | /5Phos/GGCCCTTCAGNNNCACTNNNGATCGTCGGACTGTAGAACTCTGAACCCCTATAGTGAGTCGTATTACCGGGAGCTT |
| ABID_SBC_bot_309 | ATCGCGTC | /5Phos/GGCCCTATCGNNNCGTCNNNGATCGTCGGACTGTAGAACTCTGAACCCCTATAGTGAGTCGTATTACCGGGAGCTT |
| ABID_SBC_bot_310 | CATGTATC | /5Phos/GGCCCTCATGNNNTATCNNNGATCGTCGGACTGTAGAACTCTGAACCCCTATAGTGAGTCGTATTACCGGGAGCTT |
| ABID_SBC_bot_311 | GTGTTCCT | /5Phos/GGCCCTGTGTNNNTCCTNNNGATCGTCGGACTGTAGAACTCTGAACCCCTATAGTGAGTCGTATTACCGGGAGCTT |
| ABID_SBC_bot_312 | ATGAAGTG | /5Phos/GGCCCTATGANNNAGTGNNNGATCGTCGGACTGTAGAACTCTGAACCCCTATAGTGAGTCGTATTACCGGGAGCTT |
| ABID_SBC_bot_313 | TTCGTCTA | /5Phos/GGCCCTTTCGNNNTCTANNNGATCGTCGGACTGTAGAACTCTGAACCCCTATAGTGAGTCGTATTACCGGGAGCTT |
| ABID_SBC_bot_314 | GCATATCA | /5Phos/GGCCCTGCATNNNATCANNNGATCGTCGGACTGTAGAACTCTGAACCCCTATAGTGAGTCGTATTACCGGGAGCTT |
| ABID_SBC_bot_315 | TGCAATAG | /5Phos/GGCCCTTGCANNNATAGNNNGATCGTCGGACTGTAGAACTCTGAACCCCTATAGTGAGTCGTATTACCGGGAGCTT |
| ABID_SBC_bot_316 | TACAACGC | /5Phos/GGCCCTTACANNNACGCNNNGATCGTCGGACTGTAGAACTCTGAACCCCTATAGTGAGTCGTATTACCGGGAGCTT |
| ABID_SBC_bot_317 | GGTCTCTG | /5Phos/GGCCCTGGTCNNNTCTGNNNGATCGTCGGACTGTAGAACTCTGAACCCCTATAGTGAGTCGTATTACCGGGAGCTT |
| ABID_SBC_bot_318 | TGGTAGGT | /5Phos/GGCCCTTGGTNNNAGGTNNNGATCGTCGGACTGTAGAACTCTGAACCCCTATAGTGAGTCGTATTACCGGGAGCTT |
| ABID_SBC_bot_319 | AATCGAAG | /5Phos/GGCCCTAATCNNNGAAGNNNGATCGTCGGACTGTAGAACTCTGAACCCCTATAGTGAGTCGTATTACCGGGAGCTT |
| ABID_SBC_bot_320 | GGATCTCG | /5Phos/GGCCCTGGATNNNCTCGNNNGATCGTCGGACTGTAGAACTCTGAACCCCTATAGTGAGTCGTATTACCGGGAGCTT |
| ABID_SBC_bot_321 | TAGCACAG | /5Phos/GGCCCTTAGCNNNACAGNNNGATCGTCGGACTGTAGAACTCTGAACCCCTATAGTGAGTCGTATTACCGGGAGCTT |
| ABID_SBC_bot_322 | TATCGCTA | /5Phos/GGCCCTTATCNNNGCTANNNGATCGTCGGACTGTAGAACTCTGAACCCCTATAGTGAGTCGTATTACCGGGAGCTT |
| ABID_SBC_bot_323 | TCTAGAAC | /5Phos/GGCCCTTCTANNNGAACNNNGATCGTCGGACTGTAGAACTCTGAACCCCTATAGTGAGTCGTATTACCGGGAGCTT |
| ABID_SBC_bot_324 | AAGCCGAG | /5Phos/GGCCCTAAGCNNNCGAGNNNGATCGTCGGACTGTAGAACTCTGAACCCCTATAGTGAGTCGTATTACCGGGAGCTT |
| ABID_SBC_bot_325 | GGATACTA | /5Phos/GGCCCTGGATNNNACTANNNGATCGTCGGACTGTAGAACTCTGAACCCCTATAGTGAGTCGTATTACCGGGAGCTT |
| ABID_SBC_bot_326 | ATTGCTGA | /5Phos/GGCCCTATTGNNNCTGANNNGATCGTCGGACTGTAGAACTCTGAACCCCTATAGTGAGTCGTATTACCGGGAGCTT |
| ABID_SBC_bot_327 | TTAGCAGA | /5Phos/GGCCCTTTAGNNNCAGANNNGATCGTCGGACTGTAGAACTCTGAACCCCTATAGTGAGTCGTATTACCGGGAGCTT |
| ABID_SBC_bot_328 | CGTATCCG | /5Phos/GGCCCTCGTANNNTCCGNNNGATCGTCGGACTGTAGAACTCTGAACCCCTATAGTGAGTCGTATTACCGGGAGCTT |
| ABID_SBC_bot_329 | TCTCCACG | /5Phos/GGCCCTTCTCNNNCACGNNNGATCGTCGGACTGTAGAACTCTGAACCCCTATAGTGAGTCGTATTACCGGGAGCTT |
| ABID_SBC_bot_330 | GTGTATGA | /5Phos/GGCCCTGTGTNNNATGANNNGATCGTCGGACTGTAGAACTCTGAACCCCTATAGTGAGTCGTATTACCGGGAGCTT |
| ABID_SBC_bot_331 | ACCTTCCG | /5Phos/GGCCCTACCTNNNTCCGNNNGATCGTCGGACTGTAGAACTCTGAACCCCTATAGTGAGTCGTATTACCGGGAGCTT |
| ABID_SBC_bot_332 | ACTGTAAC | /5Phos/GGCCCTACTGNNNTAACNNNGATCGTCGGACTGTAGAACTCTGAACCCCTATAGTGAGTCGTATTACCGGGAGCTT |
| ABID_SBC_bot_333 | GGTCAGAC | /5Phos/GGCCCTGGTCNNNAGACNNNGATCGTCGGACTGTAGAACTCTGAACCCCTATAGTGAGTCGTATTACCGGGAGCTT |
| ABID_SBC_bot_334 | TTGAAGAC | /5Phos/GGCCCTTTGANNNAGACNNNGATCGTCGGACTGTAGAACTCTGAACCCCTATAGTGAGTCGTATTACCGGGAGCTT |
| ABID_SBC_bot_335 | AGCAGAAC | /5Phos/GGCCCTAGCANNNGAACNNNGATCGTCGGACTGTAGAACTCTGAACCCCTATAGTGAGTCGTATTACCGGGAGCTT |
| ABID_SBC_bot_336 | TGTGCGGA | /5Phos/GGCCCTTGTGNNNCGGANNNGATCGTCGGACTGTAGAACTCTGAACCCCTATAGTGAGTCGTATTACCGGGAGCTT |
| ABID_SBC_bot_337 | ACGAGAAT | /5Phos/GGCCCTACGANNNGAATNNNGATCGTCGGACTGTAGAACTCTGAACCCCTATAGTGAGTCGTATTACCGGGAGCTT |
| ABID_SBC_bot_338 | CGGACACA | /5Phos/GGCCCTCGGANNNCACANNNGATCGTCGGACTGTAGAACTCTGAACCCCTATAGTGAGTCGTATTACCGGGAGCTT |
| ABID_SBC_bot_339 | CACATGAC | /5Phos/GGCCCTCACANNNTGACNNNGATCGTCGGACTGTAGAACTCTGAACCCCTATAGTGAGTCGTATTACCGGGAGCTT |
| ABID_SBC_bot_340 | ATTAGCAC | /5Phos/GGCCCTATTANNNGCACNNNGATCGTCGGACTGTAGAACTCTGAACCCCTATAGTGAGTCGTATTACCGGGAGCTT |

| | | |
|---|---|---|
| ABID_SBC_bot_341 | GCGATCTC | /5Phos/GGCCCTGCGANNNTCTCNNNGATCGTCGGACTGTAGAACTCTGAACCCCTATAGTGAGTCGTATTACCGGGAGCTT |
| ABID_SBC_bot_342 | TAGCAGGC | /5Phos/GGCCCTTAGCNNNAGGCNNNGATCGTCGGACTGTAGAACTCTGAACCCCTATAGTGAGTCGTATTACCGGGAGCTT |
| ABID_SBC_bot_343 | AGATTCTC | /5Phos/GGCCCTAGATNNNTCTCNNNGATCGTCGGACTGTAGAACTCTGAACCCCTATAGTGAGTCGTATTACCGGGAGCTT |
| ABID_SBC_bot_344 | ACAGGCTG | /5Phos/GGCCCTACAGNNNGCTGNNNGATCGTCGGACTGTAGAACTCTGAACCCCTATAGTGAGTCGTATTACCGGGAGCTT |
| ABID_SBC_bot_345 | ACATACAC | /5Phos/GGCCCTACATNNNACACNNNGATCGTCGGACTGTAGAACTCTGAACCCCTATAGTGAGTCGTATTACCGGGAGCTT |
| ABID_SBC_bot_346 | CACAGTCT | /5Phos/GGCCCTCACANNNGTCTNNNGATCGTCGGACTGTAGAACTCTGAACCCCTATAGTGAGTCGTATTACCGGGAGCTT |
| ABID_SBC_bot_347 | GACATATC | /5Phos/GGCCCTGACANNNTATCNNNGATCGTCGGACTGTAGAACTCTGAACCCCTATAGTGAGTCGTATTACCGGGAGCTT |
| ABID_SBC_bot_348 | GGAGTCCA | /5Phos/GGCCCTGGAGNNNTCCANNNGATCGTCGGACTGTAGAACTCTGAACCCCTATAGTGAGTCGTATTACCGGGAGCTT |
| ABID_SBC_bot_349 | TACGATCA | /5Phos/GGCCCTTACGNNNATCANNNGATCGTCGGACTGTAGAACTCTGAACCCCTATAGTGAGTCGTATTACCGGGAGCTT |
| ABID_SBC_bot_350 | GCCGACTA | /5Phos/GGCCCTGCCGNNNACTANNNGATCGTCGGACTGTAGAACTCTGAACCCCTATAGTGAGTCGTATTACCGGGAGCTT |
| ABID_SBC_bot_351 | TACATGGT | /5Phos/GGCCCTTACANNNTGGTNNNGATCGTCGGACTGTAGAACTCTGAACCCCTATAGTGAGTCGTATTACCGGGAGCTT |
| ABID_SBC_bot_352 | CTTGTAGA | /5Phos/GGCCCTCTTGNNNTAGANNNGATCGTCGGACTGTAGAACTCTGAACCCCTATAGTGAGTCGTATTACCGGGAGCTT |
| ABID_SBC_bot_353 | AGCACTCT | /5Phos/GGCCCTAGCANNNCTCTNNNGATCGTCGGACTGTAGAACTCTGAACCCCTATAGTGAGTCGTATTACCGGGAGCTT |
| ABID_SBC_bot_354 | CGCACAAG | /5Phos/GGCCCTCGCANNNCAAGNNNGATCGTCGGACTGTAGAACTCTGAACCCCTATAGTGAGTCGTATTACCGGGAGCTT |
| ABID_SBC_bot_355 | GCGCATTA | /5Phos/GGCCCTGCGCNNNATTANNNGATCGTCGGACTGTAGAACTCTGAACCCCTATAGTGAGTCGTATTACCGGGAGCTT |
| ABID_SBC_bot_356 | ACGCAGCA | /5Phos/GGCCCTACGCNNNAGCANNNGATCGTCGGACTGTAGAACTCTGAACCCCTATAGTGAGTCGTATTACCGGGAGCTT |
| ABID_SBC_bot_357 | GCTCACTC | /5Phos/GGCCCTGCTCNNNACTCNNNGATCGTCGGACTGTAGAACTCTGAACCCCTATAGTGAGTCGTATTACCGGGAGCTT |
| ABID_SBC_bot_358 | TTATCGCG | /5Phos/GGCCCTTTATNNNCGCGNNNGATCGTCGGACTGTAGAACTCTGAACCCCTATAGTGAGTCGTATTACCGGGAGCTT |
| ABID_SBC_bot_359 | GAGAACAT | /5Phos/GGCCCTGAGANNNACATNNNGATCGTCGGACTGTAGAACTCTGAACCCCTATAGTGAGTCGTATTACCGGGAGCTT |
| ABID_SBC_bot_360 | ATCTGAGT | /5Phos/GGCCCTATCTNNNGAGTNNNGATCGTCGGACTGTAGAACTCTGAACCCCTATAGTGAGTCGTATTACCGGGAGCTT |
| ABID_SBC_bot_361 | CTTATGGC | /5Phos/GGCCCTCTTANNNTGGCNNNGATCGTCGGACTGTAGAACTCTGAACCCCTATAGTGAGTCGTATTACCGGGAGCTT |
| ABID_SBC_bot_362 | AGGTTACA | /5Phos/GGCCCTAGGTNNNTACANNNGATCGTCGGACTGTAGAACTCTGAACCCCTATAGTGAGTCGTATTACCGGGAGCTT |
| ABID_SBC_bot_363 | GTAGCTGT | /5Phos/GGCCCTGTAGNNNCTGTNNNGATCGTCGGACTGTAGAACTCTGAACCCCTATAGTGAGTCGTATTACCGGGAGCTT |
| ABID_SBC_bot_364 | GCTCCAAC | /5Phos/GGCCCTGCTCNNNCAACNNNGATCGTCGGACTGTAGAACTCTGAACCCCTATAGTGAGTCGTATTACCGGGAGCTT |
| ABID_SBC_bot_365 | GCCTATTC | /5Phos/GGCCCTGCCTNNNATTCNNNGATCGTCGGACTGTAGAACTCTGAACCCCTATAGTGAGTCGTATTACCGGGAGCTT |
| ABID_SBC_bot_366 | ATACACGA | /5Phos/GGCCCTATACNNNACGANNNGATCGTCGGACTGTAGAACTCTGAACCCCTATAGTGAGTCGTATTACCGGGAGCTT |
| ABID_SBC_bot_367 | TAACGCAT | /5Phos/GGCCCTTAACNNNGCATNNNGATCGTCGGACTGTAGAACTCTGAACCCCTATAGTGAGTCGTATTACCGGGAGCTT |
| ABID_SBC_bot_368 | TCGCCAGT | /5Phos/GGCCCTTCGCNNNCAGTNNNGATCGTCGGACTGTAGAACTCTGAACCCCTATAGTGAGTCGTATTACCGGGAGCTT |
| ABID_SBC_bot_369 | ATTGAGGC | /5Phos/GGCCCTATTGNNNAGGCNNNGATCGTCGGACTGTAGAACTCTGAACCCCTATAGTGAGTCGTATTACCGGGAGCTT |
| ABID_SBC_bot_370 | TCCGTATC | /5Phos/GGCCCTTCCGNNNTATCNNNGATCGTCGGACTGTAGAACTCTGAACCCCTATAGTGAGTCGTATTACCGGGAGCTT |
| ABID_SBC_bot_371 | ATACTGAG | /5Phos/GGCCCTATACNNNTGAGNNNGATCGTCGGACTGTAGAACTCTGAACCCCTATAGTGAGTCGTATTACCGGGAGCTT |
| ABID_SBC_bot_372 | CAATCTTG | /5Phos/GGCCCTCAATNNNCTTGNNNGATCGTCGGACTGTAGAACTCTGAACCCCTATAGTGAGTCGTATTACCGGGAGCTT |
| ABID_SBC_bot_373 | CACATCTA | /5Phos/GGCCCTCACANNNTCTANNNGATCGTCGGACTGTAGAACTCTGAACCCCTATAGTGAGTCGTATTACCGGGAGCTT |

| | | |
|---|---|---|
| ABID_SBC_bot_374 | ATCTCATG | /5Phos/GGCCCTATCTNNNCATGNNNGATCGTCGGACTGTAGAACTCTGAACCCCTATAGTGAGTCGTATTACCGGGAGCTT |
| ABID_SBC_bot_375 | CGATTAAG | /5Phos/GGCCCTCGATNNNTAAGNNNGATCGTCGGACTGTAGAACTCTGAACCCCTATAGTGAGTCGTATTACCGGGAGCTT |
| ABID_SBC_bot_376 | GGCATAAT | /5Phos/GGCCCTGGCANNNTAATNNNGATCGTCGGACTGTAGAACTCTGAACCCCTATAGTGAGTCGTATTACCGGGAGCTT |
| ABID_SBC_bot_377 | GAGCCACT | /5Phos/GGCCCTGAGCNNNCACTNNNGATCGTCGGACTGTAGAACTCTGAACCCCTATAGTGAGTCGTATTACCGGGAGCTT |
| ABID_SBC_bot_378 | GCACGTGT | /5Phos/GGCCCTGCACNNNGTGTNNNGATCGTCGGACTGTAGAACTCTGAACCCCTATAGTGAGTCGTATTACCGGGAGCTT |
| ABID_SBC_bot_379 | CACGTAAG | /5Phos/GGCCCTCACGNNNTAAGNNNGATCGTCGGACTGTAGAACTCTGAACCCCTATAGTGAGTCGTATTACCGGGAGCTT |
| ABID_SBC_bot_380 | CAACCACG | /5Phos/GGCCCTCAACNNNCACGNNNGATCGTCGGACTGTAGAACTCTGAACCCCTATAGTGAGTCGTATTACCGGGAGCTT |
| ABID_SBC_bot_381 | ACTATCCA | /5Phos/GGCCCTACTANNNTCCANNNGATCGTCGGACTGTAGAACTCTGAACCCCTATAGTGAGTCGTATTACCGGGAGCTT |
| ABID_SBC_bot_382 | TTGAGATC | /5Phos/GGCCCTTTGANNNGATCNNNGATCGTCGGACTGTAGAACTCTGAACCCCTATAGTGAGTCGTATTACCGGGAGCTT |
| ABID_SBC_bot_383 | AAGTCGGC | /5Phos/GGCCCTAAGTNNNCGGCNNNGATCGTCGGACTGTAGAACTCTGAACCCCTATAGTGAGTCGTATTACCGGGAGCTT |
| ABID_SBC_bot_384 | ATCGAGAT | /5Phos/GGCCCTATCGNNNAGATNNNGATCGTCGGACTGTAGAACTCTGAACCCCTATAGTGAGTCGTATTACCGGGAGCTT |
